# Paralogs and off-target sequences improve phylogenetic resolution in a densely-sampled study of the breadfruit genus (*Artocarpus*, Moraceae)

**DOI:** 10.1101/854232

**Authors:** Elliot M. Gardner, Matthew G. Johnson, Joan T. Pereira, Aida Shafreena Ahmad Puad, Deby Arifiani, Sahromi, Norman J. Wickett, Nyree J.C. Zerega

## Abstract

We present a 517-gene phylogenetic framework for the breadfruit genus *Artocarpus* (ca. 70 spp., Moraceae), making use of silica-dried leaves from recent fieldwork and herbarium specimens (some up to 106 years old) to achieve 96% taxon sampling. We explore issues relating to assembly, paralogous loci, partitions, and analysis method to reconstruct a phylogeny that is robust to variation in data and available tools. While codon partitioning did not result in any substantial topological differences, the inclusion of flanking non-coding sequence in analyses significantly increased the resolution of gene trees. We also found that increasing the size of datasets increased convergence between analysis methods but did not reduce gene tree conflict. We optimized the HybPiper targeted-enrichment sequence assembly pipeline for short sequences derived from degraded DNA extracted from museum specimens. While the subgenera of *Artocarpus* were monophyletic, revision is required at finer scales, particularly with respect to widespread species. We expect our results to provide a basis for further studies in *Artocarpus* and provide guidelines for future analyses of datasets based on target enrichment data, particularly those using sequences from both fresh and museum material, counseling careful attention to the potential of off-target sequences to improve resolution.

---

Reduced-representation methods such as target enrichment (HybSeq) have become important tools for phylogenetic studies, enabling high-throughput and cost-effective sequencing of hundreds of loci (Faircloth et al. 2012; Mandel et al. 2014; Weitemier et al. 2014). In this study, we employ HybSeq to investigate the breadfruit genus (*Artocarpus* J.R. Forst. & G. Forst., Moraceae), analyzing the utility of paralogs, partitioning, noncoding sequences, and herbarium specimens in reconstructing the most data-rich phylogeny of the genus to date.

HybSeq involves hybridizing a randomly-sheared sequencing library to bait sequences, typically exons from one or more taxa within or near the target clade. Researchers have employed HybSeq in studies ranging from deep phylogenetics (Prum et al. 2015; Liu et al. 2019) to within-species phylogeography (Villaverde et al. 2018). It is particularly useful for recovering sequences from museum specimens, because target enrichment is suitable for very small DNA fragments and can help overcome the presence of contaminating non-endogenous DNA (Staats et al. 2013; Buerki and Baker 2016; Hart et al. 2016; Brewer et al. 2019). However, making the most of HybSeq datasets, which can comprise hundreds of thousands of characters, requires careful attention to assembly and analysis methods, particularly for degraded DNA from museum specimens. This particularly true because divergent analysis methods can sometimes lead to divergent topologies, all with apparently high statistical support.

The mechanics of HybSeq frequently result in the recovery of non-targeted sequences such as paralogs similar to the target sequences (Hart et al. 2016; Johnson et al. 2016, 2019; Liu et al. 2019) and non-coding sequences flanking the target sequences (e.g. (Medina et al. 2019). Both were the case with HybSeq baits we previously developed for Moraceae phylogenetics (Gardner et al. 2016), many of which were represented as paralogous pairs in *Artocarpus* due to an ancient whole-genome duplication. In almost all cases they were diverged enough to sort and analyze separately (Johnson et al. 2016). The same targets also typically recovered a several-hundred bp “splash zone” of flanking non-coding sequences (Johnson et al., 2016). The impact of off-target by-catch on phylogenetic reconstruction remains unclear but has the potential to greatly increase the number of phylogenetically informative genes. However, analysis of mixed coding and non-coding sequences can make it difficult to ensure that exons are aligned in frame, particularly when frameshifts are present (Ranwez 2011), hampering partitioning of datasets by codon position. How these issues impact phylogenetic reconstruction remains unclear (Xi et al. 2012; Lanfear et al. 2014).

It is by now well understood that high bootstrap values obtained by concatenating all loci into a supermatrix should not be overinterpreted because near-perfect bootstrap support can mask substantial discordance among gene histories due to incomplete lineage sorting (Kubatko and Degnan 2007; Degnan and Rosenberg 2009; Sayyari and Mirarab 2016). Although there is an increased availability of efficient methods based on the multi-species coalescent model, clear results can be obscured if the underlying gene trees are uninformative (Smith et al. 2015; Sayyari et al. 2017). A major advantage of HybSeq over methods with short, anonymous loci, or large amounts of missing data, is that loci obtained via HybSeq are both long enough to generate single-gene phylogenies and subject to few enough missing taxa per locus for those single-gene phylogenies to be informative. These and other issues are explored below to develop a robust phylogenomic framework that will guide future work in *Artocarpus* and serve as a model for work in other systems.

## Study system

*Artocarpus* (Fig. 1) contains approximately 70 species of trees with a center of diversity in Borneo and a native range that extends from India to the Solomon Islands (Williams et al. 2017). The genus is best known for important but underutilized crops such as breadfruit (*A. altilis* (Parkinson) Fosberg) and jackfruit (*A. heterophyllus* Lam.) (Zerega et al. 2010, 2015; Wang et al. 2018; Witherup et al. 2019).

**Figure 1.**
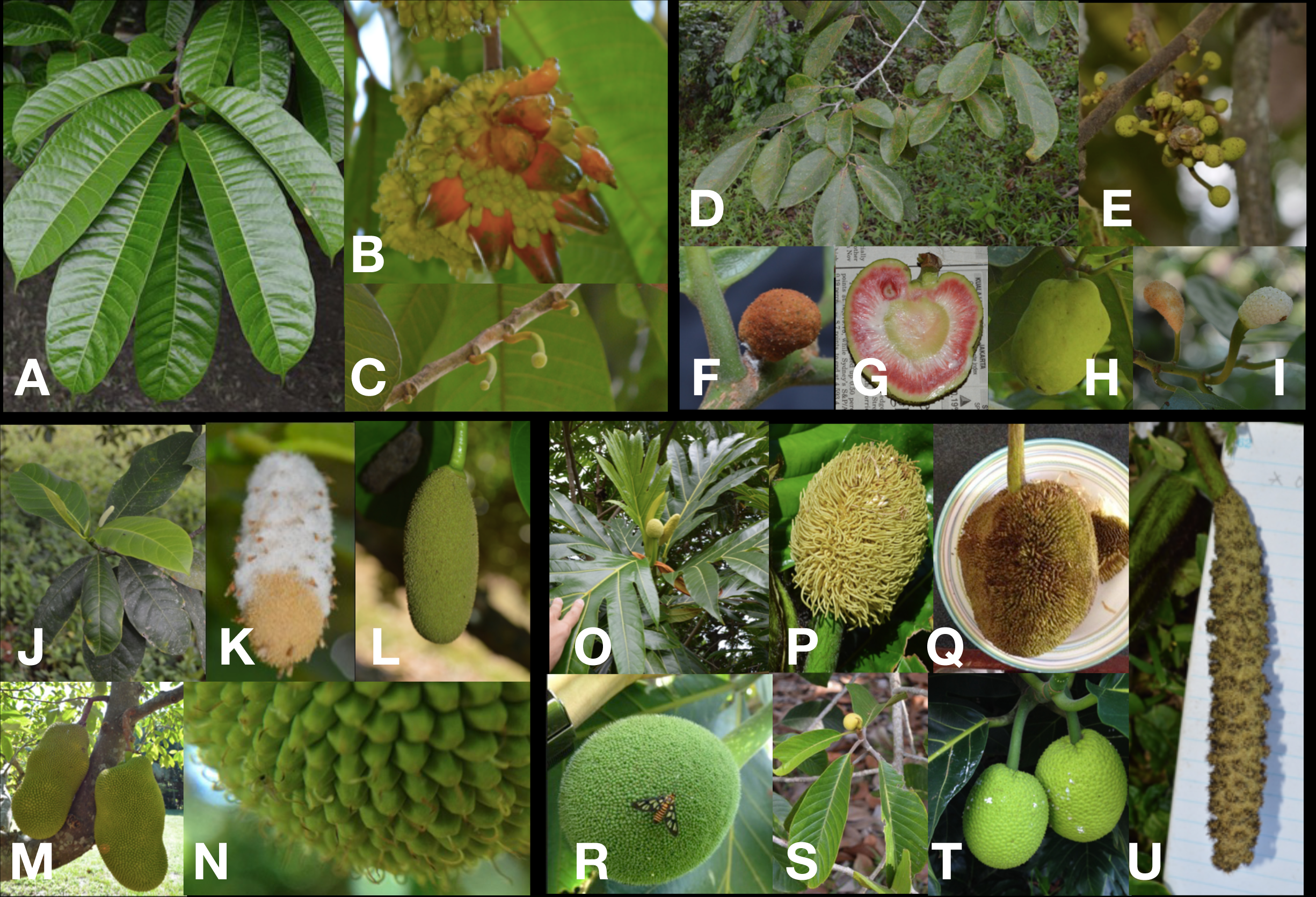
Diversity of *Artocarpus*. **Subg**. *Prainea*— (A) leaves, (B) syncarp, and (C) immature inflorescences of *A. limpato*. **Subg**. *Pseudojaca —* (D) leaves and (E) staminate inflorescences of *A. fretessii;* (F) carpellate inflorescence of *A. nitidus ssp. borneensis;* (G) syncarp of *A. primackii;* (H) syncarp of *A. nitidus ssp. lingnanensis;* and (I) staminate (left) and carpellate (right) inflorescences of *A. hypargyreus*. **Subg**. *Cauliflori*— (J) leaves of *A. integer*; (K–L) staminate inflorescences, (M) syncarps, and (N) carpellate inflorescence of *A. heterophyllus*. **Subg**. *Artocarpus*— (O) leaves and inflorescences of *A. altilis*; (P) carpellate inflorescence of *A. tamaran;* (Q) syncarp and (R) carpellate inflorescence of *A. odoratissimus*; (S) leaves and staminate inflorescence of *A. rigidus*; (T) syncarps of *A. altilis*; and (U) staminate inflorescence of *A. tamaran*.

*Artocarpus* is monecious, with spicate to globose staminate (“male”) inflorescences composed of tiny flowers bearing one stamen each. Pistillate (“female”) inflorescences are composed of tightly packed tiny flowers, and in most cases adjacent flowers are at least partially fused together. Pistillate inflorescences develop into tightly packed accessory fruits composed mainly of fleshy floral tissue, ranging from a few centimeters in diameter in some species to over half a meter long in jackfruit. The tribe Artocarpeae also includes two smaller Neotropical genera, *Batocarpus* H. Karst. (3 spp.) and *Clarisia* Ruiz & Pav. (3 spp.); these always have spicate staminate inflorescences; pistillate flowers may be solitary or condensed into globose heads, but adjacent flowers are never fused.

The most recent complete revision of *Artocarpus* (Jarrett 1959, 1960) recognized two subgenera, *Artocarpus* and *Pseudojaca* Tréc., distinguished by phyllotaxy (leaf arrangement), and the degree of fusion between adjacent pistillate flowers. Since then, several new species have been described (Jarrett, 1975; Zhengyi and Xiushi, 1989; Kochummen, 1998; Berg, 2005; Gardner et al. in prep.; Gardner and Zerega in prep.). Berg et al. (2006) revised the Malesian species for the *Flora Malesiana*, in a few cases combining several taxa into a broadly-circumscribed single species. Examples include *A. altilis* (encompassing *A. altilis, A. camansi, A. mariannensis* Tréc., *A. horridus* F.M. Jarrett, *A. blancoi* Merr., *A. pinnatisectus* Merr., and *A. multifidus* F.M. Jarrett) and *A. lacucha* Buch.-Ham. (encompassing *A. lacucha, A. dadah* Miq., *A. fretessii* Teijsm. & Binnend., *A. ovatus* Blanco, and *A. vrieseanus* var. *refractus* Becc. (F.M. Jarrett)). For clarity, we generally follow Jarrett’s nomenclature. The most recent circumscription of *Artocarpus* recognized four subgenera (Fig. 1) and was based on two gene regions and approximately 50% of taxa (Zerega et al. 2010). The subgenera are distinguished by phyllotaxy, the degree of fusion between adjacent pistillate flowers, and the position of inflorescences on the tree: axillary (from a leaf-joint on a small twig) or cauliflorous (from the trunk or a main brank). A well-sampled phylogenetic framework for *Artocarpus* is necessary to inform future taxonomic revision and to clarify relationships within this important genus, in particular the relationships between crop species and their wild relatives, whose conservation is a priority (Castañeda-Álvarez et al. 2016).

In this study, we used near-complete (80/83) taxon sampling (at the subspecies level or above) in *Artocarpus* to reconstruct the most data-rich phylogeny to date for *Artocarpus*, taking into account the impact of paralogs, codon partitions, noncoding sequences, and analysis method (species tree versus concatenated supermatrix) on phylogenetic reconstruction in order to develop a truly robust phylogenetic hypothesis. We also used this data set to improve the target capture assembly pipeline HybPiper, which is now optimized for accurately scaffolding small disconnected contigs resulting from degraded DNA. The objectives of the study were to (1) use broad sampling from silica dried material and herbarium specimens over 100 years old to achieve near complete taxon sampling for *Artocarpus*; (2) test the monophyly of the current taxonomic divisions within *Artocarpus* to provide a phylogenetic framework for future studies on the taxonomy, conservation, and ecology of the genus; and (3) examine the impact of paralogs, partitions, and analysis method on phylogenetic reconstruction.

## Materials and methods

A summary of our methods follows. Further details, including protocol modifications for herbarium material and software parameters, can be found in Appendix 1.

### Data accessibility

Raw reads have been deposited in GenBank (BioProject no. PRJNA322184), and alignments and trees have been deposited in the Dryad Data Repository (accession no. TBA). HybPiper and related scripts used in this study are available at https://github.com/mossmatters/HybPiper and https://github.com/mossmatters/phyloscripts.

### Taxon sampling

We sampled all *Artocarpus* taxa at the subspecies level or above (Jarrett 1959, 1960; Berg et al., 2006; Kochummen, 1998; Wu and Zhang, 1989) and nine taxa of questionable affinities, replicating sampling across geographic or morphological ranges when possible, for a total of 167 ingroup samples belonging to 83 names. Outgroups included one species per genus in the Neotropical Artocarpeae and the sister tribe Moreae. Samples came from field collections preserved in silica gel, botanic gardens, and herbaria (up to 106 years old), totaling 179 samples (Table S1).

### Sample preparation and sequencing

DNA extracted from ca. 0.5 cm^2^ of leaf tissue was quantified on a Qubit fluorometer (Invitrogen, Life Technologies, CA, USA) and assessed on an agarose gel or a High-Sensitivity DNA Assay on a BioAnalyzer 2100 (Agilent). Samples with an average fragment size of >500bp were sonicated to ca. 550bp using a Covaris M220 (Covaris, Wobum, Massachussetts, USA), and libraries were prepared with the Illumina TruSeq Nano HT DNA Library Preparation Kit (Illumina, San Diego, California, USA) or the KAPA Hyper Prep DNA Library Kit (KAPA, Cape Town, South Africa), using 200ng of input DNA when possible. Pools of 6–24 libraries were enriched for 333 phylogenetic markers (Gardner et al., 2016) with a MYbaits kit (MYcroarray, Ann Arbor, Michigan, USA) and reamplified with 14 PCR cycles. Sequencing took place on an Illumina MiSeq (2×300bp, v3) in runs of 30–99 samples.

### Sequence quality control and analyses

In addition to samples prepared for this study, our analyses included reads from all *Artocarpus* samples from Johnson et al. (2016) as well as the original 333 orthologues from *Morus notabilis* described by Gardner et al. (2016). Demultiplexed and adapter-trimmed reads were quality trimmed using Trimmomatic 0.39 (Bolger et al., 2014) and assembled with HybPiper 1.2 (Johnson et al. 2016), which represents a compromise between read mapping and *de novo* assembly and combines local *de novo* assemblies with scaffolding based on a reference coding sequence (Johnson et al. 2016, 2019). We used the Moraceae reference from Kates et al. (2018), supplemented with additional *Artocarpus* taxa representing all subgenera. This reference contained the original orthologs from Gardner et al. (2016) in addition to the paralogs identified in *Artocarpus* by Johnson et al. (2016); paralogs were treated as separate loci and were used for ingroup assemblies only.

HybPiper output used here includes (1) the predicted coding sequence for each target gene (“exon”) and (2) for the ingroup only, the entire contig assembled for each gene, including non-coding sequences (“supercontig”). To reduce bias from sequencing errors, assemblies were masked to remove positions covered by fewer than two reads (Li and Durbin 2009; Li et al. 2009; Quinlan and Hall 2010; Broad Institute 2016). To reduce noise associated with high amounts of missing data, within each gene alignment we removed samples whose *exon* sequences were less than 150 bp or 20% of the average sequence length for that gene, and samples with fewer than 100 genes after filtering were excluded entirely.

For “exon” sequences, we created in-frame alignments using MACSE 1.02 (Ranwez 2011). For supercontig output, we used MAFFT 7.211 for alignment (--maxiter 1000) (Katoh and Standley 2013). We trimmed alignments to remove columns with >75% gaps using Trimal (Capella-Gutiérrez et al. 2009). Finally, we built gene trees from the exon alignments using FastTree (Price et al. 2009) and visually inspected them for long internal branches to identify alignments containing obvious improperly sorted paralogous sequences; alignments were visually inspected with AliView (Larsson 2014), and 12 genes were discarded, resulting in a final set of 517 genes, including all of the original 333 genes plus 184 paralogs.

We used the trimmed alignments to create three datasets:

1. *CDS:* exon alignments (in-frame), not partitioned by codon position;
2. *Partitioned CDS*: exon alignments (in-frame), partitioned by codon position; and
3. *CDS+noncoding*: supercontig alignments, not partitioned within genes

Each dataset was each analyzed with and without paralogs, using the following two methods, for a total of 12 analyses: (A) *Concatenated supermatrix:* all sequences concatenated and partitioned by gene (or by gene and codon, depending on the dataset) and analyzed using RAxML 10 (Stamatakis, 2006) under the GTRCAT model with 200 rapid bootstrap replicates; (B) *Species tree:* each gene alignment analyzed using RAxML 10 under GTRCAT with 200 rapid bootstrap replicates. Nodes with <33% support were collapsed using SumTrees 4.3.0 (Sukumaran and Holder 2010), and the resulting trees were used to estimate a species tree with ASTRAL-III 5.5.6 (Mirarab and Warnow, 2015), calculating bootstrap (-r, 160) and quartet support (-t 1) for each node (Mirarab and Warnow 2015; Zhang et al. 2017). We used SumTrees to calculate the proportion of gene trees supporting each split; however, quartet support is less sensitive to occasional out-of-place taxa than raw gene-tree support. The *exon* analyses were repeated using the GTRGAMMA model in RAxML. Attempts to produce a partitioned “supercontig” dataset were not successful, because aligning noncoding sequences separately produced unreliable alignments (Appendix 2). Supermatrix analyses took place on the CIPRES Science Gateway (Miller et al. 2010), and all others took place on a cluster at the Chicago Botanic Garden. Most processes were run in parallel using GNU Parallel (Tange 2018).

To summarize the overall bootstrap support of each tree with a single statistic, we calculated “percent resolution,” which represents the proportion of bipartitions with >50% bootstrap support (Kates et al. 2018). We visualized trees using FigTree 1.4.3 (Rambaut 2016) and analyzed and compared trees in R 3.5.1 (R Core Development Team, 2008) using ape 5.2 (Paradis et al. 2004), phytools 0.6-60 (Revell 2012), Lattice 0.20-38 (Sarkar 2008) and Phangorn 2.4.0 (Schliep 2011). Analyses included analyses of differences in topologies and pairwise Robison-Foulds (RF) distances (the sum of disagreeing bipartitions) for all trees.

## Results

### Sequencing and assembly

Of the 179 sequenced accessions, 164 resulted in assemblies with at least 25 genes (Fig. S1, Table S1), including all attempted taxa except for *A. nigrifolius* C.Y. Wu and *A. nanchuanensis*, C.Y. Wu, two species closely allied to *A. hypargyreus* Hance, which was assembled, and *A. scandens* Miq. sensu Jarrett, considered conspecific with *A. frutescens* Becc. by Berg et al. (2006), which was also assembled. Less successful samples generally had few reads and may have been out-competed by other samples during hybridization, reamplification, or both. Fewer reads were also associated with shorter assembled sequences (Fig. S1). Only samples with at least 100 genes were used for phylogenetic analyses, resulting in the loss of five additional samples and one taxon, *A. reticulatus* Miq. Adding the *Morus notabilis* sequences resulted in a final data set of 160 samples representing 80 out of 83 named *Artocarpus* taxa at the subspecies/variety level or above (96%) and nine taxa of uncertain affinity.

Overall, samples collected more recently showed improved sequencing results (Fig. S2), primarily because the majority of samples collected since 2000 were dried on silica gel. Whether a sample was dried on silica gel was significantly associated with increased gene length as a percentage of average length (*R*^*2*^ = 0.33, *P* < 0.0001) and to a lesser extent with the total number of genes recovered (*R*^*2*^ = 0.17, *P* < 0.0001). All 16 unsuccessful (<25 genes) assemblies were taken from herbarium sheets (collected between 1917 and 1997), rather than silica-dried material. Among 67 successfully-assembled herbarium samples, younger age was associated with increased gene length, although the model was a poor fit (*R*^*2*^ = 0.06, *P* = 0.02728), but not with an increase in the number of genes recovered (*P* = 0.2833) (Fig. 3). By the same token, we observed a decrease in average DNA fragment size in older samples (Fig. S3). Lowering the maximum assembly k-mer values for herbarium samples with under 400 genes increased recovery by an average of 20 genes.

**Figure 2.**
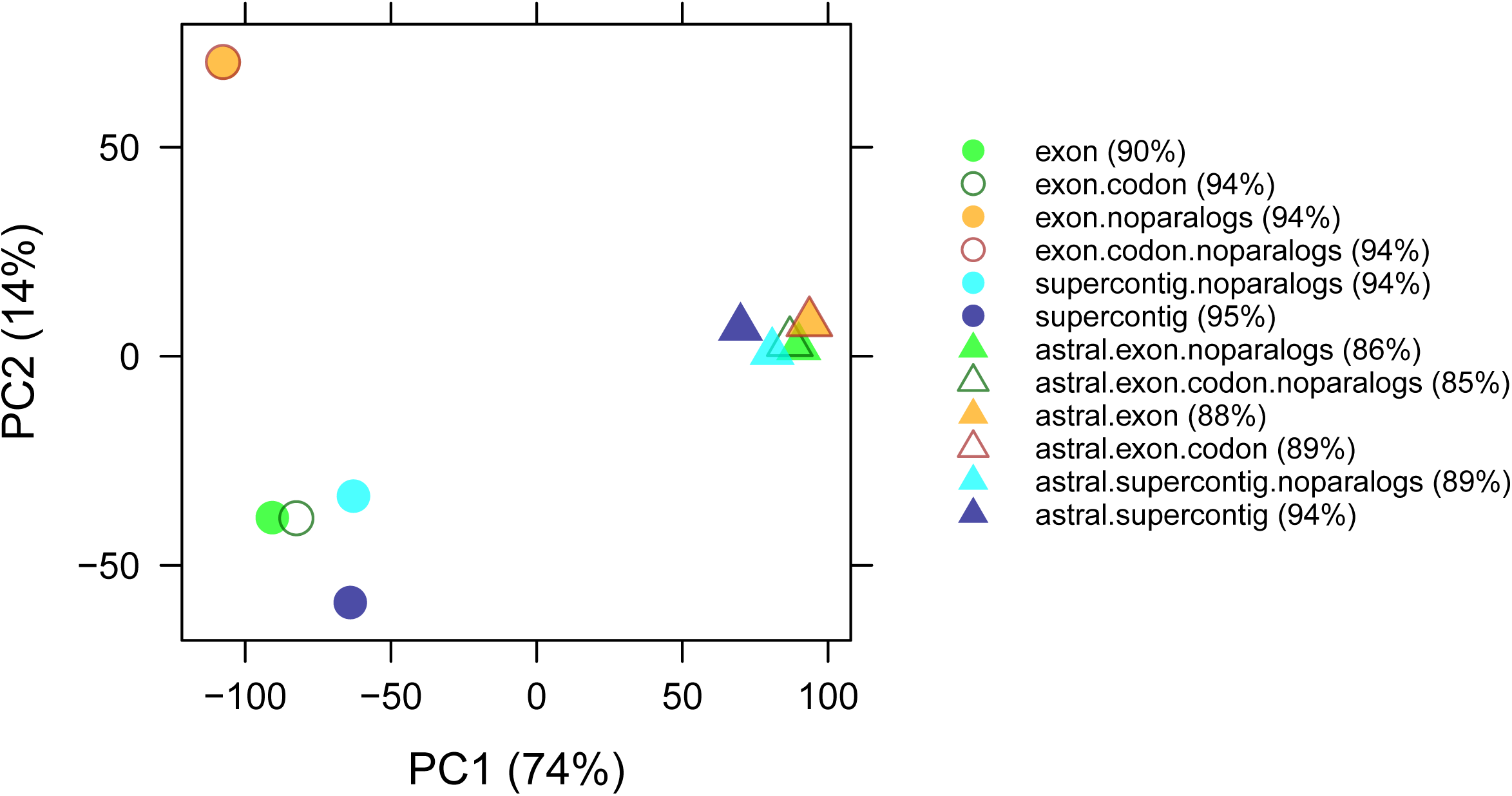
PCA of Robinson-Foulds (RF) distances between all 12 main analyses.

**Figure 3.**
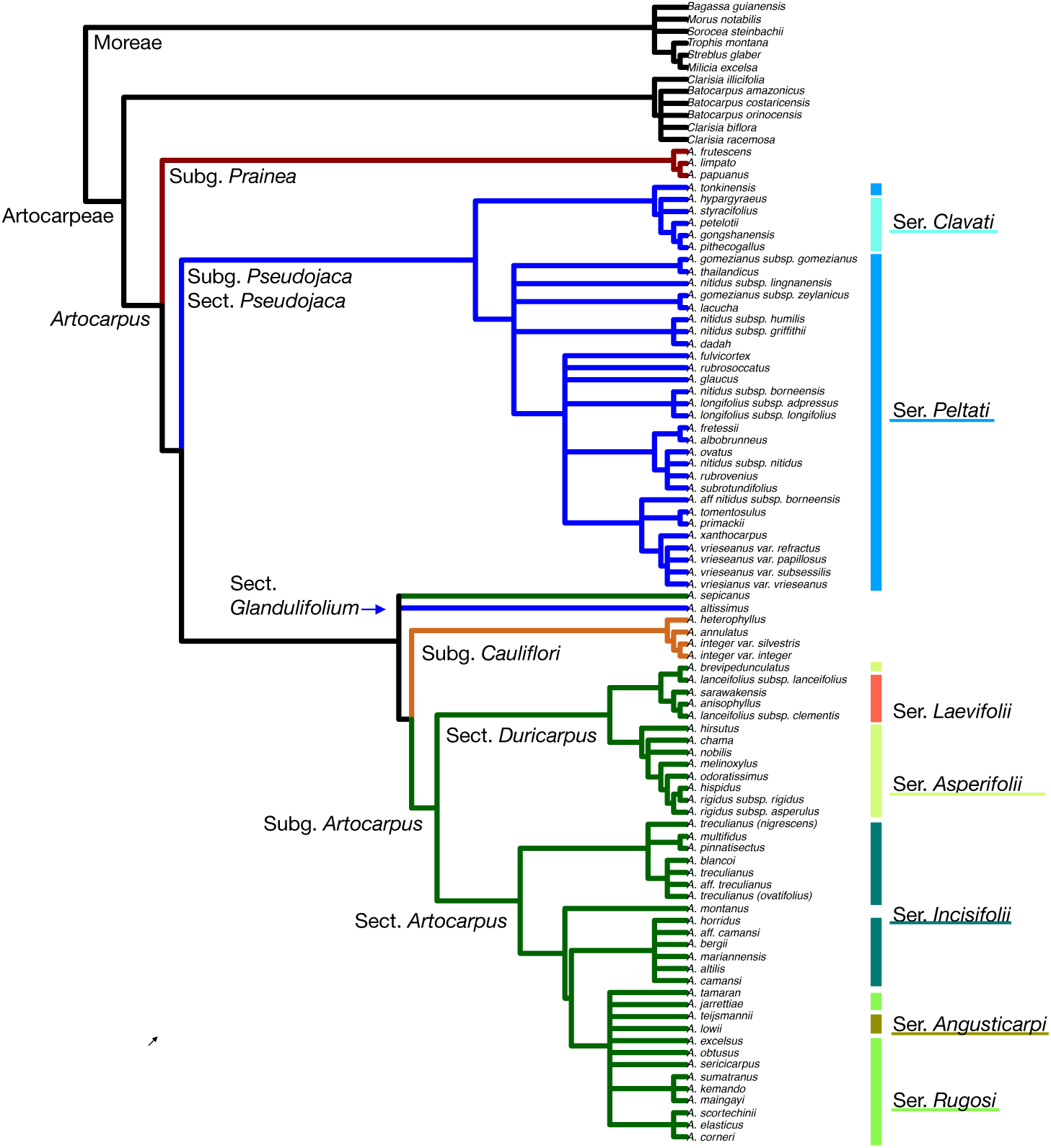
Strict consensus of all 12 main-analysis trees (excluding only those analyses in which exons and introns were aligned separately, for reasons discussed in the text). Colored boxes reflect Jarrett’s (1959, 1960) taxonomic divisions, as modified by Zerega et al. (2010). Recently-described taxa that were split from older taxa recognized by Jarrett are classified according to Jarrett’s species concepts. Labels to the right of the tree denote major non-monophyletic taxonomic divisions.

Gene recovery was high; the average sample (of the 160 passing the final filter) had sequences for 448/517 genes (87%). In the final filtered dataset of 333 genes, the average gene had exon sequences for 151/160 samples (94%, range 57–160, median 154) and non-coding sequences for 130 (81%, range 49–151, median 131). For the 184 paralogs, the average gene in the final filtered dataset had exon sequences for 117/160 samples (73%, range 32–148, median 126) and intron sequences for 101 (63%, range 30–132, median 110) (Table S2). The supermatrix of trimmed *exon* alignments for the primary 333 genes contained 407,310 characters; and the full set of 517 *exon* alignments, including 184 paralogs, contained 569,796 characters. The supermatrix of 333 trimmed *supercontig* alignments contained 813,504 characters, and the full set of 517 genes contained 1,181,279 characters. The full set of *exon* alignments had 21% gaps or undetermined characters, while the full set of *supercontig* alignments was 36.87% gaps or undetermined characters.

### Phylogenetic disagreement

A strict consensus of the 12 species trees under GTRCAT (henceforth: “main analysis”) had 100/159 (63%) nodes resolved (mean RF distance 53), revealing agreement in backbone relationships between the major subgenera but substantial disagreement at shallower nodes (Fig. 3). The six ASTRAL phylogenies differed little from one another, whereas supermatrix analyses had somewhat greater divergence (Fig. 2, Table S3).

#### Partitions and model selection

In *exon* datasets, partitioning by codon position (Figs. 2, S4–S7) had little impact on final topology, with only a single within-species rearrangement (RF 4), but in the ASTRAL analysis, partitioning by codon position caused *A. sepicanus* + *A. altissimus* to form a grade rather than a clade, as in all other analyses (RF 12). The choice of model (GTRCAT vs GTRGAMMA) also produced only minor changes (Figs. S8–S12).

#### Paralogs

Addition of paralogs led to slightly more disagreement (Figs. 2, S13–S16, Table S3). In the *exon* dataset, changes to the positions of *A. nitidus* ssp. *lingnanensis* (Merr.) F.M. Jarrett and *A. gomezianus* Wall. ex Tréc. affected the backbone of ser. *Peltati* F.M. Jarrett, subg. *Pseudojaca*, in the supermatrix analysis (RF 58); in the ASTRAL analysis, there were fewer rearrangements, mainly in the same clade (RF 20). However, disagreement was reduced when noncoding sequences were included (supermatrix RF 22; ASTRAL RF 8).

#### Introns

Inclusion of non-coding sequences (Figs. 2, S17–S20, Table S3) led to similar amounts of disagreement, with rearrangements at the series level in subg. *Pseudojaca* and subg. *Artocarpus*. Disagreement was greater in the supermatrix analyses (no paralogs) (RF 62) than in ASTRAL analyses (RF 36). Addition of paralogs reduced disagreement in both cases (supermatrix RF 38; ASTRAL RF 26).

#### Analysis

The greatest differences among the 12 trees were between ASTRAL and supermatrix trees (Figs. 2, 4, S21–23, Table S3), with a mean RF distance between the six supermatrix trees and six ASTRAL trees of 78. Again, addition of noncoding regions or paralogs reduced disagreement between supermatrix and ASTRAL analyses; average RF distance for exons/noparalogs was 85, exons+paralogs 77, supercontig/noparalogs 71, and supercontigs+paralogs 70. Agreement was higher among ASTRAL trees (mean RF 21, 138/159 nodes in agreement) than among supermatrix trees (mean RF 48, 116/159 nodes in agreement). Differences (RF 66) between ASTRAL and supermatrix analyses for the full dataset (supercontigs+paralogs for all genes) at the species level can be ascribed to mostly minor repositionings within subclades involving two outgroup taxa (*Bagassa guianensis* Aubl. and *Batocarpus orinoceros* H. Karst.) and 14 ingroup taxa (Fig. 4).

**Figure 4.**
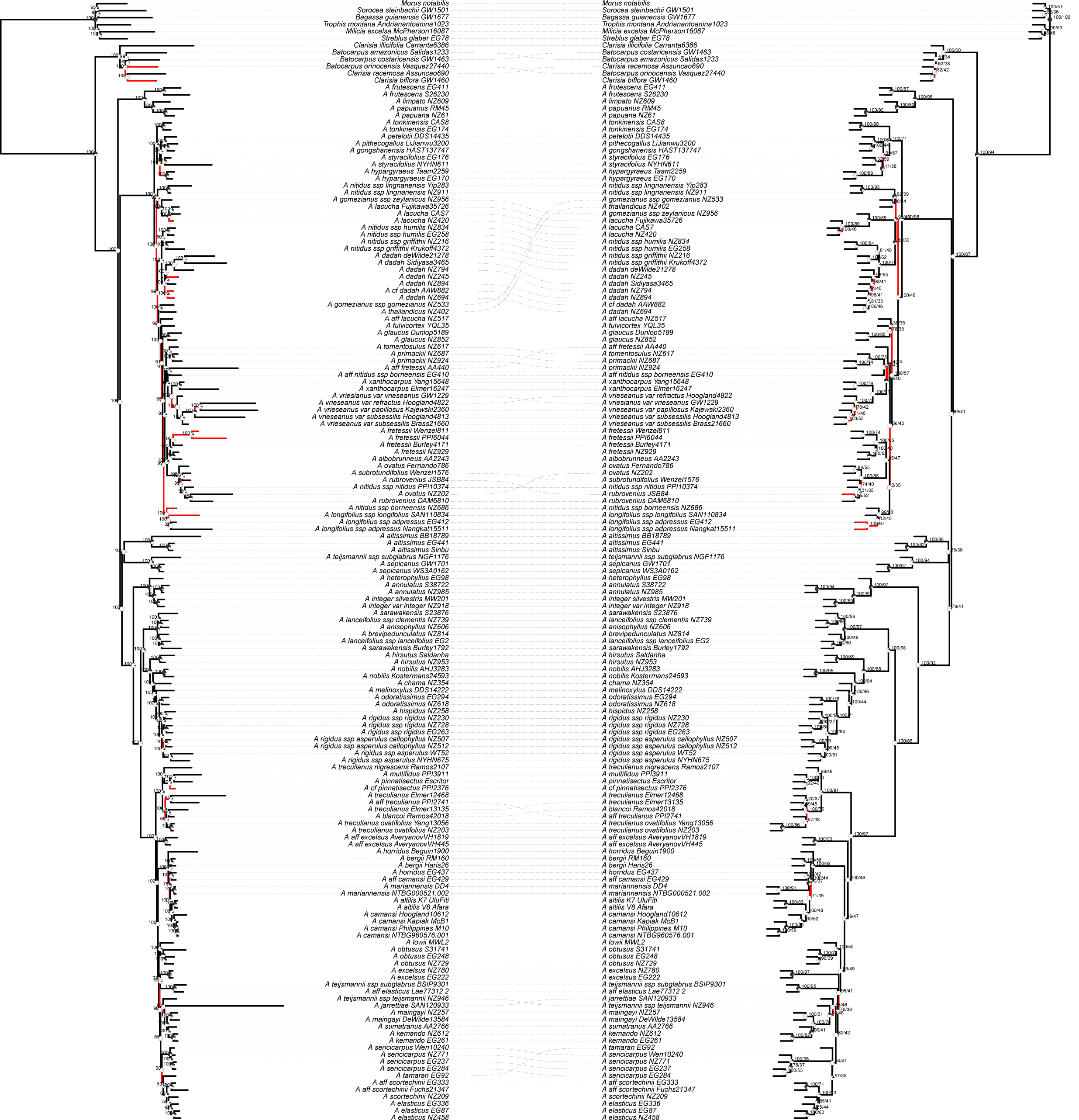
Comparison between the full-dataset (supercontigs for all genes) supermatrix and ASTRAL trees, with disagreeing branches in red, showing moderate disagreement at shallow phylogenetic depths but complete agreement at deeper nodes. Left: maximum-likelihood tree based on all supercontigs, partitioned by gene, including all paralogs; all branch lengths are proportional to mean substitutions per site. Right: ASTRAL tree based on all supercontigs; internal branch lengths are proportional to coalescent units; terminal branch lengths were arbitrarily assigned to improve visualization. Pie charts at nodes represent the proportion of gene trees supporting each split, and numbers represent bootstrap support

### Phylogenetic resolution

Percent resolution based on bootstrap values was 90–95% for all supermatrix trees in the main analysis and did not differ materially between analyses. Among ASTRAL trees, resolution was between 84% and 97% for all analyses. By slight margins, the best-resolved trees for both supermatrix and ASTRAL analyses were those based on the largest dataset (Table S3). Resolution based on quartet support for ASTRAL trees was between 57% and 60%, reflecting substantial gene tree discordance (Table S4, Fig. 4). For resolution measured by gene tree support (percentage of nodes supported by at least half of the 517 gene trees), scores ranged from 17–24%. In general, analyses including paralogs had reduced gene tree support, and trees based on supercontigs with no paralogs had the highest scores (24% for both ASTRAL and supermatrix).

More detailed analysis of differences between species trees based on the *exon* and *supercontig* datasets revealed that even if final species trees had similar resolution, *supercontig* trees were based on more information because the gene trees were significantly more informative. Inclusion of non-coding sequences significantly the number of splits with over 30% support (mean of +8). Because nodes under 30% were collapsed for species tree estimation, the species tree in the *supercontig* dataset was based on 9% more splits across the 517 gene trees (total of 51,307) than the species tree in the *exon* dataset (total 47,067). These patterns persisted in no-paralog datasets (mean increase in nodes over 30%: +9; overall difference in splits for 333 collapsed trees: 36,373 vs. 33,404 or 9%). Because addition of non-coding sequences also increased agreement between supermatrix and ASTRAL analyses (see above), this suggests at least some disagreement between supermatrix and species-tree analyses arises not only from incomplete lineage sorting but also from lack of resolution at the gene tree level, something that has also been observed at deeper phylogenetic scales (Pease et al. 2018). While we used low bootstrap support as an indicator of poor resolution in single-locus gene trees, we again caution against over-interpreting high bootstrap values, particularly in multi-locus trees.

### Phylogenetic relationships

The genus *Artocarpus* was monophyletic in all 12 main analyses, as were subgenera *Cauliflori* F.M. Jarrett and *Prainea* (King) Zerega, Supardi, & Motley (Table 1). Subgenus *Artocarpus* was monophyletic excluding *A. sepicanus* Diels, and subgenus *Pseudojaca* was monophyletic excluding *A. altissimus* J.J. Smith. In 10/12 analyses, *Artocarpus sepicanus* and *A. altissimus* formed a clade sister to subgenera *Cauliflori* and *Artocarpus*; however, in codon-partitioned ASTRAL analyses, they formed a grade in the same position (Figs. S5, S7). The backbone phylogeny was otherwise identical in all 12 trees: subgenus *Prainea* was sister to all other *Artocarpus*, which comprised a grade in this order: subgenus *Pseudojaca, A. sepicanus* + *A. altissimus* (usually), followed by subgenus *Cauliflori* + subgenus *Artocarpus*. Apart from the monophyly of the genus, which was supported by 61% of gene trees in the complete dataset (supercontig, all genes), subgeneric relationships had much less support at the gene tree level. The position of subg. *Prainea* was supported by 28% of gene trees; subg. *Pseudojaca* by 7%, and subgenera *Artocarpus/Cauliflori* by 4%. Quartet support was substantially higher (Fig. 4).

**Table 1.**
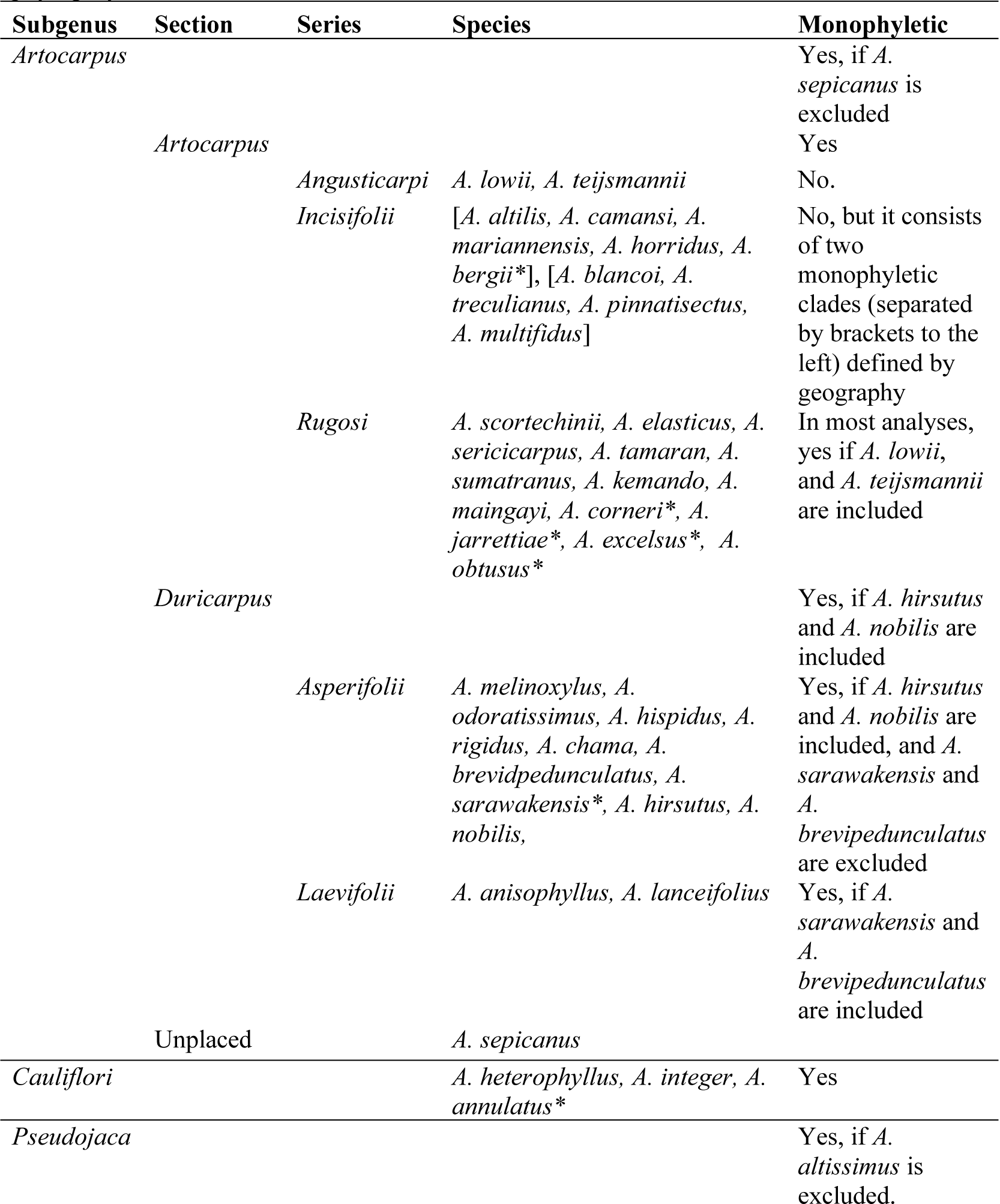
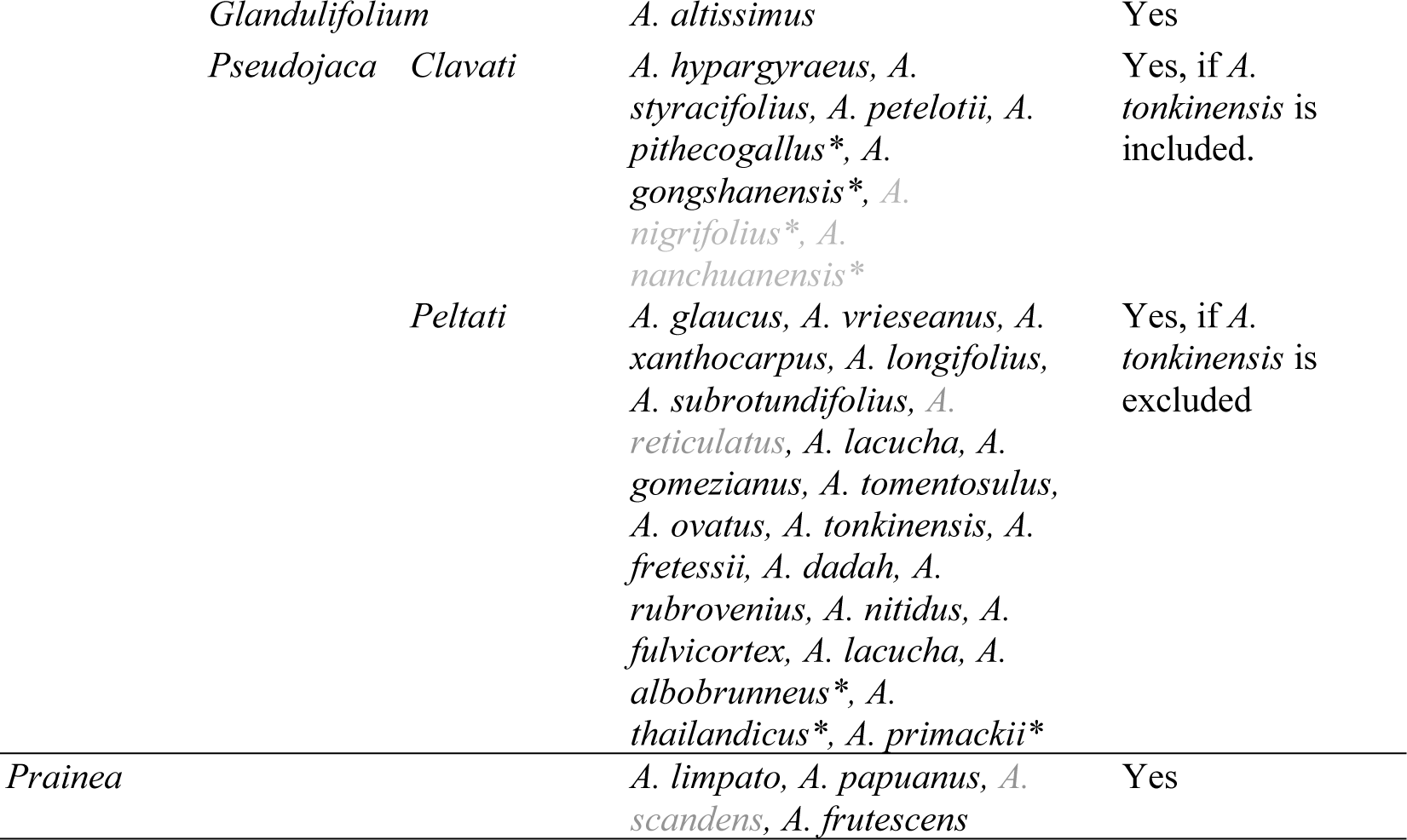
A summary of *Artocarpus* taxonomy following Zerega et al. (2010) at the subgeneric level, and Jarrett (1959–1960) at the section and series level. Species marked with an asterisk (*) were described after Jarrett’s revision; we have placed them into taxonomic divisions based on the phylogeny presented in this study. Species in gray text were not included in the phylogeny.

Within subgenus *Artocarpus*, both of Jarrett’s sections were monophyletic (excepting *A. sepicanus, A, hirsutus*, and *A. nobilis*, which she considered anomalous and did not place in sections), but none of the five series were monophyletic. However, series *Rugosi* F.M. Jarrett, characterized by rugose staminate inflorescences, was “nearly monophyletic” in most analyses, requiring only the inclusion of three non-rugose species (*A. teijsmannii* Miq., *A. lowii* King, and *A. excelsus* F.M. Jarrett). Members of series *Incisifolii* F.M. Jarrett, characterized by incised adult leaves, formed two non-sister monophyletic clades, one in the Philippines and one ranging from Indonesia to Oceania. Within subgenus *Pseudojaca*, section *Pseudojaca* was monophyletic (excluding *A. altissimus*), as was series *Clavati* F.M. Jarrett—characterized by clavate interfloral bracts. Series *Peltati* F.M. Jarrett—characterized by peltate interfloral bracts—would be monophyletic if *A. tonkinensis* A. Chev. were excluded, the latter species being sister to series *Clavati* in all main analyses.

Most species (for which we included at least two samples) were monophyletic, but several were not monophyletic in any analysis, including *A. treculianus* Elmer, *A. sarawakensis* F.M. Jarrett, *A. lanceifolius* Roxb., *A. rigidus* Blume, *A. teijsmanni*, and *A. nitidus* Tréc. The type of *A. teijsmannii* ssp. *subglabrus* C.C. Berg was sister to *A. sepicanus* in all analyses, while *ssp. teijsmannii* and other accessions of *ssp. subglabrus* were elsewhere within subgenus *Artocarpus*.

The neotropical Artocarpeae formed a clade sister to *Artocarpus* in all 12 trees. While *Batocarpus* was monophyletic in all supermatrix analyses, neither *Batocarpus* nor *Clarisia* was monophyletic in any ASTRAL tree.

## Discussion

### Taxon sampling

Although other studies have successfully applied target enrichment to recover sequences from herbarium and museum material (Guschanski et al. 2013; Hart et al. 2016), to our knowledge, this is among the first to use herbarium collections to achieve near-complete taxon sampling in a tropical plant genus of this size (ca. 70 spp.). The ability to successfully sequence herbarium material was indispensable for this study. For 34 of 90 (38%) ingroup taxa in the final analyses (including subspecies and the nine individuals of uncertain affinities), we did not have access to any fresh or silica-dried material and relied exclusively on herbarium specimens. In some cases, the only readily available samples were approximately 100 years old (e.g. *A. treculianus* sensu stricto (coll. 1910–1911: 369–370 genes recovered after filtering), *A. nigrescens* Elmer (coll. 1919: 431 genes), and *A. pinnatisectus* (type coll. 1913: 425 genes)). Although old samples had a lower success rate than silica-dried material, and sample degradation contributed to shorter assembled contigs, age alone was not significantly associated with recovery of fewer loci. Instead, the number of reads obtained was the most important factor in determining the number of loci recovered (Fig. S2). We hope these results encourage others to aim for complete taxon sampling with minimally-destructive sampling from natural history collections when newly-collected material is not available, so long as identifications can be confirmed by taxonomic experts. We note that during the course of this study, we corrected a substantial number of misidentifications.

While fieldwork remains among the most important aspects for systematic biology studies, phylogenetic reconstruction can benefit dramatically from incorporation of DNA from museum specimens. In this study, we successfully sequenced several DNA extractions from museum specimens that had been unusable for Sanger sequencing because PCR amplification failed (presumably due to small fragment size) (Zerega et al. 2010; Williams et al., 2017). The ability to achieve near-complete taxon sampling from museum material will open new opportunities for phylogeny-based analyses of clades with species that are difficult to collect, rare, or extinct, but present in herbarium collections. Our results suggest that near-complete taxon sampling can improve consistency between analyses, resulting in more reliable phylogenies. A previous study (Kates et al. 2018) using a smaller dataset of 22 *Artocarpus* species, found substantial disagreement between analyses in the backbone phylogeny of *Artocarpus*. Here, all 12 main analyses recovered almost identical backbones, disagreeing occasionally regarding positions of *A. altissimus* and *A. sepicanus*. Others have likewise found that missing taxa can substantially impact phylogenetic reconstructions (de la Torre-Bárcena et al. 2009). Robust taxon sampling also has serious implications for biodiversity conservation. *Artocarpus treculianus* is listed as Vulnerable by IUCN (World Conservation Monitoring Centre 1998). Due to availability of sequences from century-old herbarium sheets, we now know that this species is not monophyletic and that two obsolete taxa (*A. ovatifolius* Merr. and *A. nigrescens*) that Jarrett sunk into *A. treculianus* (Jarrett 1959) should probably be reinstated. Splitting a Vulnerable species into three will result, at the very least, in three Vulnerable species. Availability of material from collections has also revealed new species including *A. bergii* E.M. Gardner, Zerega, and Arifiani (Gardner et al., in review), a close ally of breadfruit from the Maluku Islands and *A. montanus* E.M. Gardner and N.J.C. Zerega (Gardner and Zerega, in review), a montane species endemic to Vietnam.

### Impact of various analysis methods

Analyzing data in different ways can help produce more robust phylogenetic hypotheses by revealing which relationships are independent of analysis method. Of the variants we tested, codon partitioning had the smallest impact, resulting in no major topological changes except for the relationship of *A. sepicanus* and *A. altissimus*. This is not surprising, as RAxML’s GTRCAT model provides for rate heterogeneity even absent explicit partitioning (Stamatakis 2006). The other comparisons revealed more disagreement, mostly at shallow phylogenetic depths. However, in all cases, disagreement decreased if additional sequences (paralogs or non-coding) were added to a dataset. This suggests more data can lead to a certain amount of convergence in analyses, even though simply adding more data to a supermatrix may not improve the accuracy of the resulting species tree (Degnan and Rosenberg 2009).

Based on these results, we conclude that for our study, greater benefit resulted from analyzing more data, in particular noncoding sequences, than from partitioning by codon position, and the same may be true for analyses at similar phylogenetic scales, particularly when methods provide for rate heterogeneity. Moreover, the results of attempts to produce a codon-partitioned *supercontig* data set suggest that overpartitioning may bias analyses, particularly in the presence of missing data; the ASTRAL analyses of that data set, which effectively had sub-partitions because each gene tree containing three partitions was estimated separately, was more congruent with the main analyses (Appendix 2).

We therefore recommend that when possible, flanking non-coding sequences be included in analyses. The benefits of gene trees with fewer polytomies, and thus more reliable species trees, likely outweigh any minimal advantage gained in partitioning by codon position, at least for a data set like ours. In light of increased congruence between analyses as our data set was enlarged, we suggest using as many loci and as much flanking noncoding sequence as is available, with the caveat to exercise caution with regard to taxa with excessive missing data. The cutoffs we used, >20% of the average sequence length and >∼20% of loci, might be made more stringent, as some inter-analysis disagreement appeared to center around samples with more missing data.

Although adding or extending loci may reduce disagreement between analyses, it may not always increase phylogenetic resolution. A handful of genes may have insufficient informative characters to resolve a phylogeny, and resolution may increase as loci are added, but with hundreds of genes, lack of informative characters is not the problem. Here, consistently high bootstrap values masked substantial gene tree discordance, which actually increased when paralogous loci were added. Other phylogenomic studies have also found high rates of gene tree discordance (Degnan and Rosenberg 2009; Wickett et al. 2014; Copetti et al. 2017; Pease et al. 2018; Liu et al. 2019). Because gene tree discordance can result from biological processes such as incomplete lineage sorting or ancient hybridization, it may reflect not a lack of phylogenetic resolution, but the non-existence of a fixed, absolute species tree. Nonetheless, just as bootstrap support can convey a misleading sense of certainty, support measured by the rate of gene-tree support can exaggerate uncertainty. For example, if a gene tree generally supports a clade, but has one out-of-place taxon, perhaps due to an incomplete or erroneous sequence, that gene tree will not be counted as supporting the clade in question. Support measured as the proportion of gene tree quartets supporting each node, not the frequency of the exact clade being tested, may provide a more realistic measure of support (Sayyari and Mirarab 2016); in our analyses, they were generally lower than bootstrap values but substantially higher than gene-tree support.

### Taxonomic considerations

Our results provide a phylogenetic framework for a taxonomic revision of *Artocarpus*, currently in progress (Table 1). A summary of taxonomic implications is discussed here, and more details with regard to characters can be found in the appendix. The subgeneric divisions made by Jarrett (1959, 1960) and Zerega et al. (2010) can be maintained with minor modifications to account for the anomalous *A. sepicanus* and *A. altissimus*, which in 10/12 main analyses formed a clade. It is curious that these species should be closely allied (having different leaf phyllotaxy and differences in degree of perianth tissue fusion of adjacent pistillate flowers – the defining characters of the subgenera). The disagreement as to their affinity in the codon-partitioned ASTRAL analyses warrants further investigation, raising the possibility that the apparent affinity may be due to long-branch attraction (Roch et al. 2019). The only apparent morphological affinity between them are bifid styles, a moraceous plesiomorphy (Clement and Weiblen 2009), present occasionally in subgenus *Artocarpus* but unique to *A. altissimus* in subgenus *Pseudojaca*.

In addition, the phylogeny supports the broad outlines of Jarrett’s (1959, 1960) sections, validating her careful morphological and anatomical studies, which built on those of Renner (1907). The sections within subgenus *Artocarpus* might be maintained with the inclusion of *A. hirsutus* Lam. and *A. nobilis* Thwaites in section *Duricarpus*. Jarrett noted that those species had characters intermediate between sections *Artoarpus* and *Duriarpus*, and indeed, their positions in all main analyses were sister to most of the rest of section *Duricarpus*.

At the series level within subgenus *Artocarpus*, a wholesale reconsideration is probably necessary, especially in series *Angusticarpi*, which never formed a consistent clade or grade. *Artocarpus teijsmannii* subsp. *subglabrus*, which differs from *A. sepicanus* only in petiole characters, appears to be conspecific with the latter. Of special interest in the clade containing *A. altilis* are putative new species that are wild relatives of breadfruit (*A. bergii*, endemic to Maluku Islands and one accessions of uncertain affinity from the Bogor Botanical Gardens (*cf. camansi*). The status of *A. horridus* is unclear; one accession fell in its expected place together with other samples from the Moluccas, but the position of the other, sister to the entire clade, must be treated with caution, as that sample had among the highest proportions of missing data.

Within subgenus *Pseudojaca*, to the extent we included multiple accessions per species, our results mostly supported Jarrett’s (1960) revision. The series were largely monophyletic, with the exception of the position of *A. tonkinensis* (with peltate interfloral bracts) nested within the clade distinguished by clavate interfloral bracts. The ancestral state for interfloral bracts is likely peltate (Clement and Weiblen 2009), so *A. tonkinensis* may simply represent a plesiomorphic taxon sister to a derived clade. At the species levels, some taxonomic changes will be necessary. For example, as Williams et al. (2017) found, the four species sunk into *A. lacucha* by Berg et al. (2006) (*A. dadah* Miq., *A. ovatus* Blanco, *A. fretesii*, and *A. vrieseanus* var. *refractus*) do not belong together. Additionally, the subspecies of *A. nitidus* are not all sister to one another, as is also the case in *A. gomezianus* subspecies. However, the varieties of *A. vrieseanus* form a clade.

The Chinese species described since Jarrett’s (1960) revision all belong to Series *Clavati*, Our sampling did not include *A. nanchuanensis*, but this species is morphologically similar to *A. hypargyreus*, and subsequent sequencing after the main analyses were complete confirmed the affinity. We were unable to successfully sequence *A. nigrifolius*, but an examination of the type suggests that it is conspecific with *A. hypargyreus*.

Pending a complete revision, we propose the following adjustments to achieve monophyletic sections: *Artocarpus hirsutus* and *A. nobilis* are transferred to sect. *Duricarpus*, and *Artocarpus teijsmannii* subsp. *subglabrus* is reduced to the synonomy of *A. sepicanus*.

## Conclusion

We provide a robust phylogenetic framework for *Artocarpus*, making use of herbarium specimens up to 106 years old to supplement our own collections and achieve near-complete taxon sampling, demonstrating the value of even very old natural history collections in improving phylogenetic studies. Our results will inform future evolutionary and systematic studies of this important group of plants. More generally, the results may guide future analyses of HybSeq datasets, particularly those combining fresh with museum material, by counseling careful attention to dataset construction and analysis method to produce the most informative phylogenetic hypotheses.

The increasing availability of phylogenomic datasets has dramatically changed the practice of revisionary systematics. Datasets containing hundreds or thousands of loci produce trees with extremely high statistical support, apparently providing ironclad frameworks for making taxonomic decisions. However, apparent high support for relationships may often be an artifact of the massive number of characters available for phylogenetic inference, masking real uncertainties, revealed only by employing a variety of analytical methods. By the same token, focusing on exclusively conserved coding regions—an inherent feature of some reference-based assembly methods—can result in unnecessarily uninformative gene trees, leading to poor support at the species tree level. Using a data set with near-complete taxon sampling, we demonstrated that decisions made in how to conduct analyses can substantially affect phylogenetic reconstruction, resulting in discordant phylogenies, each with high statistical support. Employing multiple analytical methods can help separate truly robust phylogenetic relationships from those that only appear to be well-supported but are inconsistent across analyses. While codon partitioning and model choice did not substantially alter our phylogeny, inclusion of flanking non-coding sequences in analyses significantly increased the number of informative splits at the gene tree level, resulting ultimately in more robust species trees. In general, increasing the size of datasets, through inclusion of paralogous genes, increased convergence between analysis methods without reducing gene tree conflict. This likely resulted from biological, not analytical processes; for this reason, we prefer quartet-based scoring methods as the most informative ways of determining support for species trees.

## Acknowledgements

We thank Postar Miun, Jeisin Jumian, Markus Gubilil, Aloysius Laim, S. Brono, Jegong anak Suka, Salang anak Nyegang, Jugah anak Tagi, Wan Nuur Fatiha Wan Zakaria, and Harto for assistance in the field; the Pritzker Laboratory for Molecular Systematics at the Field Museum of Natural History (K. Feldheim) for the use of sequencing facilities; J. Fant, E. Williams, H. Noble, R. Overson, and B. Cooper for assistance in the lab; the Sabah Agriculture Department for access to collections; and the following herbaria for access to collections for examination and/or sampling: BM, BKF, BO, F, IBSC, K, FRIM, FTBG, L, MIN, MO, NY, PNH, SAN, SING, SAR, SNP.

