## Supplemental figures and captions for "Paralogs and off-target sequences improve phylogenetic resolution in a densely-sampled study of the breadfruit genus (*Artocarpus*, Moraceae)"

Figure S1. (A) Heatmap of gene recovery as a percentage of the average recovered sequence length for each gene. Rows represent samples, and columns represent genes. Darker colors indicate more complete recovery; white indicates no recovery. (B) Age (x-axis) for each sample. (C) Number of reads on target (x-axis) for each sample.

Figure S2. Comparison between herbarium sample age and total genes recovered (top) and mean gene length as a proportion of average gene length for each gene (bottom)

Figure S3. BioAnalyzer traces of genomic DNA extracted from nine herbarium samples spanning a one-hundred year age range, showing fragment size (x-axis) versus fragment quantity (y-axis).

Figure S4. Comparison of trees with and without partitioning by codon position for the supermatrix “exon” analyses for all loci. For each pair of trees, incongruent branches are colored red.

Figure S5. Comparison of trees with and without partitioning by codon position for the ASTRAL “exon” analyses for all loci. For each pair of trees, incongruent branches are colored red.

Figure S6. Comparison of trees with and without partitioning by codon position for the supermatrix “exon” analyses, without paralogs. For each pair of trees, incongruent branches are colored red.

Figure S7. Comparison of trees with and without partitioning by codon position for the ASTRAL “exon” analyses, without paralogs. For each pair of trees, incongruent branches are colored red.

Figure S8. Comparison of trees estimated under the GTRCAT and GTRGAMMA models for supermatrix “exon” analyses, partitioned only by gene. For each pair of trees, incongruent branches are colored red.

Figure S9. Comparison of trees estimated under the GTRCAT and GTRGAMMA models for supermatrix “exon” analyses, partitioned by codon position. For each pair of trees, incongruent branches are colored red.

Figure S10. Comparison of species trees based on gene trees estimated under the GTRCAT and GTRGAMMA models for ASTRAL “exon” analyses, not partitioned within gene alignments. For each pair of trees, incongruent branches are colored red.

Figure S11. Comparison of species trees based on gene trees estimated under the GTRCAT and GTRGAMMA models for ASTRAL “exon” analyses, partitioned by codon position. For each pair of trees, incongruent branches are colored red.

Figure S12. PCA of Robinson-Foulds (RF) distances analyses of the “exon” data set conducted under the GTRCAT and GTRGAMMA models.

Figure S13. Comparison of trees based on the core 333 genes and all 517 genes (including 184 paralogs) for the supermatrix “exon” analyses, partitioned only by gene. For each pair of trees, incongruent branches are colored red.

Figure S14. Comparison of trees based on the core 333 genes and all 517 genes (including 184 paralogs) for the ASTRAL “exon” analyses, not partitioned within gene alignments. For each pair of trees, incongruent branches are colored red.

Figure S15. Comparison of trees based on the core 333 genes and all 517 genes (including 184 paralogs) for the supermatrix “supercontig” analyses, partitioned only by gene. For each pair of trees, incongruent branches are colored red.

Figure S16. Comparison of trees based on the core 333 genes and all 517 genes (including 184 paralogs) for the ASTRAL “supercontig” analyses, not partitioned within gene alignments. For each pair of trees, incongruent branches are colored red.

Figure S17. Comparison of trees with (“supercontig”) and without (“exon”) non-coding sequences for supermatrix analyses, partitioned by gene only. For each pair of trees, incongruent branches are colored red.

Figure S18. Comparison of trees with (“supercontig”) and without (“exon”) non-coding sequences for ASTRAL analyses, partitioned by gene only. For each pair of trees, incongruent branches are colored red.

Figure S19. Comparison of trees with (“supercontig”) and without (“exon”) non-coding sequences for supermatrix analyses, partitioned by gene only, without paralogs. For each pair of trees, incongruent branches are colored red.

Figure S20. Comparison of trees with (“supercontig”) and without (“exon”) non-coding sequences for ASTRAL analyses, partitioned by gene only, without paralogs. For each pair of trees, incongruent branches are colored red.

Figure S21. Comparison of trees estimated from concatenated supermatrices (RAxML) and reconciled gene trees (ASTRAL), based on all 517 genes, partitioned by gene only and not by codon position, for the “exon” dataset. For each pair of trees, incongruent branches are colored red.

Figure S22. Comparison of trees estimated from concatenated supermatrices (RAxML) and reconciled gene trees (ASTRAL), based on 333 genes, partitioned by gene only and not by codon position, for the “exon” dataset, excluding paralogs. For each pair of trees, incongruent branches are colored red.

Figure S23. Comparison of trees estimated from concatenated supermatrices (RAxML) and reconciled gene trees (ASTRAL), based on 333 genes, partitioned by gene only and not by codon position, for the “supercontig” dataset, excluding paralogs. For each pair of trees, incongruent branches are colored red.

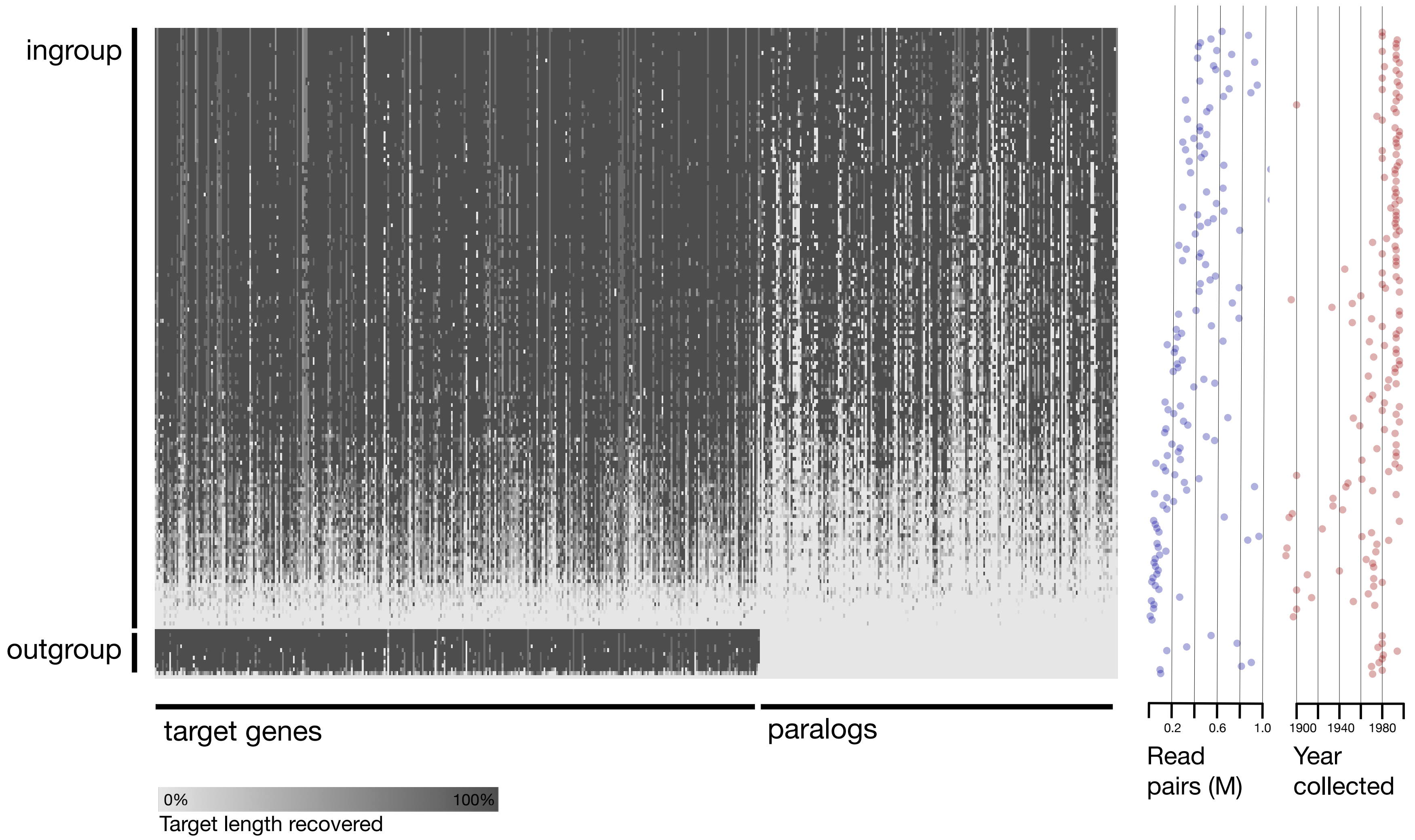

ingroup
outgroup
target genes
paralogs
0%
100%
Target length recovered
0.2
0.6
1.0
Read pairs (M)
1900
1940
1980
Year collected

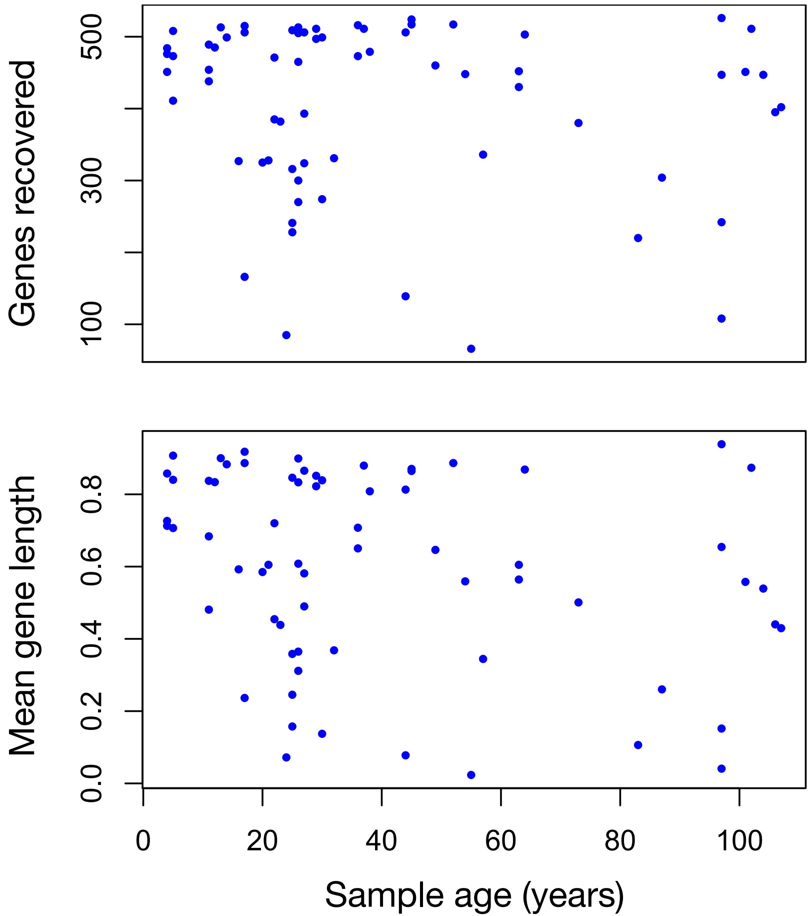

Genes recovered
500
300
100
Mean gene length
0.8
0.6
0.4
0.2
0.0
0
20
40
60
80
100
Sample age (years)

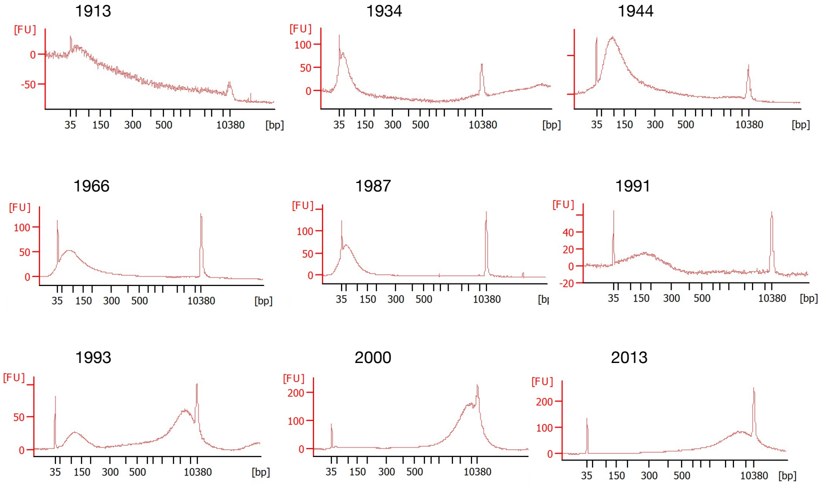
1913
[FU]
0
-50
35
150
300
500
10380
[bp]
1934
[FU]
100
50
0
35
150
300
500
10380
[bp]
1944
35
150
300
500
10380
[bp]
1966
[FU]
100
50
0
35
150
300
500
10380
[bp]
1987
[FU]
100
50
0
35
150
300
500
10380
1991
[FU]
40
20
0
-20
35
150
300
500
10380
[bp]
1993
[FU]
50
0
35
150
300
500
10380
[bp]
2000
[FU]
200
100
0
35
150
300
500
10380
2013
[FU]
200
100
0
35
150
300
500
10380
[bp]

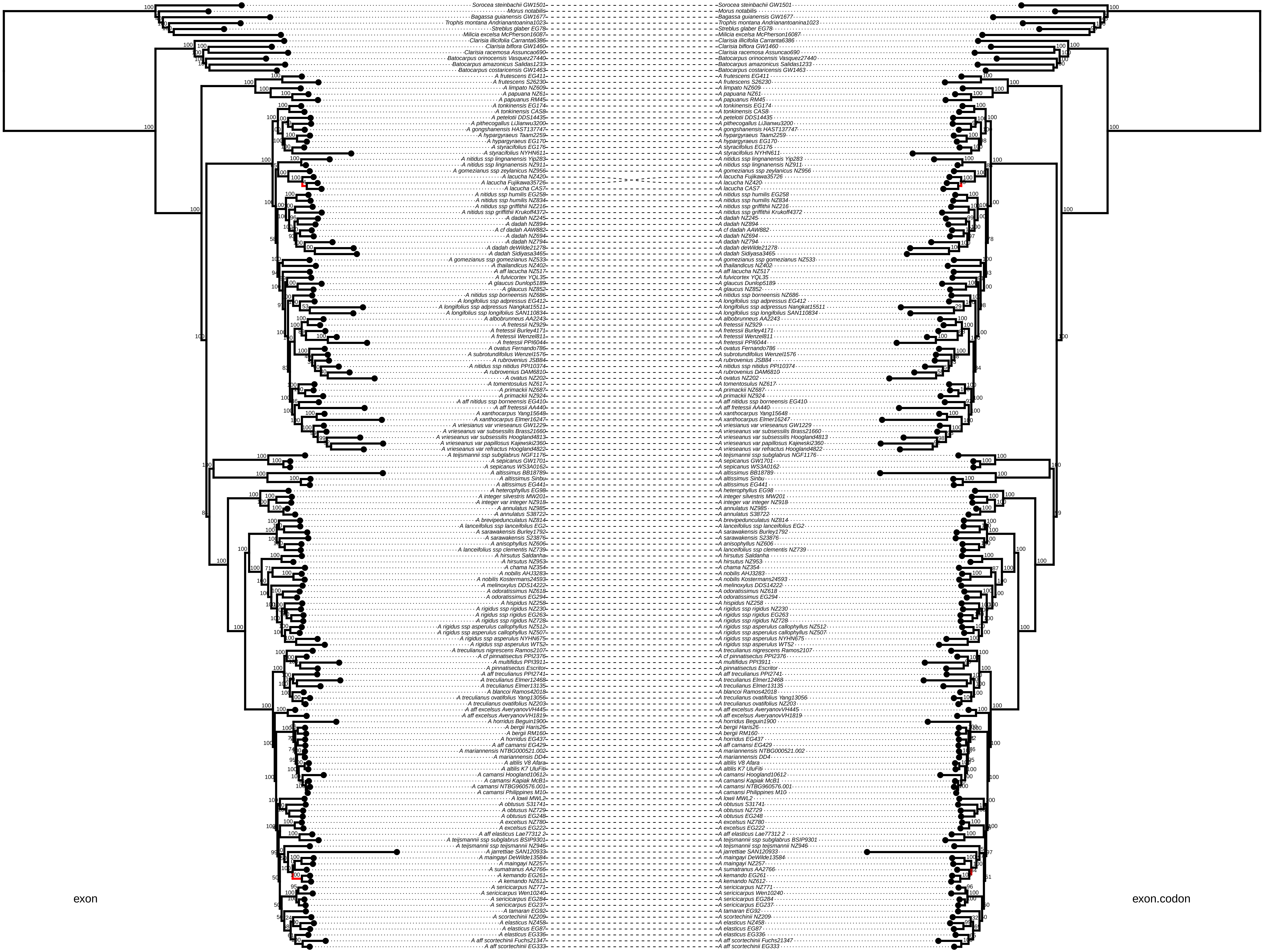

exon
exon.codon
Sorocea steinbachii GW1501
Morus notabilis
Bagassa guianensis GW1677
Trophis montana Andrianantoanina1023
Streblus glaber EG78
Milicia excelsa McPherson16087
Clarisia illicifolia Carranta6386
Clarisia biflora GW1460
Clarisia racemosa Assuncao690
Batocarpus orinocensis Vasquez27440
Batocarpus amazonicus Salidas1233
Batocarpus costaricensis GW1463
A frutescens EG411
A frutescens S26230
A limpato NZ609
A papuana NZ61
A papuanus RM45
A tonkinensis EG174
A tonkinensis CAS8
A petelotii DDS14435
A pithecogallus LiJianwu3200
A gongshanensis HAST137747
A hypargyraeus Taam2259
A hypargyraeus EG170
A styracifolius EG176
A styracifolius NYHN611
A nitidus ssp lingnanensis Yip283
A nitidus ssp lingnanensis NZ911
A gomezianus ssp zeylanicus NZ956
A lacucha NZ420
A lacucha Fujikawa35726
A lacucha CAS7
A nitidus ssp humilis EG258
A nitidus ssp humilis NZ834
A nitidus ssp griffithii NZ216
A nitidus ssp griffithii Krukoff4372
A dadah NZ245
A dadah NZ894
A cf dadah AAW882
A dadah NZ694
A dadah NZ794
A dadah deWilde21278
A dadah Sidiyasa3465
A gomezianus ssp gomezianus NZ533
A thailandicus NZ402
A aff lacucha NZ517
A fulvicortex YQL35
A glaucus Dunlop5189
A glaucus NZ852
A nitidus ssp borneensis NZ686
A longifolius ssp adpressus EG412
A longifolius ssp adpressus Nangkat15511
A longifolius ssp longifolius SAN110834
A albobrunneus AA2243
A fretessii NZ929
A fretessii Burley4171
A fretessii Wenzel811
A fretessii PPI6044
A ovatus Fernando786
A subrotundifolius Wenzel1576
A rubrovenius JSB84
A nitidus ssp nitidus PPI10374
A rubrovenius DAM6810
A ovatus NZ202
A tomentosulus NZ617
A primackii NZ687
A primackii NZ924
A aff nitidus ssp borneensis EG410
A aff fretessii AA440
A xanthocarpus Yang15648
A xanthocarpus Elmer16247
A vriesianus var vrieseanus GW1229
A vrieseanus var subsessilis Brass21660
A vrieseanus var subsessilis Hoogland4813
A vrieseanus var papillosus Kajewski2360
A vrieseanus var refractus Hoogland4822
A teijsmannii ssp subglabrus NGF1176
A sepicanus GW1701
A sepicanus WS3A0162
A altissimus BB18789
A altissimus Sinbu
A altissimus EG441
A heterophyllus EG98
A integer silvestris MW201
A integer var integer NZ918
A annulatus NZ985
A annulatus S38722
A brevipedunculatus NZ814
A lanceifolius ssp lanceifolius EG2
A sarawakensis Burley1792
A sarawakensis S23876
A anisophyllus NZ606
A lanceifolius ssp clementis NZ739
A hirsutus Saldanha
A hirsutus NZ953
A chama NZ354
A nobilis AHJ3283
A nobilis Kostermans24593
A melinoxylus DDS14222
A odoratissimus NZ618
A odoratissimus EG294
A hispidus NZ258
A rigidus ssp rigidus NZ230
A rigidus ssp rigidus EG263
A rigidus ssp rigidus NZ728
A rigidus ssp asperulus callophyllus NZ512
A rigidus ssp asperulus callophyllus NZ507
A rigidus ssp asperulus NYHN675
A rigidus ssp asperulus WT52
A treculianus nigrescens Ramos2107
A cf pinnatisectus PPI2376
A multifidus PPI3911
A pinnatisectus Escritor
A aff treculianus PPI2741
A treculianus Elmer12468
A treculianus Elmer13135
A blancoi Ramos42018
A treculianus ovatifolius Yang13056
A treculianus ovatifolius NZ203
A aff excelsus AveryanovVH445
A aff excelsus AveryanovVH1819
A horridus Beguin1900
A bergii Haris26
A bergii RM160
A horridus EG437
A aff camansi EG429
A mariannensis NTBG000521.002
A mariannensis DD4
A altilis V8 Afara
A altilis K7 UluFiti
A camansi Hoogland10612
A camansi Kapiak McB1
A camansi NTBG960576.001
A camansi Philippines M10
A lowii MWL2
A obtusus S31741
A obtusus NZ729
A obtusus EG248
A excelsus NZ780
A excelsus EG222
A aff elasticus Lae77312 2
A teijsmannii ssp subglabrus BSIP9301
A teijsmannii ssp teijsmannii NZ946
A jarrettiae SAN120933
A maingayi DeWilde13584
A maingayi NZ257
A sumatranus AA2766
A kemando EG261
A kemando NZ612
A sericicarpus NZ771
A sericicarpus Wen10240
A sericicarpus EG284
A sericicarpus EG237
A tamaran EG92
A scortechinii NZ209
A elasticus NZ458
A elasticus EG87
A elasticus EG336
A aff scortechinii Fuchs21347
A aff scortechinii EG333

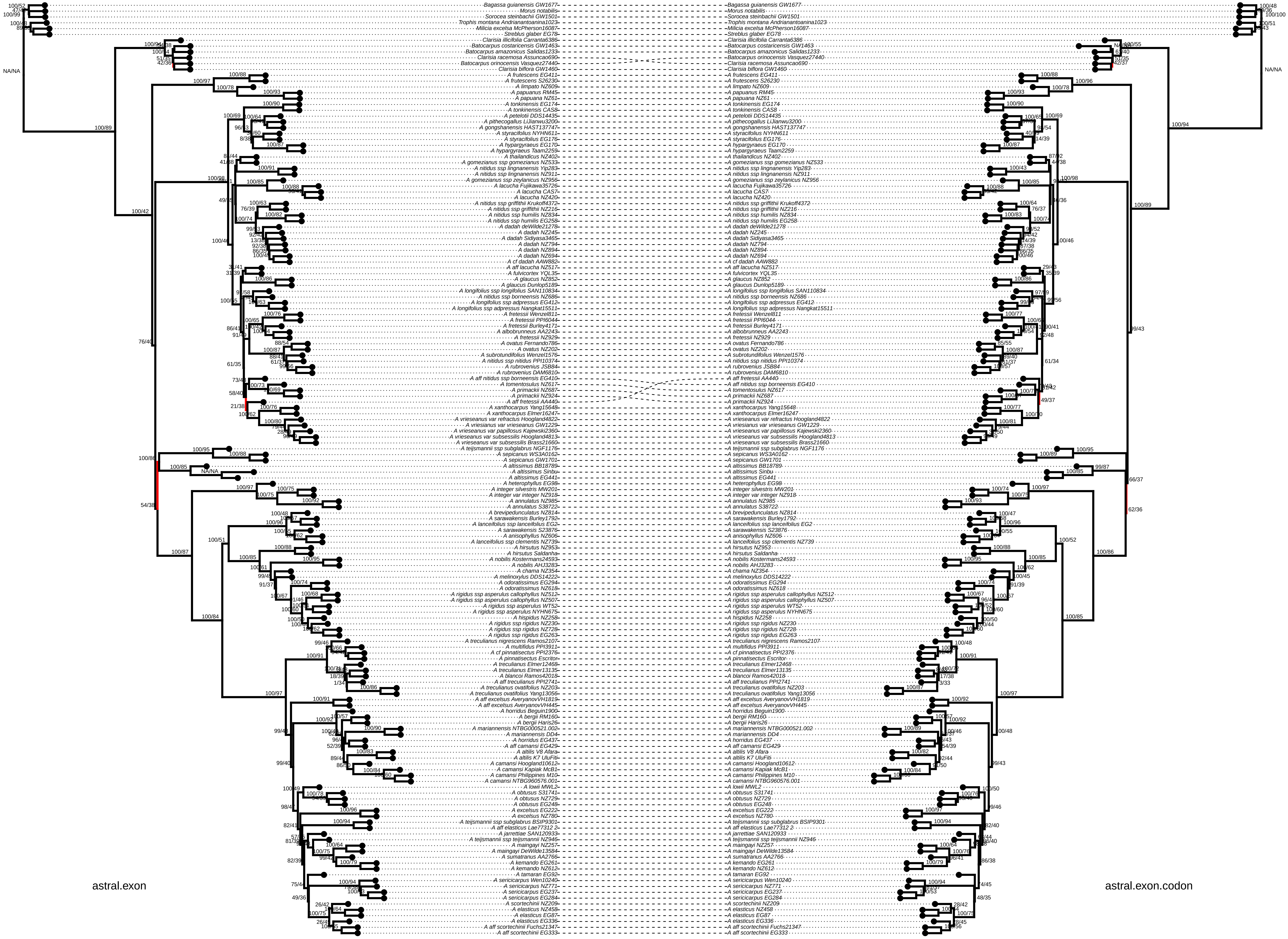
astral.exon
astral.exon.codon
Bagassa guianensis GW1677
Morus notabilis
Sorocea steinbachii GW1501
Trophis montana Andrianantoanina1023
Milicia excelsa McPherson16087
Streblus glaber EG78
Clarisia illicifolia Carranta6386
Batocarpus costaricensis GW1463
Batocarpus amazonicus Salidas1233
Clarisia racemosa Assuncao690
Batocarpus orinocensis Vasquez27440
Clarisia biflora GW1460
A frutescens EG411
A frutescens S26230
A limpato NZ609
A papuanus RM45
A papuana NZ61
A tonkinensis EG174
A tonkinensis CAS8
A petelotii DDS14435
A pithecogallus LiJianwu3200
A gongshanensis HAST137747
A styracifolius NYHN611
A styracifolius EG176
A hypargyraeus EG170
A hypargyraeus Taam2259
A thailandicus NZ402
A gomezianus ssp gomezianus NZ533
A nitidus ssp lingnanensis Yip283
A nitidus ssp lingnanensis NZ911
A gomezianus ssp zeylanicus NZ956
A lacucha Fujikawa35726
A lacucha CAS7
A lacucha NZ420
A nitidus ssp griffithii Krukoff4372
A nitidus ssp griffithii NZ216
A nitidus ssp humilis NZ834
A nitidus ssp humilis EG258
A dadah deWilde21278
A dadah NZ245
A dadah Sidiyasa3465
A dadah NZ794
A dadah NZ894
A dadah NZ694
A cf dadah AAW882
A aff lacucha NZ517
A fulvicortex YQL35
A glaucus NZ852
A glaucus Dunlop5189
A longifolius ssp longifolius SAN110834
A nitidus ssp borneensis NZ686
A longifolius ssp adpressus EG412
A longifolius ssp adpressus Nangkat15511
A fretessii Wenzel811
A fretessii PPI6044
A fretessii Burley4171
A albobrunneus AA2243
A fretessii NZ929
A ovatus Fernando786
A ovatus NZ202
A subrotundifolius Wenzel1576
A nitidus ssp nitidus PPI10374
A rubrovenius JSB84
A rubrovenius DAM6810
A aff nitidus ssp borneensis EG410
A tomentosulus NZ617
A primackii NZ687
A primackii NZ924
A aff fretessii AA440
A xanthocarpus Yang15648
A xanthocarpus Elmer16247
A vrieseanus var refractus Hoogland4822
A vriesianus var vrieseanus GW1229
A vrieseanus var papillosus Kajewski2360
A vrieseanus var subsessilis Hoogland4813
A vrieseanus var subsessilis Brass21660
A teijsmannii ssp subglabrus NGF1176
A sepicanus WS3A0162
A sepicanus GW1701
A altissimus BB18789
A altissimus Sinbu
A altissimus EG441
A heterophyllus EG98
A integer silvestris MW201
A integer var integer NZ918
A annulatus NZ985
A annulatus S38722
A brevipedunculatus NZ814
A sarawakensis Burley1792
A lanceifolius ssp lanceifolius EG2
A sarawakensis S23876
A anisophyllus NZ606
A lanceifolius ssp clementis NZ739
A hirsutus NZ953
A hirsutus Saldanha
A nobilis Kostermans24593
A nobilis AHJ3283
A chama NZ354
A melinoxylus DDS14222
A odoratissimus EG294
A odoratissimus NZ618
A rigidus ssp asperulus callophyllus NZ512
A rigidus ssp asperulus callophyllus NZ507
A rigidus ssp asperulus WT52
A rigidus ssp asperulus NYHN675
A hispidus NZ258
A rigidus ssp rigidus NZ230
A rigidus ssp rigidus NZ728
A rigidus ssp rigidus EG263
A treculianus nigrescens Ramos2107
A multifidus PPI3911
A cf pinnatisectus PPI2376
A pinnatisectus Escritor
A treculianus Elmer12468
A treculianus Elmer13135
A blancoi Ramos42018
A aff treculianus PPI2741
A treculianus ovatifolius NZ203
A treculianus ovatifolius Yang13056
A aff excelsus AveryanovVH1819
A aff excelsus AveryanovVH445
A horridus Beguin1900
A bergii RM160
A bergii Haris26
A mariannensis NTBG000521.002
A mariannensis DD4
A horridus EG437
A aff camansi EG429
A altilis V8 Afara
A altilis K7 UluFiti
A camansi Hoogland10612
A camansi Kapiak McB1
A camansi Philippines M10
A camansi NTBG960576.001
A lowii MWL2
A obtusus S31741
A obtusus NZ729
A obtusus EG248
A excelsus EG222
A excelsus NZ780
A teijsmannii ssp subglabrus BSIP9301
A aff elasticus Lae77312 2
A jarrettiae SAN120933
A teijsmannii ssp teijsmannii NZ946
A maingayi NZ257
A maingayi DeWilde13584
A sumatranus AA2766
A kemando EG261
A kemando NZ612
A tamaran EG92
A sericicarpus Wen10240
A sericicarpus NZ771
A sericicarpus EG237
A sericicarpus EG284
A scortechinii NZ209
A elasticus NZ458
A elasticus EG87
A elasticus EG336
A aff scortechinii Fuchs21347
A aff scortechinii EG333

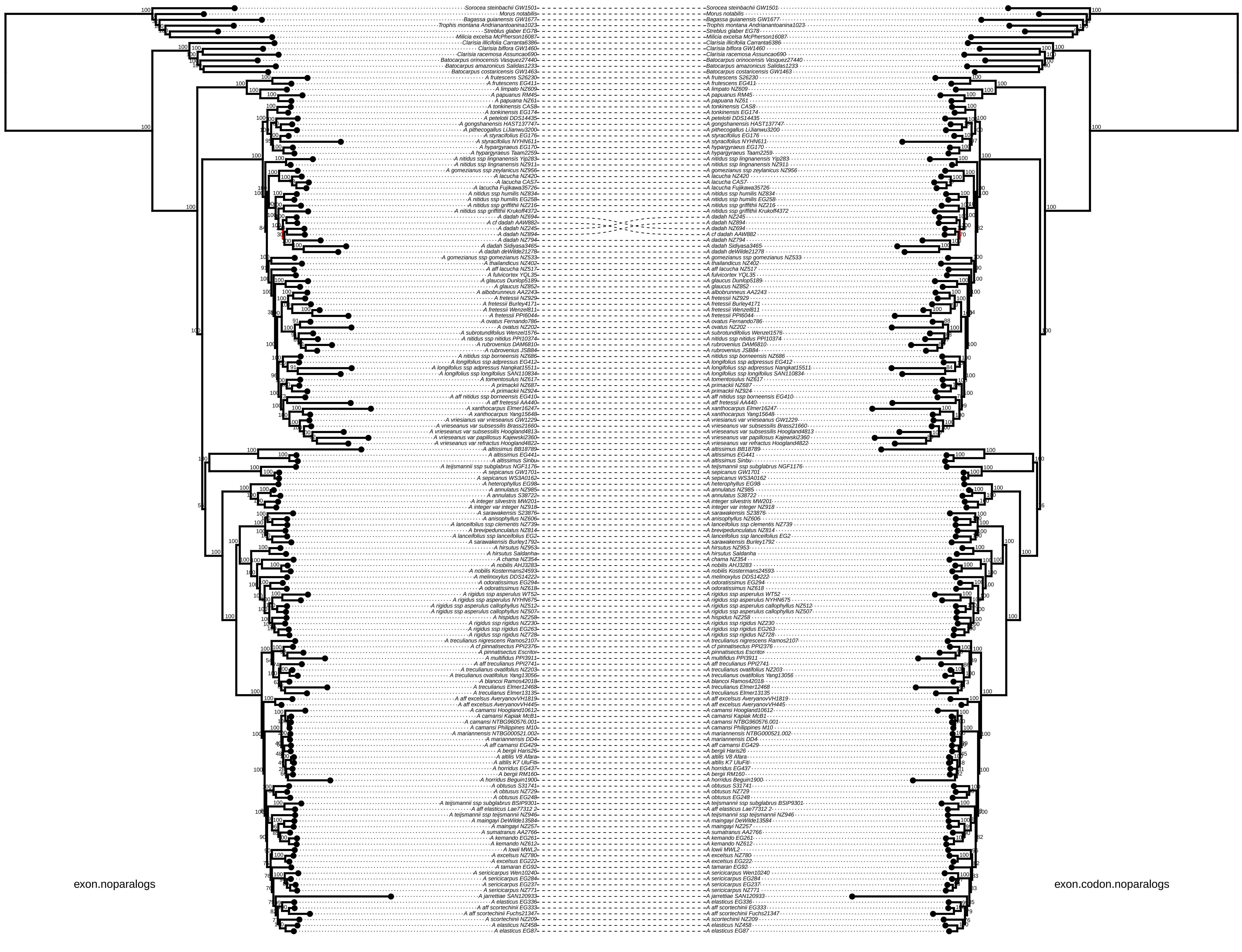
exon.noparalogs
exon.codon.noparalogs
Sorocea steinbachii GW1501
Morus notabilis
Bagassa guianensis GW1677
Trophis montana Andrianantoanina1023
Streblus glaber EG78
Milicia excelsa McPherson16087
Clarisia illicifolia Carranta6386
Clarisia biflora GW1460
Clarisia racemosa Assuncao690
Batocarpus orinocensis Vasquez27440
Batocarpus amazonicus Salidas1233
Batocarpus costaricensis GW1463
A frutescens S26230
A frutescens EG411
A limpato NZ609
A papuanus RM45
A papuana NZ61
A tonkinensis CAS8
A tonkinensis EG174
A petelotii DDS14435
A gongshanensis HAST137747
A pithecogallus LiJianwu3200
A styracifolius EG176
A styracifolius NYHN611
A hypargyraeus EG170
A hypargyraeus Taam2259
A nitidus ssp lingnanensis Yip283
A nitidus ssp lingnanensis NZ911
A gomezianus ssp zeylanicus NZ956
A lacucha NZ420
A lacucha CAS7
A lacucha Fujikawa35726
A nitidus ssp humilis NZ834
A nitidus ssp humilis EG258
A nitidus ssp griffithii NZ216
A nitidus ssp griffithii Krukoff4372
A dadah NZ694
A cf dadah AAW882
A dadah NZ245
A dadah NZ894
A dadah NZ794
A dadah Sidiyasa3465
A dadah deWilde21278
A gomezianus ssp gomezianus NZ533
A thailandicus NZ402
A aff lacucha NZ517
A fulvicortex YQL35
A glaucus Dunlop5189
A glaucus NZ852
A albobrunneus AA2243
A fretessii NZ929
A fretessii Burley4171
A fretessii Wenzel811
A fretessii PPI6044
A ovatus Fernando786
A ovatus NZ202
A subrotundifolius Wenzel1576
A nitidus ssp nitidus PPI10374
A rubrovenius DAM6810
A rubrovenius JSB84
A nitidus ssp borneensis NZ686
A longifolius ssp adpressus EG412
A longifolius ssp adpressus Nangkat15511
A longifolius ssp longifolius SAN110834
A tomentosulus NZ617
A primackii NZ687
A primackii NZ924
A aff nitidus ssp borneensis EG410
A aff fretessii AA440
A xanthocarpus Elmer16247
A xanthocarpus Yang15648
A vriesianus var vrieseanus GW1229
A vrieseanus var subsessilis Brass21660
A vrieseanus var subsessilis Hoogland4813
A vrieseanus var papillosus Kajewski2360
A vrieseanus var refractus Hoogland4822
A altissimus BB18789
A altissimus EG441
A altissimus Sinbu
A teijsmannii ssp subglabrus NGF1176
A sepicanus GW1701
A sepicanus WS3A0162
A heterophyllus EG98
A annulatus NZ985
A annulatus S38722
A integer silvestris MW201
A integer var integer NZ918
A sarawakensis S23876
A anisophyllus NZ606
A lanceifolius ssp clementis NZ739
A brevipedunculatus NZ814
A lanceifolius ssp lanceifolius EG2
A sarawakensis Burley1792
A hirsutus NZ953
A hirsutus Saldanha
A chama NZ354
A nobilis AHJ3283
A nobilis Kostermans24593
A melinoxylus DDS14222
A odoratissimus EG294
A odoratissimus NZ618
A rigidus ssp asperulus WT52
A rigidus ssp asperulus NYHN675
A rigidus ssp asperulus callophyllus NZ512
A rigidus ssp asperulus callophyllus NZ507
A hispidus NZ258
A rigidus ssp rigidus NZ230
A rigidus ssp rigidus EG263
A rigidus ssp rigidus NZ728
A treculianus nigrescens Ramos2107
A cf pinnatisectus PPI2376
A pinnatisectus Escritor
A multifidus PPI3911
A aff treculianus PPI2741
A treculianus ovatifolius NZ203
A treculianus ovatifolius Yang13056
A blancoi Ramos42018
A treculianus Elmer12468
A treculianus Elmer13135
A aff excelsus AveryanovVH1819
A aff excelsus AveryanovVH445
A camansi Hoogland10612
A camansi Kapiak McB1
A camansi NTBG960576.001
A camansi Philippines M10
A mariannensis NTBG000521.002
A mariannensis DD4
A aff camansi EG429
A bergii Haris26
A altilis V8 Afara
A altilis K7 UluFiti
A horridus EG437
A bergii RM160
A horridus Beguin1900
A obtusus S31741
A obtusus NZ729
A obtusus EG248
A teijsmannii ssp subglabrus BSIP9301
A aff elasticus Lae77312 2
A teijsmannii ssp teijsmannii NZ946
A maingayi DeWilde13584
A maingayi NZ257
A sumatranus AA2766
A kemando EG261
A kemando NZ612
A lowii MWL2
A excelsus NZ780
A excelsus EG222
A tamaran EG92
A sericicarpus Wen10240
A sericicarpus EG284
A sericicarpus EG237
A sericicarpus NZ771
A jarrettiae SAN120933
A elasticus EG336
A aff scortechinii EG333
A aff scortechinii Fuchs21347
A scortechinii NZ209
A elasticus NZ458
A elasticus EG87

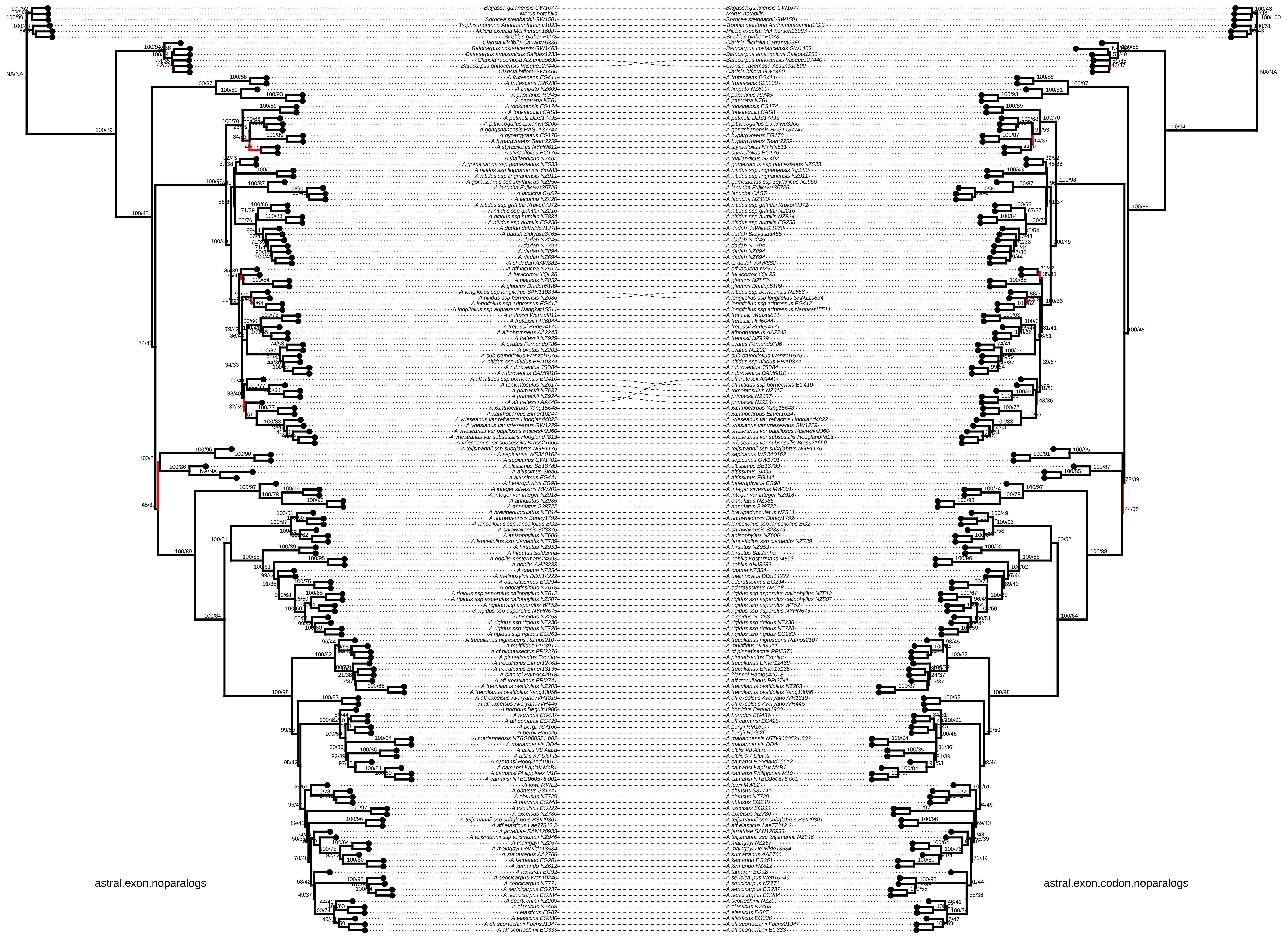
astral.exon.noparalogs
astral.exon.codon.noparalogs

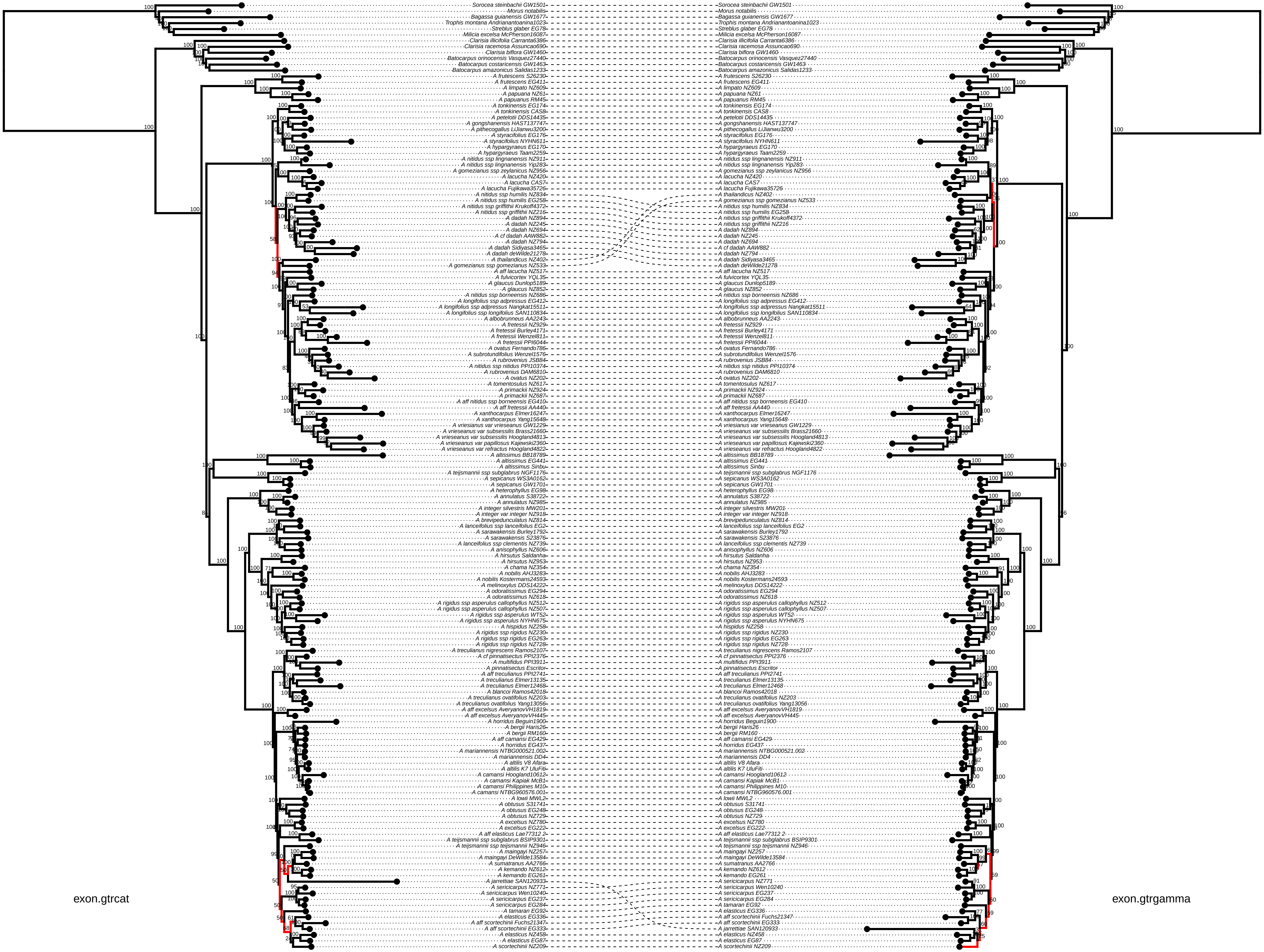
exon.gtrcat
exon.gtrgamma
Sorocea steinbachii GW1501
Morus notabilis
Bagassa guianensis GW1677
Trophis montana Andrianantoanina1023
Streblus glaber EG78
Milicia excelsa McPherson16087
Clarisia illicifolia Carranta6386
Clarisia racemosa Assuncao690
Clarisia biflora GW1460
Batocarpus orinocensis Vasquez27440
Batocarpus costaricensis GW1463
Batocarpus amazonicus Salidas1233
A frutescens S26230
A frutescens EG411
A limpato NZ609
A papuana NZ61
A papuanus RM45
A tonkinensis EG174
A tonkinensis CAS8
A petelotii DDS14435
A gongshanensis HAST137747
A pithecogallus LiJianwu3200
A styracifolius EG176
A styracifolius NYHN611
A hypargyraeus EG170
A hypargyraeus Taam2259
A nitidus ssp lingnanensis NZ911
A nitidus ssp lingnanensis Yip283
A gomezianus ssp zeylanicus NZ956
A lacucha NZ420
A lacucha CAS7
A lacucha Fujikawa35726
A nitidus ssp humilis NZ834
A nitidus ssp humilis EG258
A nitidus ssp griffithii Krukoff4372
A nitidus ssp griffithii NZ216
A dadah NZ894
A dadah NZ245
A dadah NZ694
A cf dadah AAW882
A dadah NZ794
A dadah Sidiyasa3465
A dadah deWilde21278
A thailandicus NZ402
A gomezianus ssp gomezianus NZ533
A aff lacucha NZ517
A fulvicortex YQL35
A glaucus Dunlop5189
A glaucus NZ852
A nitidus ssp borneensis NZ686
A longifolius ssp adpressus EG412
A longifolius ssp adpressus Nangkat15511
A longifolius ssp longifolius SAN110834
A albobrunneus AA2243
A fretessii NZ929
A fretessii Burley4171
A fretessii Wenzel811
A fretessii PPI6044
A ovatus Fernando786
A subrotundifolius Wenzel1576
A rubrovenius JSB84
A nitidus ssp nitidus PPI10374
A rubrovenius DAM6810
A ovatus NZ202
A tomentosulus NZ617
A primackii NZ924
A primackii NZ687
A aff nitidus ssp borneensis EG410
A aff fretessii AA440
A xanthocarpus Elmer16247
A xanthocarpus Yang15648
A vriesianus var vrieseanus GW1229
A vrieseanus var subsessilis Brass21660
A vrieseanus var subsessilis Hoogland4813
A vrieseanus var papillosus Kajewski2360
A vrieseanus var refractus Hoogland4822
A altissimus BB18789
A altissimus EG441
A altissimus Sinbu
A teijsmannii ssp subglabrus NGF1176
A sepicanus WS3A0162
A sepicanus GW1701
A heterophyllus EG98
A annulatus S38722
A annulatus NZ985
A integer silvestris MW201
A integer var integer NZ918
A brevipedunculatus NZ814
A lanceifolius ssp lanceifolius EG2
A sarawakensis Burley1792
A sarawakensis S23876
A lanceifolius ssp clementis NZ739
A anisophyllus NZ606
A hirsutus Saldanha
A hirsutus NZ953
A chama NZ354
A nobilis AHJ3283
A nobilis Kostermans24593
A melinoxylus DDS14222
A odoratissimus EG294
A odoratissimus NZ618
A rigidus ssp asperulus callophyllus NZ512
A rigidus ssp asperulus callophyllus NZ507
A rigidus ssp asperulus WT52
A rigidus ssp asperulus NYHN675
A hispidus NZ258
A rigidus ssp rigidus NZ230
A rigidus ssp rigidus EG263
A rigidus ssp rigidus NZ728
A treculianus nigrescens Ramos2107
A cf pinnatisectus PPI2376
A multifidus PPI3911
A pinnatisectus Escritor
A aff treculianus PPI2741
A treculianus Elmer13135
A treculianus Elmer12468
A blancoi Ramos42018
A treculianus ovatifolius NZ203
A treculianus ovatifolius Yang13056
A aff excelsus AveryanovVH1819
A aff excelsus AveryanovVH445
A horridus Beguin1900
A bergii Haris26
A bergii RM160
A aff camansi EG429
A horridus EG437
A mariannensis NTBG000521.002
A mariannensis DD4
A altilis V8 Afara
A altilis K7 UluFiti
A camansi Hoogland10612
A camansi Kapiak McB1
A camansi Philippines M10
A camansi NTBG960576.001
A lowii MWL2
A obtusus S31741
A obtusus EG248
A obtusus NZ729
A excelsus NZ780
A excelsus EG222
A aff elasticus Lae77312 2
A teijsmannii ssp subglabrus BSIP9301
A teijsmannii ssp teijsmannii NZ946
A maingayi NZ257
A maingayi DeWilde13584
A sumatranus AA2766
A kemando NZ612
A kemando EG261
A jarrettiae SAN120933
A sericicarpus NZ771
A sericicarpus Wen10240
A sericicarpus EG237
A sericicarpus EG284
A tamaran EG92
A elasticus EG336
A aff scortechinii Fuchs21347
A aff scortechinii EG333
A elasticus NZ458
A elasticus EG87
A scortechinii NZ209

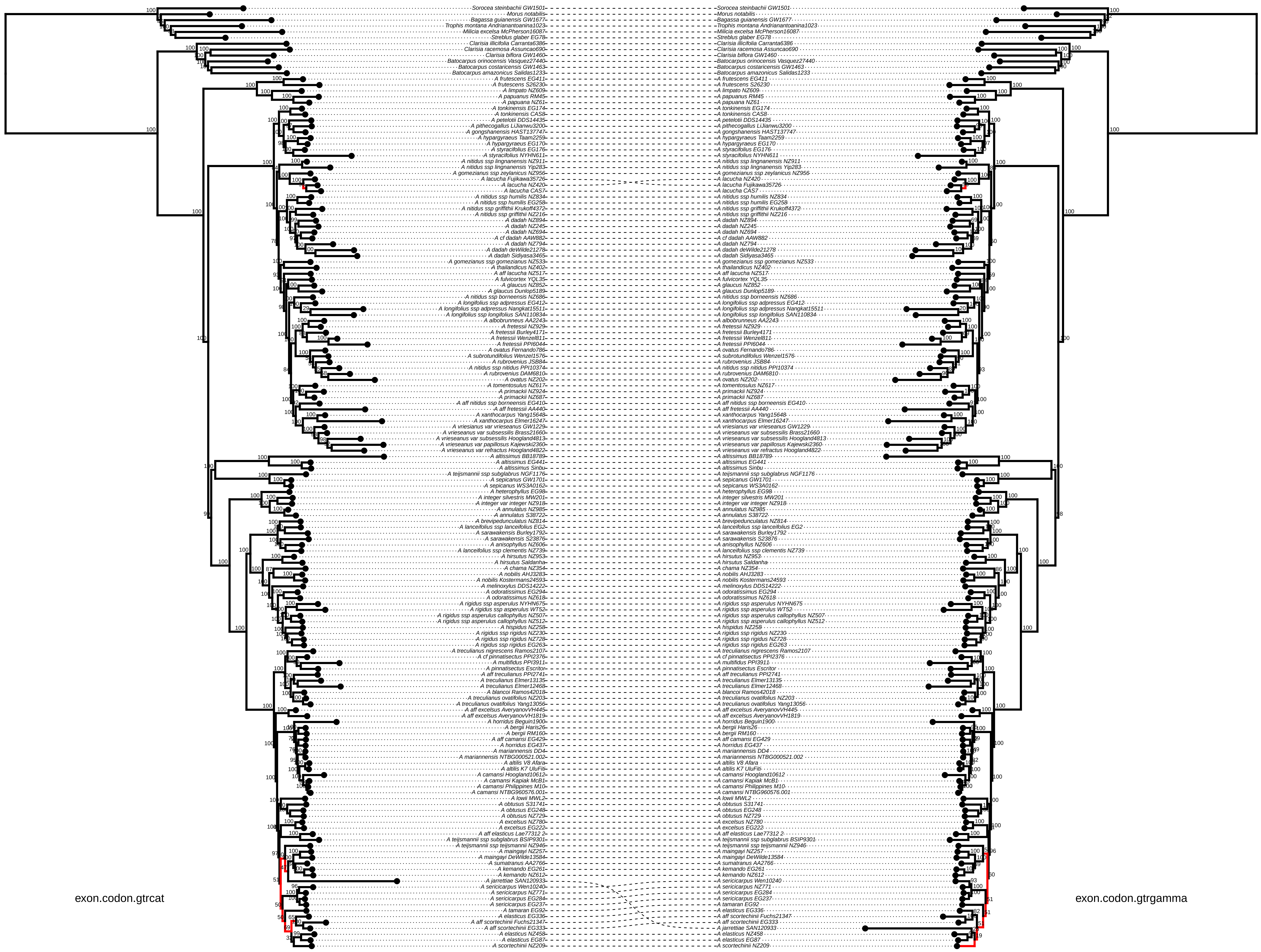
exon.codon.gtrcat
exon.codon.gtrgamma
Sorocea steinbachii GW1501
Morus notabilis
Bagassa guianensis GW1677
Trophis montana Andrianantoanina1023
Milicia excelsa McPherson16087
Streblus glaber EG78
Clarisia illicifolia Carranta6386
Clarisia racemosa Assuncao690
Clarisia biflora GW1460
Batocarpus orinocensis Vasquez27440
Batocarpus costaricensis GW1463
Batocarpus amazonicus Salidas1233
A frutescens EG411
A frutescens S26230
A limpato NZ609
A papuanus RM45
A papuana NZ61
A tonkinensis EG174
A tonkinensis CAS8
A petelotii DDS14435
A pithecogallus LiJianwu3200
A gongshanensis HAST137747
A hypargyraeus Taam2259
A hypargyraeus EG170
A styracifolius EG176
A styracifolius NYHN611
A nitidus ssp lingnanensis NZ911
A nitidus ssp lingnanensis Yip283
A gomezianus ssp zeylanicus NZ956
A lacucha Fujikawa35726
A lacucha NZ420
A lacucha CAS7
A nitidus ssp humilis NZ834
A nitidus ssp humilis EG258
A nitidus ssp griffithii Krukoff4372
A nitidus ssp griffithii NZ216
A dadah NZ894
A dadah NZ245
A dadah NZ694
A cf dadah AAW882
A dadah NZ794
A dadah deWilde21278
A dadah Sidiyasa3465
A gomezianus ssp gomezianus NZ533
A thailandicus NZ402
A aff lacucha NZ517
A fulvicortex YQL35
A glaucus NZ852
A glaucus Dunlop5189
A nitidus ssp borneensis NZ686
A longifolius ssp adpressus EG412
A longifolius ssp adpressus Nangkat15511
A longifolius ssp longifolius SAN110834
A albobrunneus AA2243
A fretessii NZ929
A fretessii Burley4171
A fretessii Wenzel811
A fretessii PPI6044
A ovatus Fernando786
A subrotundifolius Wenzel1576
A rubrovenius JSB84
A nitidus ssp nitidus PPI10374
A rubrovenius DAM6810
A ovatus NZ202
A tomentosulus NZ617
A primackii NZ924
A primackii NZ687
A aff nitidus ssp borneensis EG410
A aff fretessii AA440
A xanthocarpus Yang15648
A xanthocarpus Elmer16247
A vriesianus var vrieseanus GW1229
A vrieseanus var subsessilis Brass21660
A vrieseanus var subsessilis Hoogland4813
A vrieseanus var papillosus Kajewski2360
A vrieseanus var refractus Hoogland4822
A altissimus BB18789
A altissimus EG441
A altissimus Sinbu
A teijsmannii ssp subglabrus NGF1176
A sepicanus GW1701
A sepicanus WS3A0162
A heterophyllus EG98
A integer silvestris MW201
A integer var integer NZ918
A annulatus NZ985
A annulatus S38722
A brevipedunculatus NZ814
A lanceifolius ssp lanceifolius EG2
A sarawakensis Burley1792
A sarawakensis S23876
A anisophyllus NZ606
A lanceifolius ssp clementis NZ739
A hirsutus NZ953
A hirsutus Saldanha
A chama NZ354
A nobilis AHJ3283
A nobilis Kostermans24593
A melinoxylus DDS14222
A odoratissimus EG294
A odoratissimus NZ618
A rigidus ssp asperulus NYHN675
A rigidus ssp asperulus WT52
A rigidus ssp asperulus callophyllus NZ507
A rigidus ssp asperulus callophyllus NZ512
A hispidus NZ258
A rigidus ssp rigidus NZ230
A rigidus ssp rigidus NZ728
A rigidus ssp rigidus EG263
A treculianus nigrescens Ramos2107
A cf pinnatisectus PPI2376
A multifidus PPI3911
A pinnatisectus Escritor
A aff treculianus PPI2741
A treculianus Elmer13135
A treculianus Elmer12468
A blancoi Ramos42018
A treculianus ovatifolius NZ203
A treculianus ovatifolius Yang13056
A aff excelsus AveryanovVH445
A aff excelsus AveryanovVH1819
A horridus Beguin1900
A bergii Haris26
A bergii RM160
A aff camansi EG429
A horridus EG437
A mariannensis DD4
A mariannensis NTBG000521.002
A altilis V8 Afara
A altilis K7 UluFiti
A camansi Hoogland10612
A camansi Kapiak McB1
A camansi Philippines M10
A camansi NTBG960576.001
A lowii MWL2
A obtusus S31741
A obtusus EG248
A obtusus NZ729
A excelsus NZ780
A excelsus EG222
A aff elasticus Lae77312 2
A teijsmannii ssp subglabrus BSIP9301
A teijsmannii ssp teijsmannii NZ946
A maingayi NZ257
A maingayi DeWilde13584
A sumatranus AA2766
A kemando EG261
A kemando NZ612
A jarrettiae SAN120933
A sericicarpus Wen10240
A sericicarpus NZ771
A sericicarpus EG284
A sericicarpus EG237
A tamaran EG92
A elasticus EG336
A aff scortechinii Fuchs21347
A aff scortechinii EG333
A elasticus NZ458
A elasticus EG87
A scortechinii NZ209

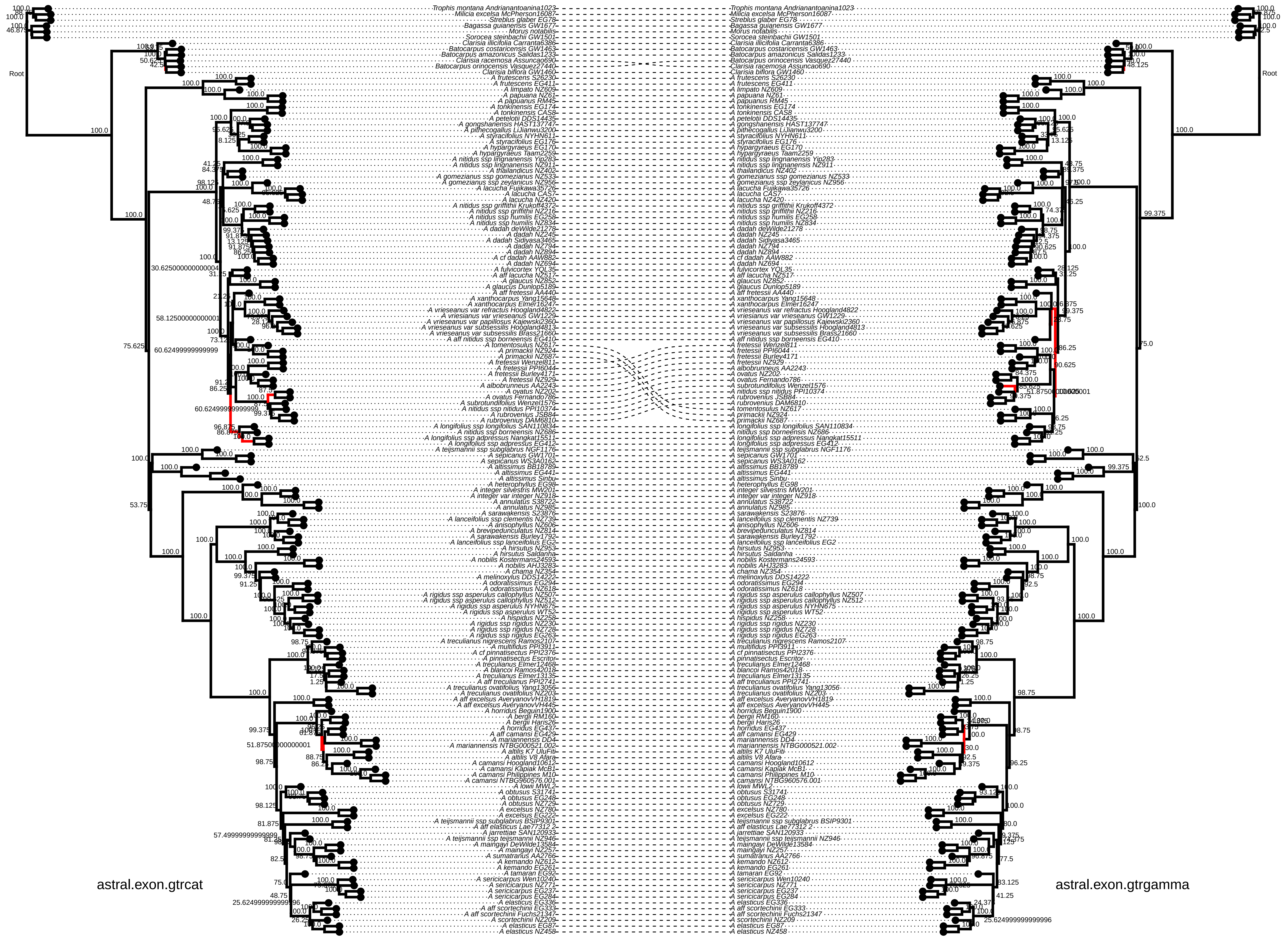
Root
astral.exon.gtrcat
astral.exon.gtrgamma

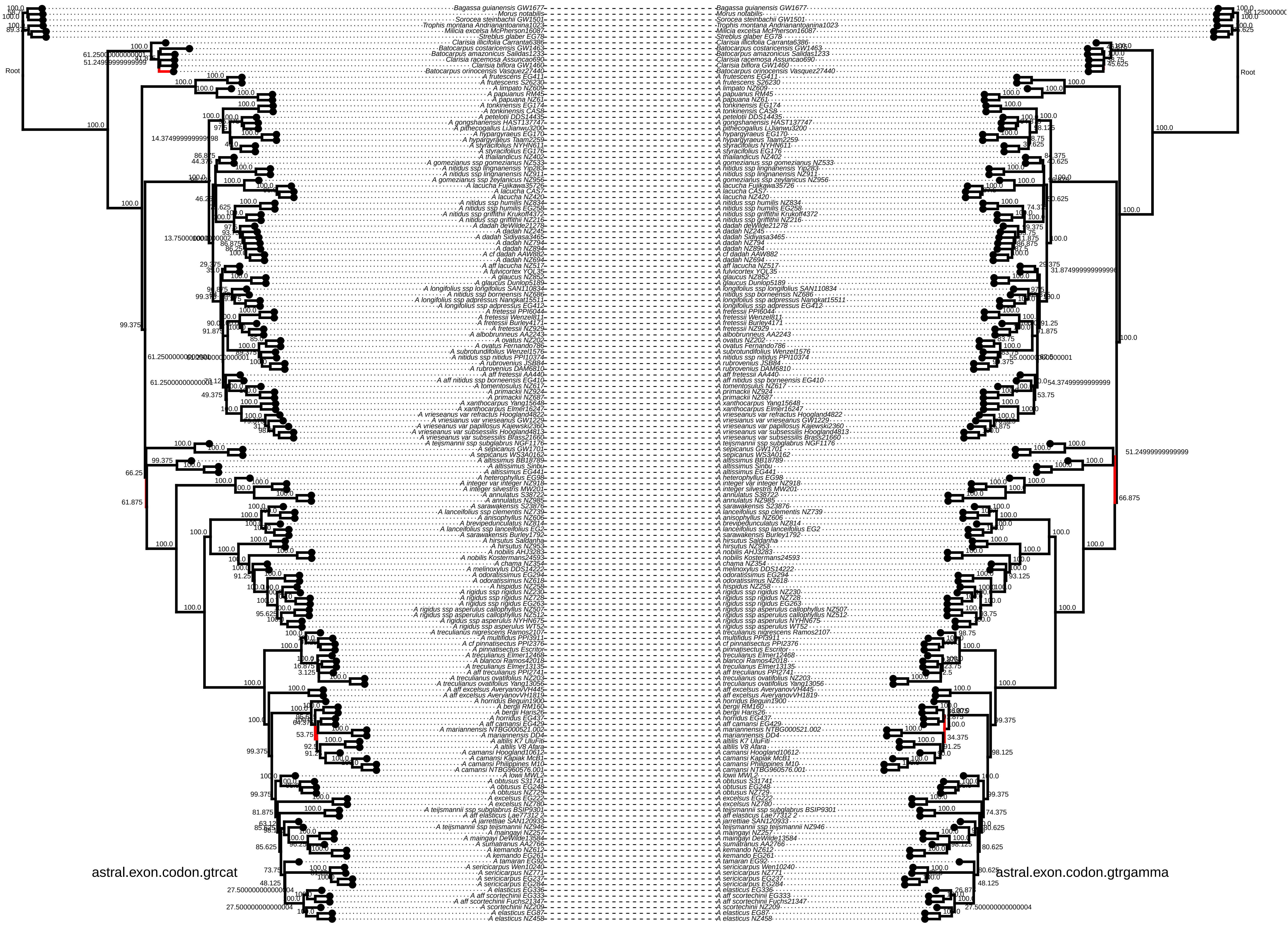
Root
astral.exon.codon.gtrcat
astral.exon.codon.gtrgamma

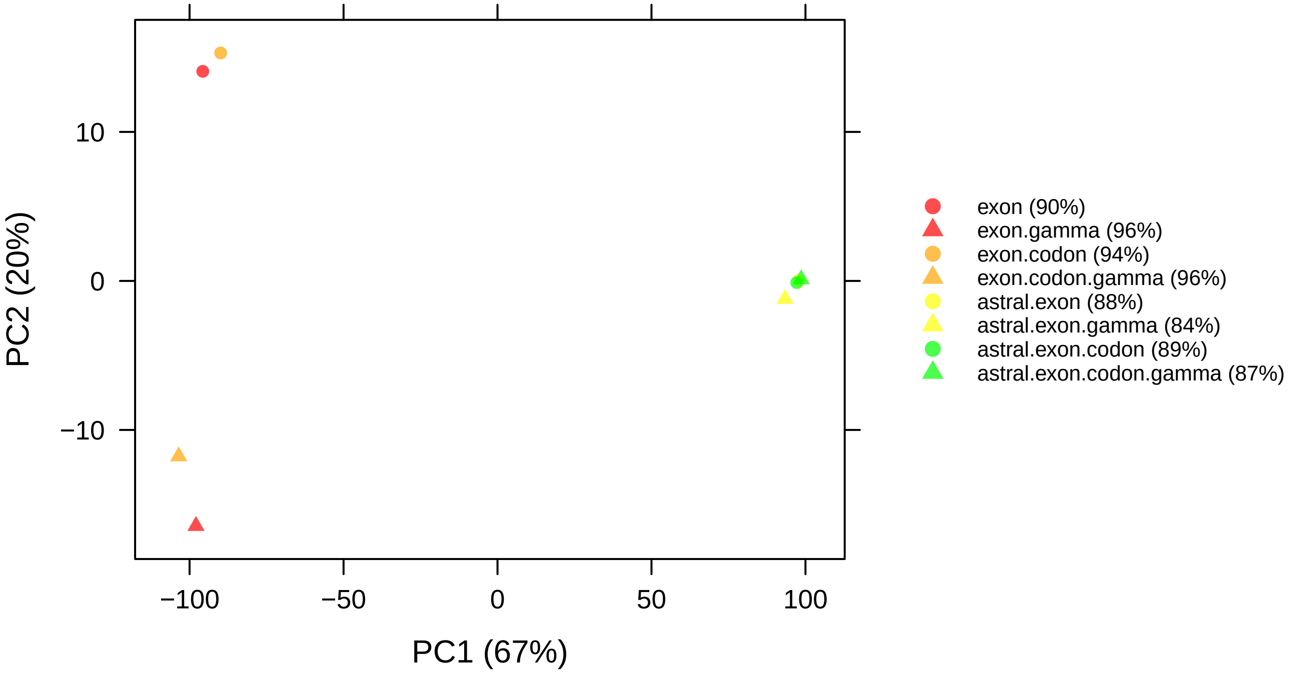

10
0
−10
−100
−50
0
50
100
PC1 (67%)
PC2 (20%)
exon (90%)
exon.gamma (96%)
exon.codon (94%)
exon.codon.gamma (96%)
astral.exon (88%)
astral.exon.gamma (84%)
astral.exon.codon (89%)
astral.exon.codon.gamma (87%)

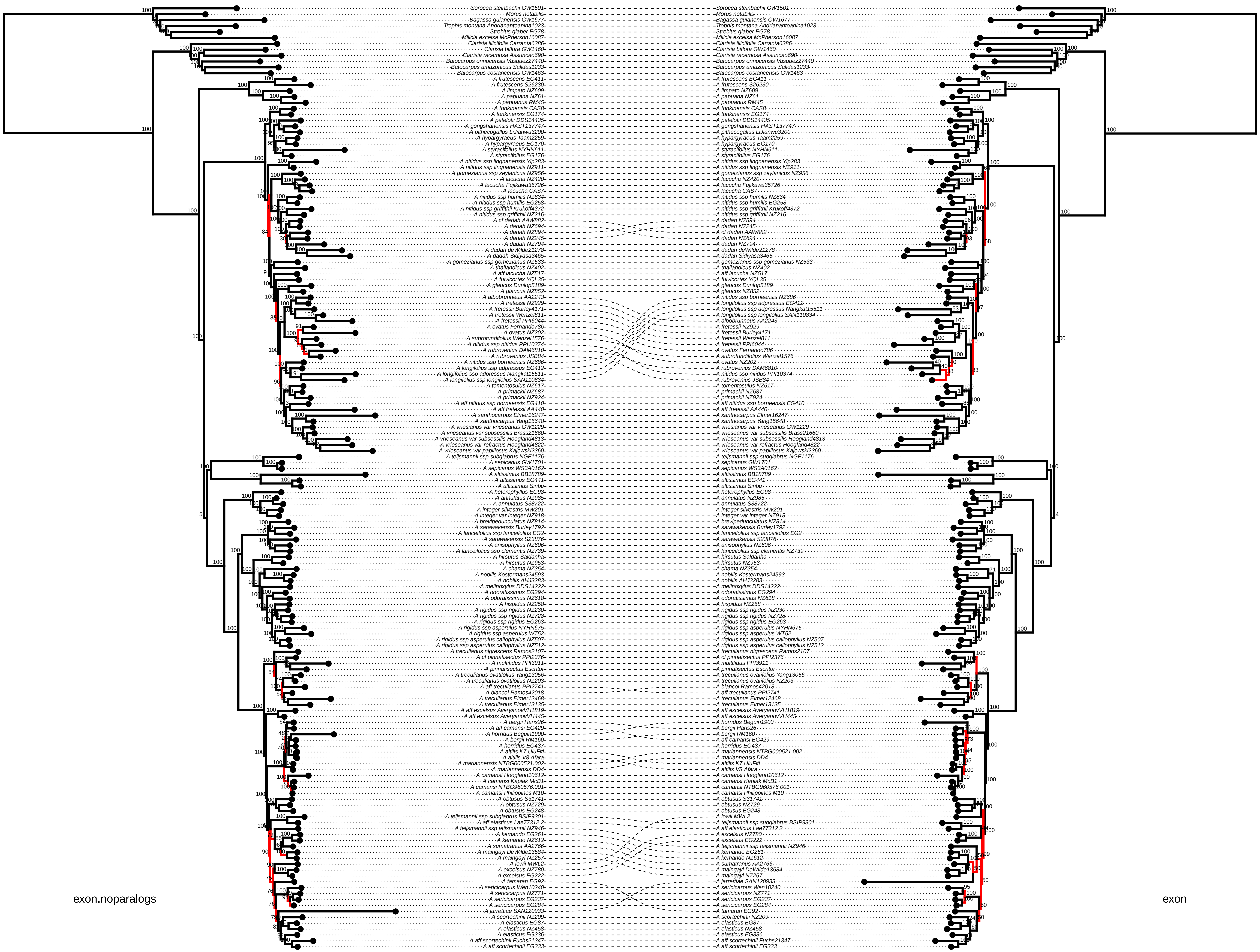
exon.noparalogs
exon
Sorocea steinbachii GW1501
Morus notabilis
Bagassa guianensis GW1677
Trophis montana Andrianantoanina1023
Streblus glaber EG78
Milicia excelsa McPherson16087
Clarisia illicifolia Carranta6386
Clarisia biflora GW1460
Clarisia racemosa Assuncao690
Batocarpus orinocensis Vasquez27440
Batocarpus amazonicus Salidas1233
Batocarpus costaricensis GW1463
A frutescens EG411
A frutescens S26230
A limpato NZ609
A papuanus NZ61
A papuanus RM45
A tonkinensis CAS8
A tonkinensis EG174
A petelotii DDS14435
A gongshanensis HAST137747
A pithecogallus LiJianwu3200
A hypargyraeus Taam2259
A hypargyraeus EG170
A styracifolius NYHN611
A styracifolius EG176
A nitidus ssp lingnanensis Yip283
A nitidus ssp lingnanensis NZ911
A gomezianus ssp zeylanicus NZ956
A lacucha NZ420
A lacucha Fujikawa35726
A lacucha CAS7
A nitidus ssp humilis NZ834
A nitidus ssp humilis EG258
A nitidus ssp griffithii Krukoff4372
A nitidus ssp griffithii NZ216
A cf dadah AAW882
A dadah NZ694
A dadah NZ894
A dadah NZ245
A dadah NZ794
A dadah deWilde21278
A dadah Sidiyasa3465
A gomezianus ssp gomezianus NZ533
A thailandicus NZ402
A aff lacucha NZ517
A fulvicortex YQL35
A glaucus Dunlop5189
A glaucus NZ852
A albobrunneus AA2243
A fretessii NZ929
A fretessii Burley4171
A fretessii Wenzel811
A fretessii PPI6044
A ovatus Fernando786
A ovatus NZ202
A subrotundifolius Wenzel1576
A nitidus ssp nitidus PPI10374
A rubrovenius DAM6810
A rubrovenius JSB84
A nitidus ssp borneensis NZ686
A longifolius ssp adpressus EG412
A longifolius ssp adpressus Nangkat15511
A longifolius ssp longifolius SAN110834
A tomentosulus NZ617
A primackii NZ687
A primackii NZ924
A aff nitidus ssp borneensis EG410
A aff fretessii AA440
A xanthocarpus Elmer16247
A xanthocarpus Yang15648
A vriesianus var vrieseanus GW1229
A vrieseanus var subsessilis Brass21660
A vrieseanus var subsessilis Hoogland4813
A vrieseanus var refractus Hoogland4822
A vrieseanus var papillosus Kajewski2360
A teijsmannii ssp subglabrus NGF1176
A sepicanus GW1701
A sepicanus WS3A0162
A altissimus BB18789
A altissimus EG441
A altissimus Sinbu
A heterophyllus EG98
A annulatus NZ985
A annulatus S38722
A integer silvestris MW201
A integer var integer NZ918
A brevipedunculatus NZ814
A sarawakensis Burley1792
A lanceifolius ssp lanceifolius EG2
A sarawakensis S23876
A anisophyllus NZ606
A lanceifolius ssp clementis NZ739
A hirsutus Saldanha
A hirsutus NZ953
A chama NZ354
A nobilis Kostermans24593
A nobilis AHJ3283
A melinoxylus DDS14222
A odoratissimus EG294
A odoratissimus NZ618
A hispidus NZ258
A rigidus ssp rigidus NZ230
A rigidus ssp rigidus NZ728
A rigidus ssp rigidus EG263
A rigidus ssp asperulus NYHN675
A rigidus ssp asperulus WT52
A rigidus ssp asperulus callophyllus NZ507
A rigidus ssp asperulus callophyllus NZ512
A treculianus nigrescens Ramos2107
A cf pinnatisectus PPI2376
A multifidus PPI3911
A pinnatisectus Escritor
A treculianus ovatifolius Yang13056
A treculianus ovatifolius NZ203
A aff treculianus PPI2741
A blancoi Ramos42018
A treculianus Elmer12468
A treculianus Elmer13135
A aff excelsus AveryanovVH1819
A aff excelsus AveryanovVH445
A bergii Haris26
A aff camansi EG429
A horridus Beguin1900
A bergii RM160
A horridus EG437
A altilis K7 UluFiti
A altilis V8 Afara
A mariannensis NTBG000521.002
A mariannensis DD4
A camansi Hoogland10612
A camansi Kapiak McB1
A camansi NTBG960576.001
A camansi Philippines M10
A obtusus S31741
A obtusus NZ729
A obtusus EG248
A teijsmannii ssp subglabrus BSIP9301
A aff elasticus Lae77312 2
A teijsmannii ssp teijsmannii NZ946
A kemando EG261
A kemando NZ612
A sumatranus AA2766
A maingayi DeWilde13584
A maingayi NZ257
A lowii MWL2
A excelsus NZ780
A excelsus EG222
A tamaran EG92
A sericicarpus Wen10240
A sericicarpus NZ771
A sericicarpus EG237
A sericicarpus EG284
A jarrettiae SAN120933
A scortechinii NZ209
A elasticus EG87
A elasticus NZ458
A elasticus EG336
A aff scortechinii Fuchs21347
A aff scortechinii EG333

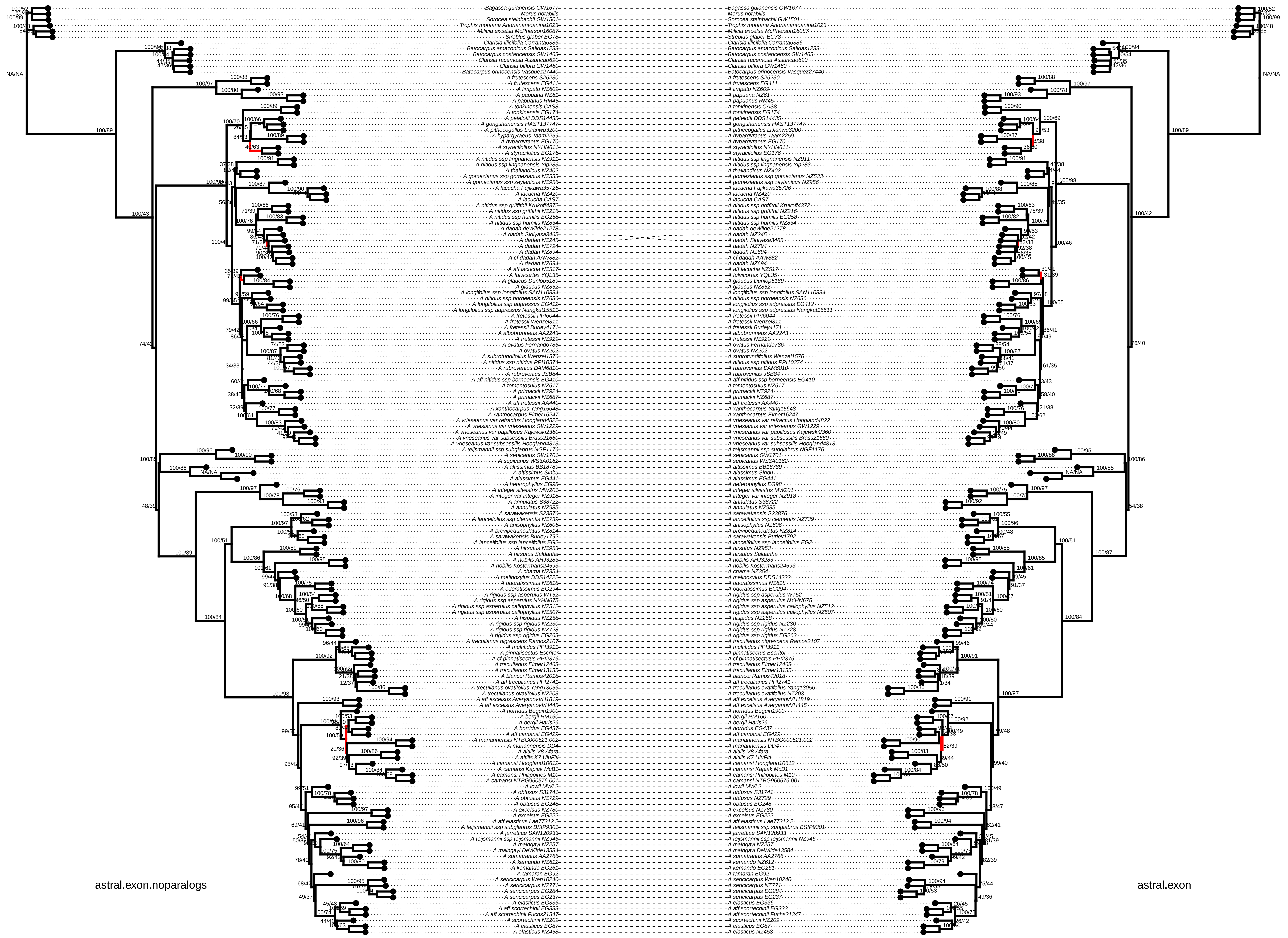
astral.exon.noparalogs
astral.exon

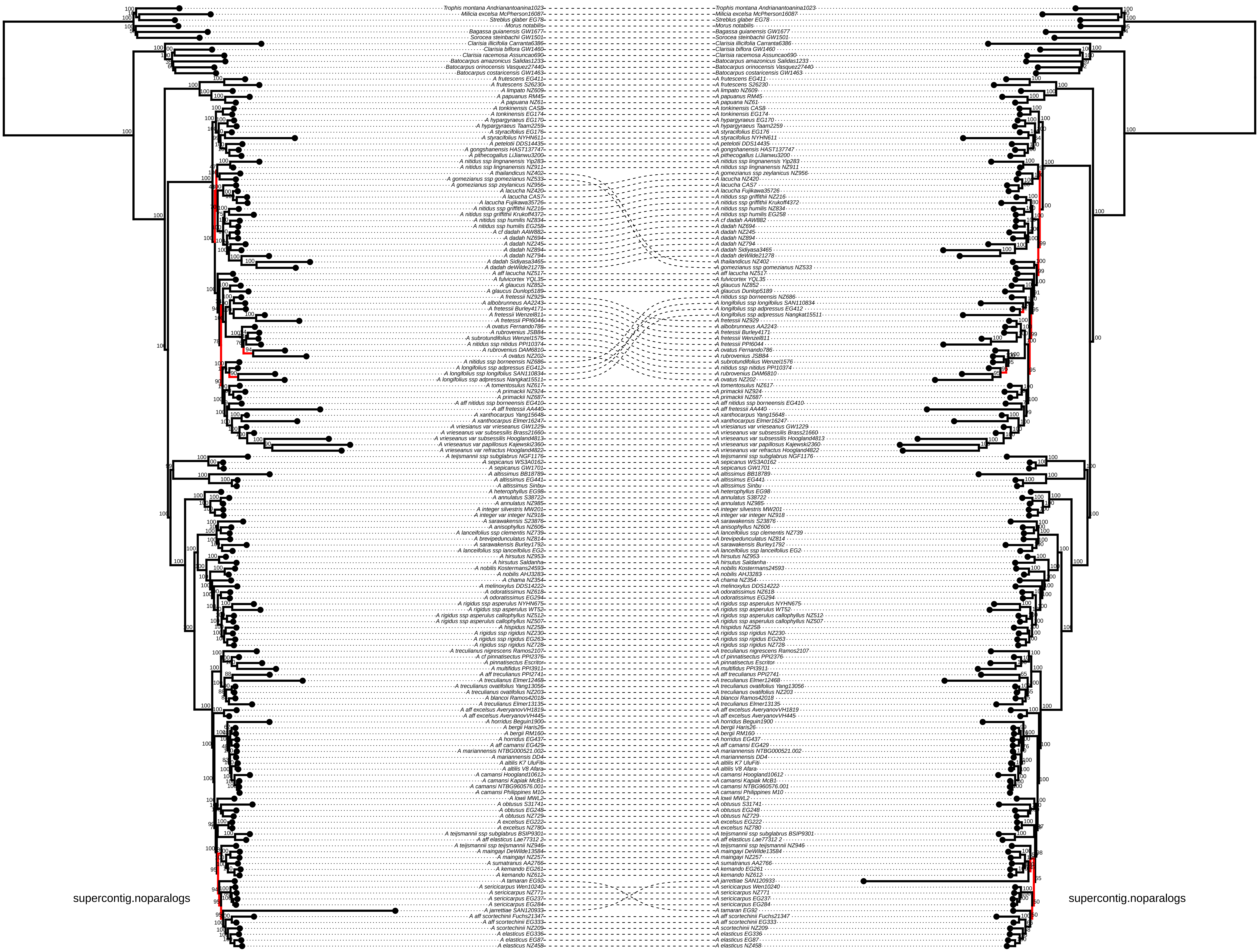
supercontig.noparalogs
supercontig.noparalogs

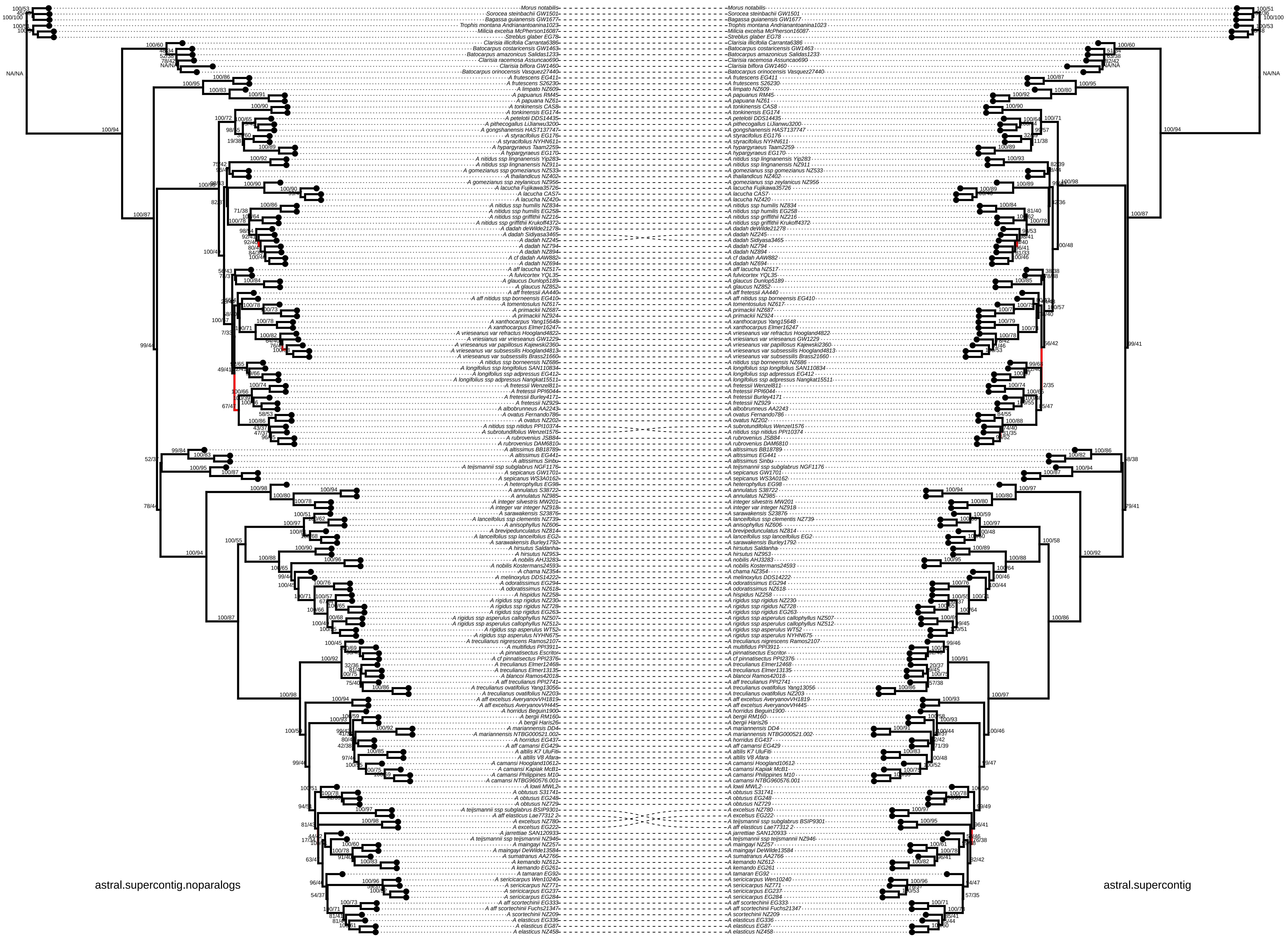
astral.supercontig.noparalogs
astral.supercontig

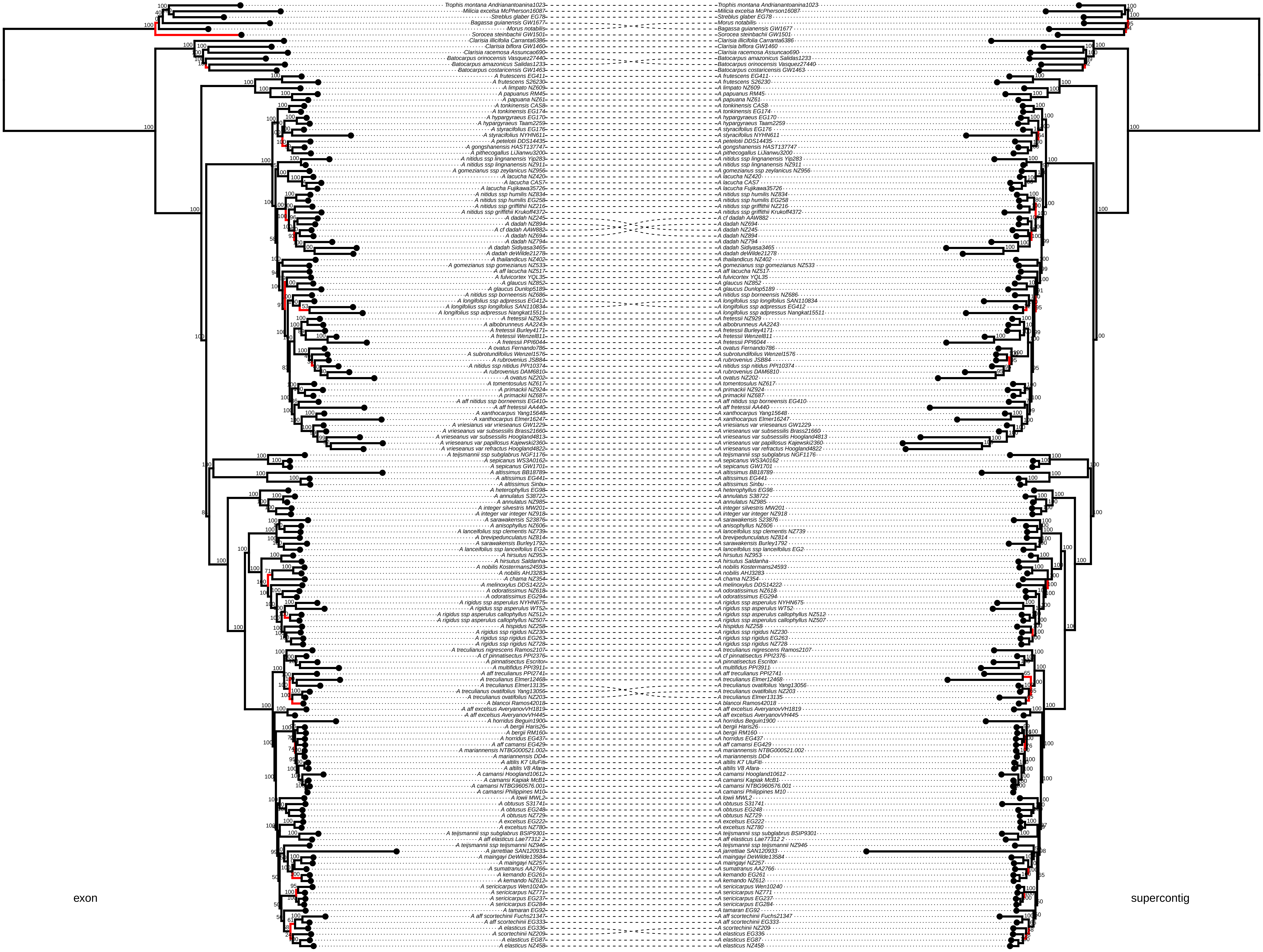
exon
supercontig
Trophis montana Andrianantoanina1023
Milicia excelsa McPherson16087
Streblus glaber EG78
Bagassa guianensis GW1677
Morus notabilis
Sorocea steinbachii GW1501
Clarisia illicifolia Carranta6386
Clarisia biflora GW1460
Clarisia racemosa Assuncao690
Batocarpus orinocensis Vasquez27440
Batocarpus amazonicus Salidas1233
Batocarpus costaricensis GW1463
A frutescens EG411
A frutescens S26230
A limpato NZ609
A papuanus RM45
A papuana NZ61
A tonkinensis CAS8
A tonkinensis EG174
A hypargyraeus EG170
A hypargyraeus Taam2259
A styracifolius EG176
A styracifolius NYHN611
A petelotii DDS14435
A gongshanensis HAST137747
A pithecogallus LiJianwu3200
A nitidus ssp lingnanensis Yip283
A nitidus ssp lingnanensis NZ911
A gomezianus ssp zeylanicus NZ956
A lacucha NZ420
A lacucha CAS7
A lacucha Fujikawa35726
A nitidus ssp humilis NZ834
A nitidus ssp humilis EG258
A nitidus ssp griffithii NZ216
A nitidus ssp griffithii Krukoff4372
A dadah NZ245
A dadah NZ894
A cf dadah AAW882
A dadah NZ694
A dadah NZ794
A dadah Sidiyasa3465
A dadah deWilde21278
A thailandicus NZ402
A gomezianus ssp gomezianus NZ533
A aff lacucha NZ517
A fulvicortex YQL35
A glaucus NZ852
A glaucus Dunlop5189
A nitidus ssp borneensis NZ686
A longifolius ssp adpressus EG412
A longifolius ssp longifolius SAN110834
A longifolius ssp adpressus Nangkat15511
A fretessii NZ929
A albobrunneus AA2243
A fretessii Burley4171
A fretessii Wenzel811
A fretessii PPI6044
A ovatus Fernando786
A subrotundifolius Wenzel1576
A rubrovenius JSB84
A nitidus ssp nitidus PPI10374
A rubrovenius DAM6810
A ovatus NZ202
A tomentosulus NZ617
A primackii NZ924
A primackii NZ687
A aff nitidus ssp borneensis EG410
A aff fretessii AA440
A xanthocarpus Yang15648
A xanthocarpus Elmer16247
A vriesianus var vrieseanus GW1229
A vrieseanus var subsessilis Brass21660
A vrieseanus var subsessilis Hoogland4813
A vrieseanus var papillosus Kajewski2360
A vrieseanus var refractus Hoogland4822
A teijsmannii ssp subglabrus NGF1176
A sepicanus WS3A0162
A sepicanus GW1701
A altissimus BB18789
A altissimus EG441
A altissimus Sinbu
A heterophyllus EG98
A annulatus S38722
A annulatus NZ985
A integer silvestris MW201
A integer var integer NZ918
A sarawakensis S23876
A anisophyllus NZ606
A lanceifolius ssp clementis NZ739
A brevipedunculatus NZ814
A sarawakensis Burley1792
A lanceifolius ssp lanceifolius EG2
A hirsutus NZ953
A hirsutus Saldanha
A nobilis Kostermans24593
A nobilis AHJ3283
A chama NZ354
A melinoxylus DDS14222
A odoratissimus NZ618
A odoratissimus EG294
A rigidus ssp asperulus NYHN675
A rigidus ssp asperulus WT52
A rigidus ssp asperulus callophyllus NZ512
A rigidus ssp asperulus callophyllus NZ507
A hispidus NZ258
A rigidus ssp rigidus NZ230
A rigidus ssp rigidus EG263
A rigidus ssp rigidus NZ728
A treculianus nigrescens Ramos2107
A cf pinnatisectus PPI2376
A pinnatisectus Escritor
A multifidus PPI3911
A aff treculianus PPI2741
A treculianus Elmer12468
A treculianus Elmer13135
A treculianus ovatifolius Yang13056
A treculianus ovatifolius NZ203
A blancoi Ramos42018
A aff excelsus AveryanovVH1819
A aff excelsus AveryanovVH445
A horridus Beguin1900
A bergii Haris26
A bergii RM160
A horridus EG437
A aff camansi EG429
A mariannensis NTBG000521.002
A mariannensis DD4
A altilis K7 UluFiti
A altilis V8 Afara
A camansi Hoogland10612
A camansi Kapiak McB1
A camansi NTBG960576.001
A camansi Philippines M10
A lowii MWL2
A obtusus S31741
A obtusus EG248
A obtusus NZ729
A excelsus EG222
A excelsus NZ780
A teijsmannii ssp subglabrus BSIP9301
A aff elasticus Lae77312 2
A teijsmannii ssp teijsmannii NZ946
A jarrettiae SAN120933
A maingayi DeWilde13584
A maingayi NZ257
A sumatranus AA2766
A kemando EG261
A kemando NZ612
A sericicarpus Wen10240
A sericicarpus NZ771
A sericicarpus EG237
A sericicarpus EG284
A tamaran EG92
A aff scortechinii Fuchs21347
A aff scortechinii EG333
A elasticus EG336
A scortechinii NZ209
A elasticus EG87
A elasticus NZ458

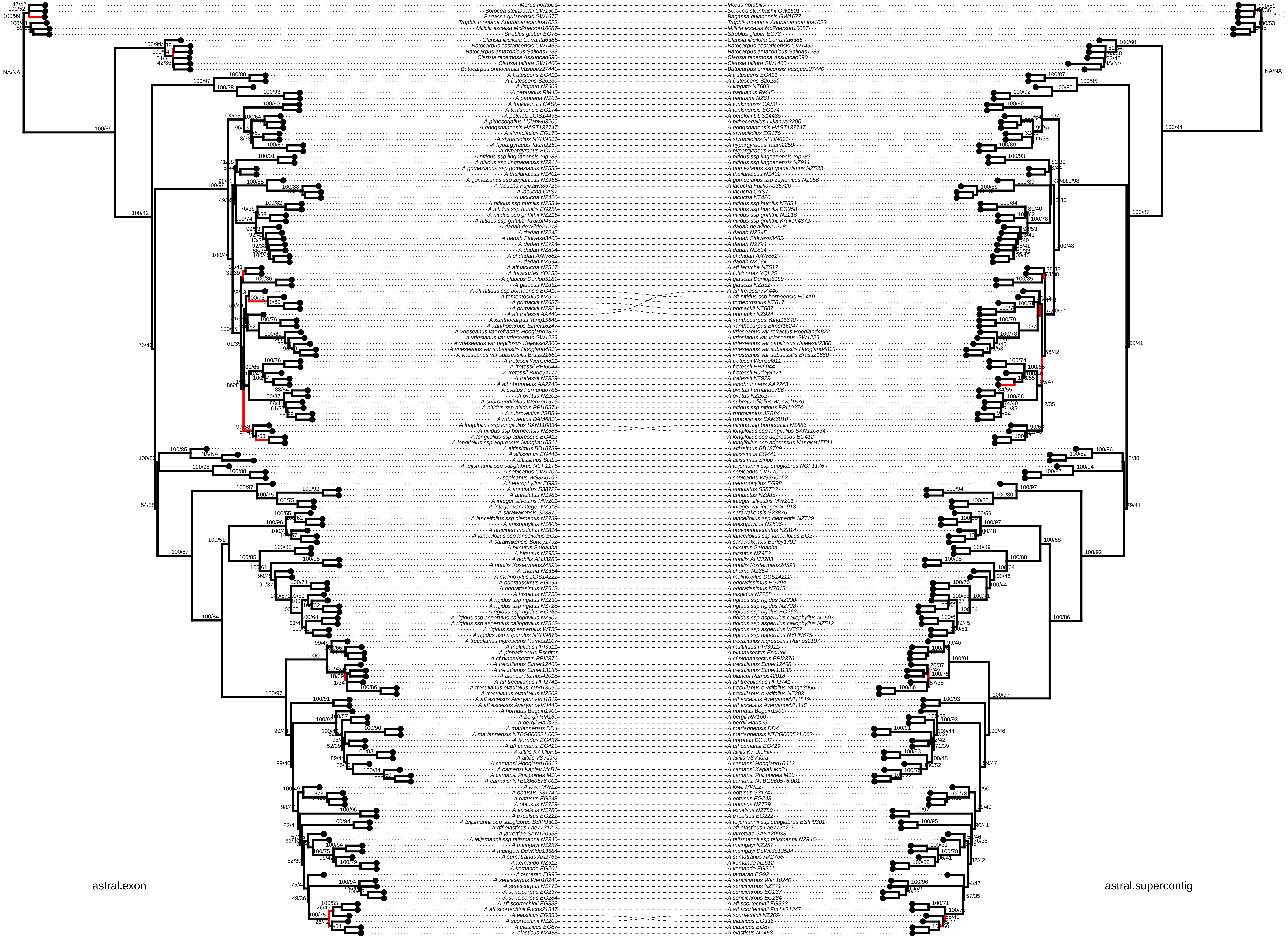
astral.exon
astral.supercontig
Morus notabilis
Sorocea steinbachii GW1501
Bagassa guianensis GW1677
Trophis montana Andrianantoanina1023
Milicia excelsa McPherson16087
Streblus glaber EG78
Clarisia illicifolia Carranta6386
Batocarpus costaricensis GW1463
Batocarpus amazonicus Salidas1233
Clarisia racemosa Assuncao690
Clarisia biflora GW1460
Batocarpus orinocensis Vasquez27440
A frutescens EG411
A frutescens S26230
A limpato NZ609
A papuanus RM45
A papuana NZ61
A tonkinensis CAS8
A tonkinensis EG174
A petelotii DDS14435
A pithecogallus LiJianwu3200
A gongshanensis HAST137747
A styracifolius EG176
A styracifolius NYHN611
A hypargyraeus Taam2259
A hypargyraeus EG170
A nitidus ssp lingnanensis Yip283
A nitidus ssp lingnanensis NZ911
A gomezianus ssp gomezianus NZ533
A thailandicus NZ402
A gomezianus ssp zeylanicus NZ956
A lacucha Fujikawa35726
A lacucha CAS7
A lacucha NZ420
A nitidus ssp humilis NZ834
A nitidus ssp humilis EG258
A nitidus ssp griffithii NZ216
A nitidus ssp griffithii Krukoff4372
A dadah deWilde21278
A dadah NZ245
A dadah Sidiyasa3465
A dadah NZ794
A dadah NZ894
A cf dadah AAW882
A dadah NZ694
A aff lacucha NZ517
A fulvicortex YQL35
A glaucus Dunlop5189
A glaucus NZ852
A aff nitidus ssp borneensis EG410
A tomentosulus NZ617
A primackii NZ687
A primackii NZ924
A aff fretessii AA440
A xanthocarpus Yang15648
A xanthocarpus Elmer16247
A vrieseanus var refractus Hoogland4822
A vriesianus var vrieseanus GW1229
A vrieseanus var papillosus Kajewski2360
A vrieseanus var subsessilis Hoogland4813
A vrieseanus var subsessilis Brass21660
A fretessii Wenzel811
A fretessii PPI6044
A fretessii Burley4171
A fretessii NZ929
A albobrunneus AA2243
A ovatus Fernando786
A ovatus NZ202
A subrotundifolius Wenzel1576
A nitidus ssp nitidus PPI10374
A rubrovenius JSB84
A rubrovenius DAM6810
A longifolius ssp longifolius SAN110834
A nitidus ssp borneensis NZ686
A longifolius ssp adpressus EG412
A longifolius ssp adpressus Nangkat15511
A altissimus BB18789
A altissimus EG441
A altissimus Sinbu
A teijsmannii ssp subglabrus NGF1176
A sepicanus GW1701
A sepicanus WS3A0162
A heterophyllus EG98
A annulatus S38722
A annulatus NZ985
A integer silvestris MW201
A integer var integer NZ918
A sarawakensis S23876
A lanceifolius ssp clementis NZ739
A anisophyllus NZ606
A brevipedunculatus NZ814
A lanceifolius ssp lanceifolius EG2
A sarawakensis Burley1792
A hirsutus Saldanha
A hirsutus NZ953
A nobilis AHJ3283
A nobilis Kostermans24593
A chama NZ354
A melinoxylus DDS14222
A odoratissimus EG294
A odoratissimus NZ618
A hispidus NZ258
A rigidus ssp rigidus NZ230
A rigidus ssp rigidus NZ728
A rigidus ssp rigidus EG263
A rigidus ssp asperulus callophyllus NZ507
A rigidus ssp asperulus callophyllus NZ512
A rigidus ssp asperulus WT52
A rigidus ssp asperulus NYHN675
A treculianus nigrescens Ramos2107
A multifidus PPI3911
A pinnatisectus Escritor
A cf pinnatisectus PPI2376
A treculianus Elmer12468
A treculianus Elmer13135
A blancoi Ramos42018
A aff treculianus PPI2741
A treculianus ovatifolius Yang13056
A treculianus ovatifolius NZ203
A aff excelsus AveryanovVH1819
A aff excelsus AveryanovVH445
A horridus Beguin1900
A bergii RM160
A bergii Haris26
A mariannensis DD4
A mariannensis NTBG000521.002
A horridus EG437
A aff camansi EG429
A altilis K7 UluFiti
A altilis V8 Afara
A camansi Hoogland10612
A camansi Kapiak McB1
A camansi Philippines M10
A camansi NTBG960576.001
A lowii MWL2
A obtusus S31741
A obtusus EG248
A obtusus NZ729
A excelsus NZ780
A excelsus EG222
A teijsmannii ssp subglabrus BSIP9301
A aff elasticus Lae77312 2
A jarrettiae SAN120933
A teijsmannii ssp teijsmannii NZ946
A maingayi NZ257
A maingayi DeWilde13584
A sumatranus AA2766
A kemando NZ612
A kemando EG261
A tamaran EG92
A sericicarpus Wen10240
A sericicarpus NZ771
A sericicarpus EG237
A sericicarpus EG284
A aff scortechinii EG333
A aff scortechinii Fuchs21347
A elasticus EG336
A scortechinii NZ209
A elasticus EG87
A elasticus NZ458

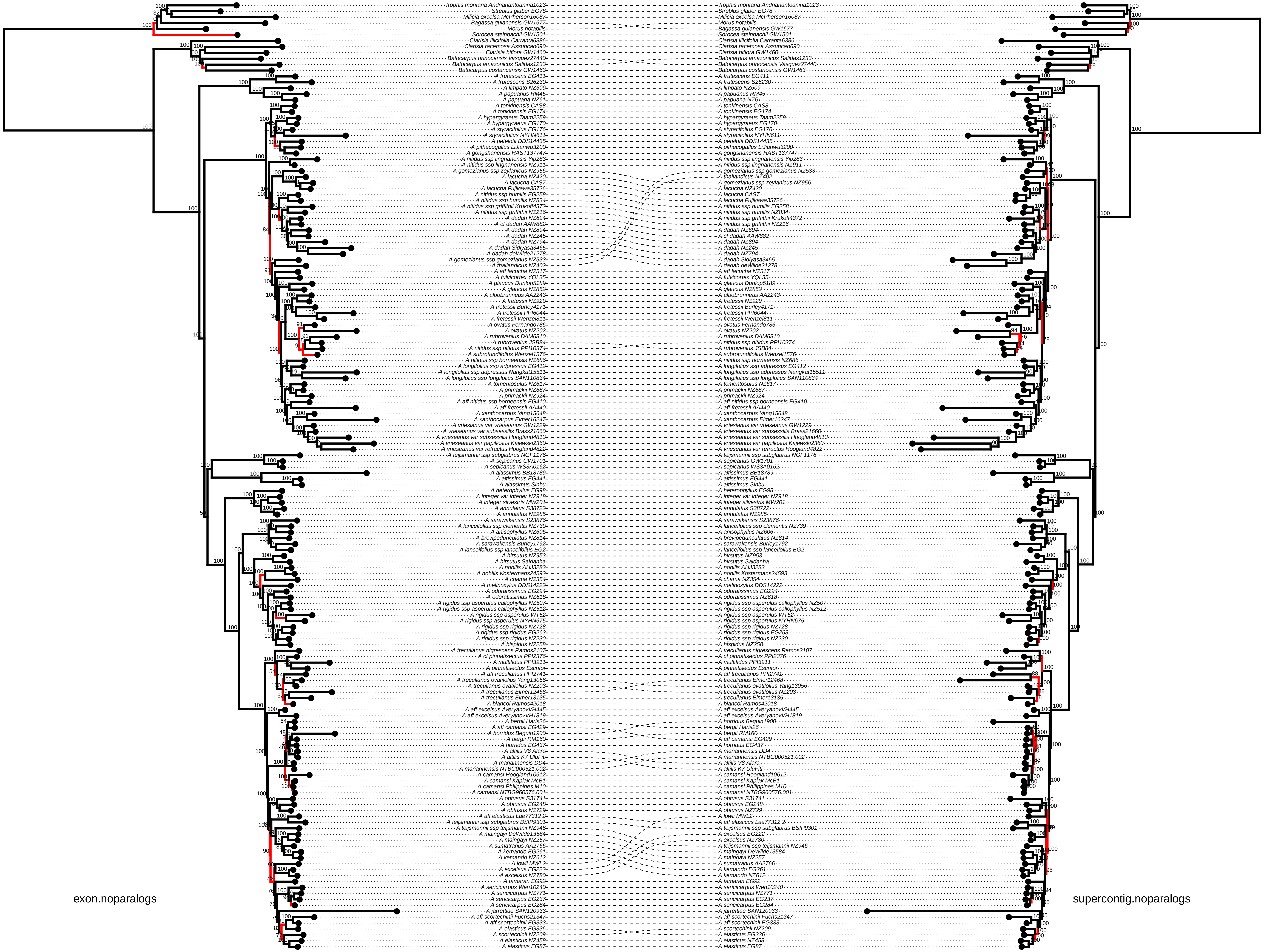
exon.noparalogs
supercontig.noparalogs

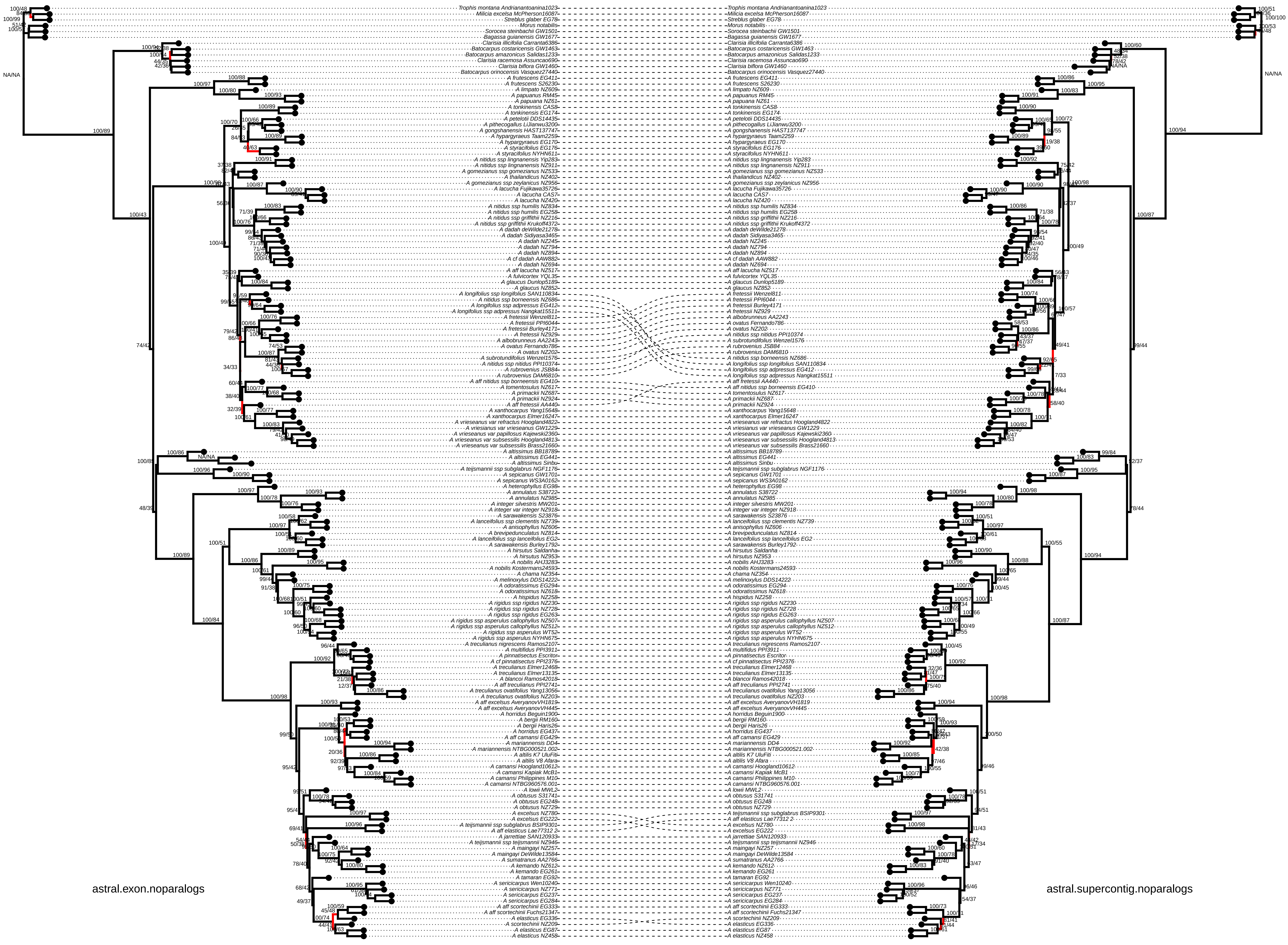
astral.exon.noparalogs
astral.supercontig.noparalogs

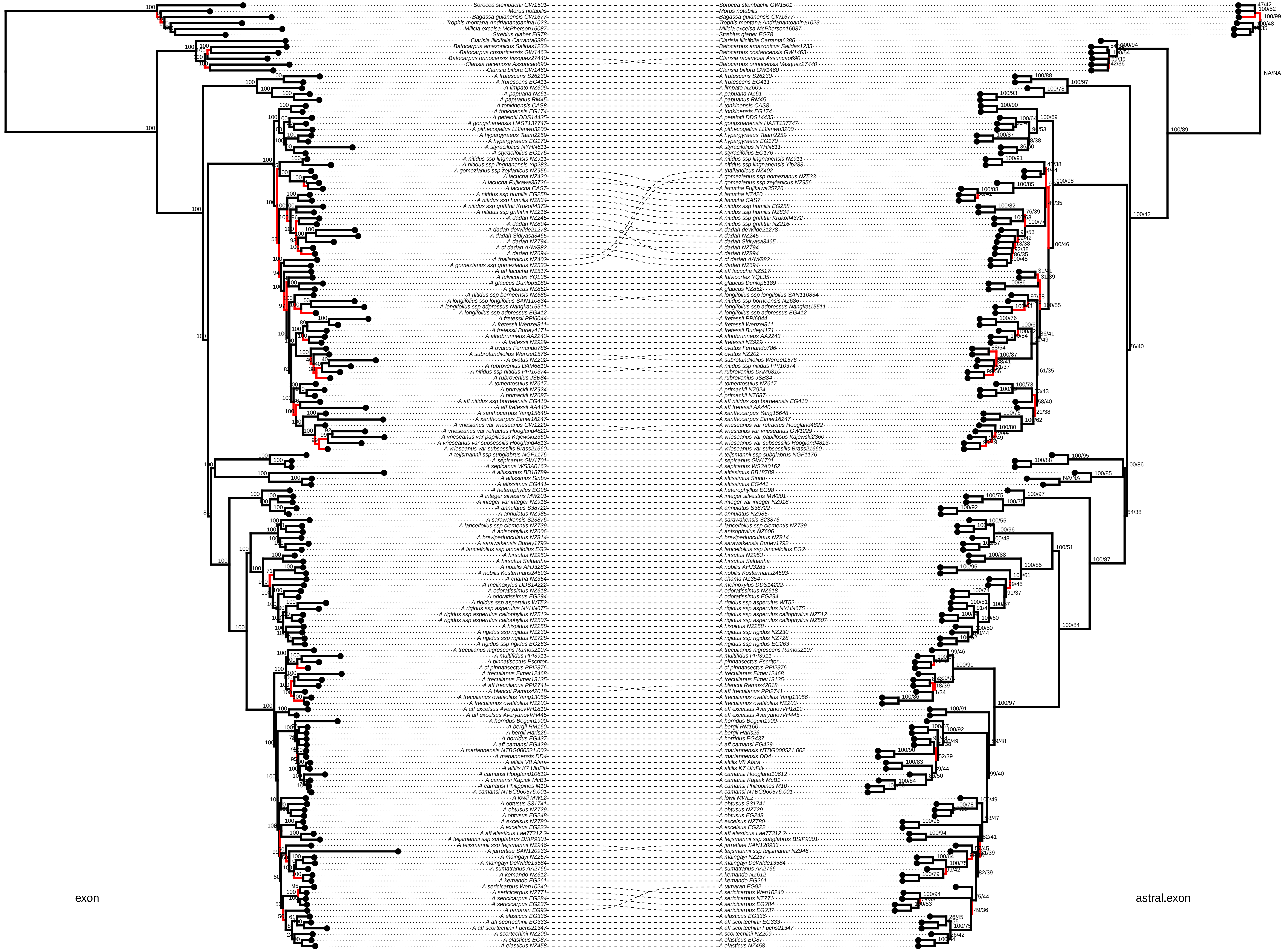
exon
astral.exon
Sorocea steinbachii GW1501
Morus notabilis
Bagassa guianensis GW1677
Trophis montana Andrianantoanina1023
Milicia excelsa McPherson16087
Streblus glaber EG78
Clarisia illicifolia Carranta6386
Batocarpus amazonicus Salidas1233
Batocarpus costaricensis GW1463
Batocarpus orinocensis Vasquez27440
Clarisia racemosa Assuncao690
Clarisia biflora GW1460
A frutescens S26230
A frutescens EG411
A limpato NZ609
A papuana NZ61
A papuanus RM45
A tonkinensis CAS8
A tonkinensis EG174
A petelotii DDS14435
A gongshanensis HAST137747
A pithecogallus LiJianwu3200
A hypargyraeus Taam2259
A hypargyraeus EG170
A styracifolius NYHN611
A styracifolius EG176
A nitidus ssp lingnanensis NZ911
A nitidus ssp lingnanensis Yip283
A gomezianus ssp zeylanicus NZ956
A lacucha NZ420
A lacucha Fujikawa35726
A lacucha CAS7
A nitidus ssp humilis EG258
A nitidus ssp humilis NZ834
A nitidus ssp griffithii Krukoff4372
A nitidus ssp griffithii NZ216
A dadah NZ245
A dadah NZ894
A dadah deWilde21278
A dadah Sidiyasa3465
A dadah NZ794
A cf dadah AAW882
A dadah NZ694
A thailandicus NZ402
A gomezianus ssp gomezianus NZ533
A aff lacucha NZ517
A fulvicortex YQL35
A glaucus Dunlop5189
A glaucus NZ852
A nitidus ssp borneensis NZ686
A longifolius ssp longifolius SAN110834
A longifolius ssp adpressus Nangkat15511
A longifolius ssp adpressus EG412
A fretessii PPI6044
A fretessii Wenzel811
A fretessii Burley4171
A albobrunneus AA2243
A fretessii NZ929
A ovatus Fernando786
A subrotundifolius Wenzel1576
A ovatus NZ202
A rubrovenius DAM6810
A nitidus ssp nitidus PPI10374
A rubrovenius JSB84
A tomentosulus NZ617
A primackii NZ924
A primackii NZ687
A aff nitidus ssp borneensis EG410
A aff fretessii AA440
A xanthocarpus Yang15648
A xanthocarpus Elmer16247
A vrieseanus var vrieseanus GW1229
A vrieseanus var refractus Hoogland4822
A vrieseanus var papillosus Kajewski2360
A vrieseanus var subsessilis Hoogland4813
A vrieseanus var subsessilis Brass21660
A teijsmannii ssp subglabrus NGF1176
A sepicanus GW1701
A sepicanus WS3A0162
A altissimus BB18789
A altissimus Sinbu
A altissimus EG441
A heterophyllus EG98
A integer silvestris MW201
A integer var integer NZ918
A annulatus S38722
A annulatus NZ985
A sarawakensis S23876
A lanceifolius ssp clementis NZ739
A anisophyllus NZ606
A brevipedunculatus NZ814
A sarawakensis Burley1792
A lanceifolius ssp lanceifolius EG2
A hirsutus NZ953
A hirsutus Saldanha
A nobilis AHJ3283
A nobilis Kostermans24593
A chama NZ354
A melinoxylus DDS14222
A odoratissimus NZ618
A odoratissimus EG294
A rigidus ssp asperulus WT52
A rigidus ssp asperulus NYHN675
A rigidus ssp asperulus callophyllus NZ512
A rigidus ssp asperulus callophyllus NZ507
A hispidus NZ258
A rigidus ssp rigidus NZ230
A rigidus ssp rigidus NZ728
A rigidus ssp rigidus EG263
A treculianus nigrescens Ramos2107
A multifidus PPI3911
A pinnatisectus Escritor
A cf pinnatisectus PPI2376
A treculianus Elmer12468
A treculianus Elmer13135
A aff treculianus PPI2741
A blancoi Ramos42018
A treculianus ovatifolius Yang13056
A treculianus ovatifolius NZ203
A aff excelsus AveryanovVH1819
A aff excelsus AveryanovVH445
A horridus Beguin1900
A bergii RM160
A bergii Haris26
A horridus EG437
A aff camansi EG429
A mariannensis NTBG000521.002
A mariannensis DD4
A altilis V8 Afara
A altilis K7 UluFiti
A camansi Hoogland10612
A camansi Kapiak McB1
A camansi Philippines M10
A camansi NTBG960576.001
A lowii MWL2
A obtusus S31741
A obtusus NZ729
A obtusus EG248
A excelsus NZ780
A excelsus EG222
A aff elasticus Lae77312 2
A teijsmannii ssp subglabrus BSIP9301
A teijsmannii ssp teijsmannii NZ946
A jarrettiae SAN120933
A maingayi NZ257
A maingayi DeWilde13584
A sumatranus AA2766
A kemando NZ612
A kemando EG261
A sericicarpus Wen10240
A sericicarpus NZ771
A sericicarpus EG284
A sericicarpus EG237
A tamaran EG92
A elasticus EG336
A aff scortechinii EG333
A aff scortechinii Fuchs21347
A scortechinii NZ209
A elasticus EG87
A elasticus NZ458

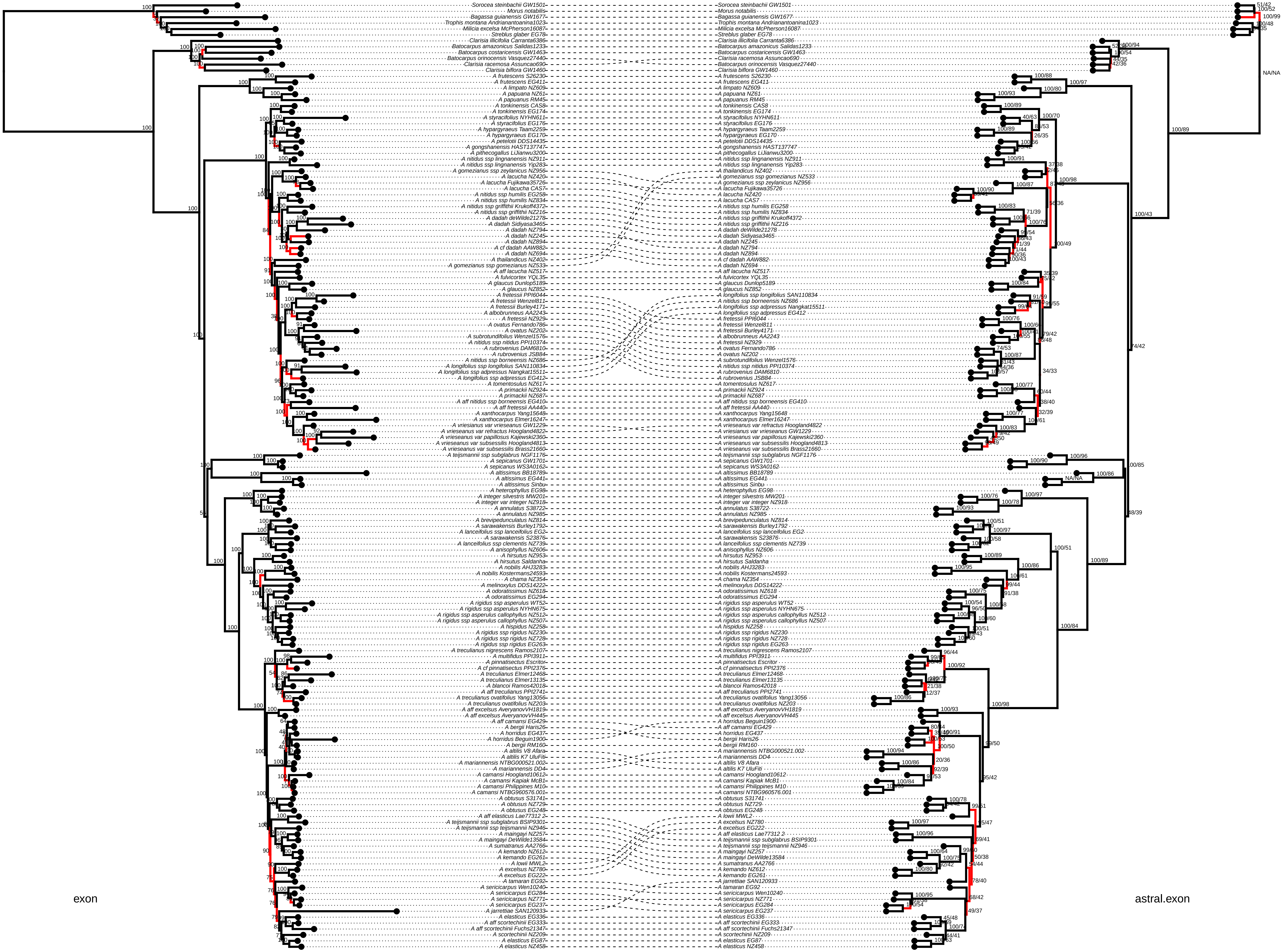
exon
astral.exon

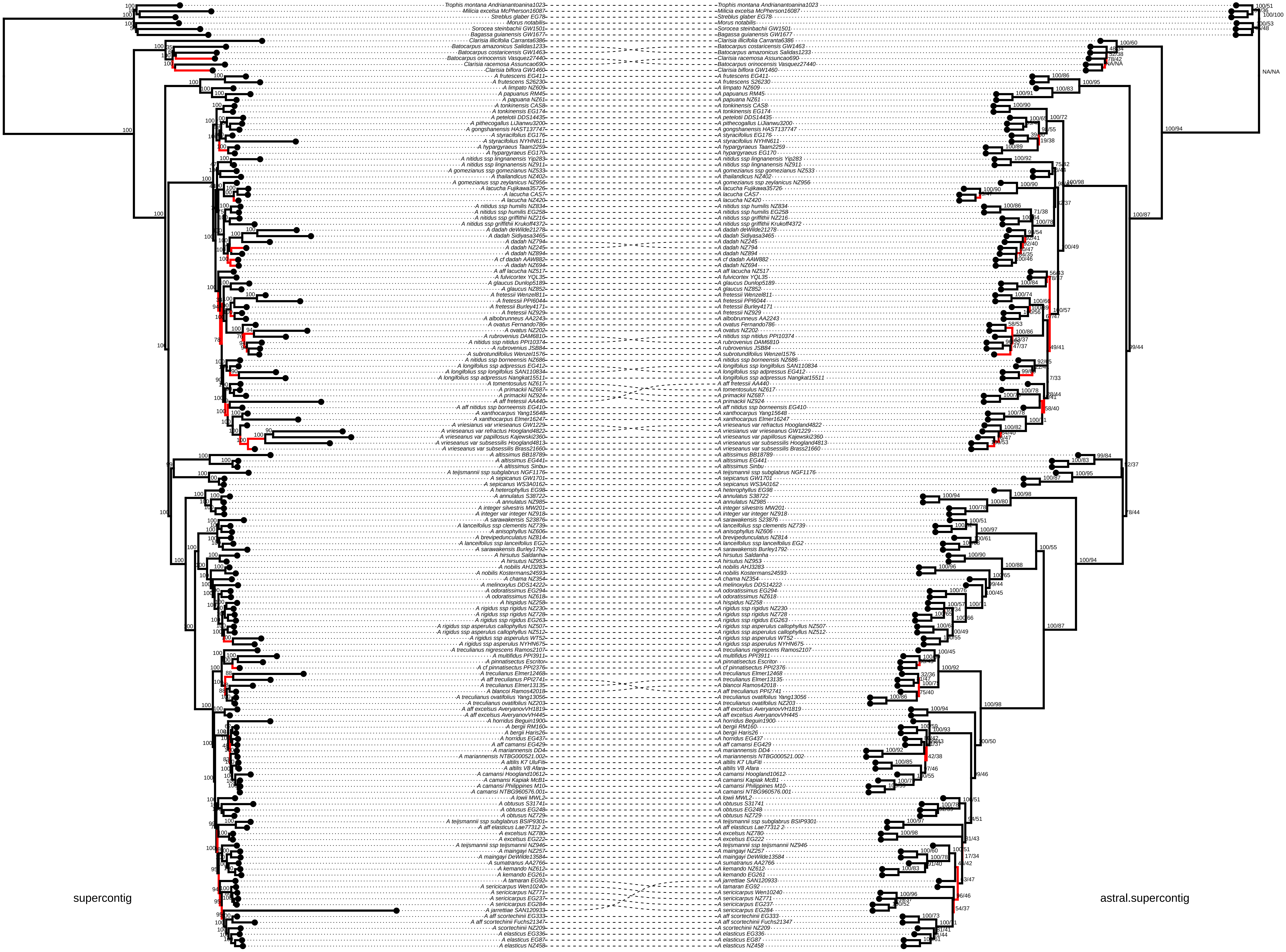
supercontig
astral.supercontig
