## Appendix 1 and 2 for "Paralogs and off-target sequences improve phylogenetic resolution in a densely-sampled study of the breadfruit genus (*Artocarpus*, Moraceae)"

### APPENDIX 1 – METHODS IN DETAIL

#### *Taxon sampling*

We sampled all *Artocarpus* taxa at the subspecies level or above recognized by Jarrett (1959, 1960), Berg et al. (2006), and Kochummen (1998), all three obsolete species that Jarrett (1959) sunk into *A. treculianus* Elmer, and all of the new species described by Wu and Zhang (1989), for a total of 83 named *Artocarpus* taxa. We also sampled nine taxa of questionable affinities. We replicated samples across geographic or morphological ranges when possible, for a total of 167 ingroup samples. As outgroups, we sampled one member of each genus in the Neotropical Artocarpeae (*Batocarpus* and *Clarisia*) and the sister tribe Moreae (*Morus* L., *Streblus* Lour., *Milicia* Sim., *Trophis* P. Browne, *Bagassa* Aubl., and *Sorocea* A. St.-Hil.). We obtained samples from our own field collections preserved in silica gel (from Malaysia, Thailand, Hong Kong, Bangladesh, and India, and from botanic gardens in Indonesia, Malaysia, and Hawai'i, USA) and from herbarium specimens up to 106 years old (from the following herbaria: BM, BO, CHIC, E, F, HAST, HK, K, KUN, L, MO, NY, KEP, S, SAN, SNP, US). In total we included 179 samples (Table S1).

#### *Sample preparation and sequencing*

We sampled approximately 0.5 $cm^2$ of dried leaf from each sample for DNA extraction. For herbarium specimens, we sampled from a fragment packet when feasible and when it was clear that the material in the fragment packet originated from the specimen on the sheet (something that cannot always be assumed with very old specimens). DNA was extracted using one of three methods; (1) the Qiagen DNeasy Plant Mini Kit (Qiagen, Valencia, California, USA) following the manufacturer's protocol; (2) the MoBio PowerPlant Pro DNA Kit, (MoBio Laboratories, Carlsbad, California, USA); or (3) a modified CTAB protocol (Doyle and Doyle 1987). For kit extractions, the protocols were modified for herbarium material by extending initial incubation times (Williams et al., 2017) and adding an additional 200 µL of ethanol to the column-binding step. CTAB extractions of herbarium specimens, which often had high but impure DNA yields, were cleaned using a 1:1.8:5 ratio of sample, SPRI beads, and isopropanol, the latter added to prevent the loss of small fragments (Lee 2014). For herbarium specimens, we sometimes combined two or more separate extractions in order to accumulate enough DNA for library preparation. We assessed degradation of DNA from herbarium specimens using either an agarose gel or a High-Sensitivity DNA Assay on a BioAnalyzer 2100 (Agilent) and did not sonicate samples whose average fragment size was less than 500bp. The remaining DNA samples were sonicated to a mean insert size of 550bp using a Covaris M220 (Covaris, Wobum, Massachussetts, USA). Libraries were prepared with either the Illumina TruSeq Nano HT DNA Library Preparation Kit (Illumina, San Diego, California, USA) or the KAPA Hyper Prep DNA Library Kit following the manufacturer's protocol, except that reactions were performed in one-third volumes to save reagent costs. We used 200ng of input DNA when possible; for some samples, input was as low as 10ng. For herbarium samples with degraded DNA, we usually did not perform size selection, unless there were some fragments that were above 550bp. We also diluted the adapters from 15 µM to 7.5 µM, and usually performed only a single SPRI bead cleanup between adapter ligation and PCR amplification. Many of these libraries contained substantial amounts of adapter dimer, so we adjusted the post-PCR SPRI bead cleanup ratio to 0.8x. Libraries were enriched for 333 phylogenetic markers (Gardner et al., 2016) with a MYbaits kit (MYcroarray, Ann Arbor, Michigan, USA) following the MYbaits manufacturer's protocol (version 3). Hybridization took place in pools of 6–24 libraries; within each pool, we used equal amounts of all libraries (20–100ng, as available), and tried to avoid pooling samples with dramatically different phylogenetic distances to the bait sequnces (*Morus* and *Artocarpus*), as closer taxa can out-compete multiplexed distant taxa in hybridization reactions, as we previously found when pooling *Dorstenia* L. and *Parartocarpus* Baill. with *Artocarpus* (Johnson et al. 2016). We reamplified enriched libraries with 14 PCR cycles using the conditions specified in the manufacturer's protocol. In some cases, adapter dimer remained even after hybridization; in those cases, we removed it either using a 0.7x SPRI bead cleanup or, in cases where the library fragments were very short (ca. 200bp, compared to 144bp for the dimer), by size-selecting the final pools to >180bp on a BluePippin size-selector using a 2% agarose gel cassette (Sage Science, Beverley, Massachussetts, USA). Pools of enriched libraries were sequenced on an Illumina MiSeq (600 cycle, version 3 chemistry) alongside samples for other studies in three multiplexed runs each containing 30–99 samples.

*Sequence quality control and analyses*

Demultiplexing and adapter trimming took place automatically through Illumina BaseSpace (basespace.illumina.com). All reads have been deposited in GenBank (BioProject no. PRJNA322184). Raw reads were quality trimmed using Trimmomatic (Bolger et al., 2014), with a quality cutoff of 20 in a 4-bp sliding window, discarding any reads trimmed to under 30 bp. In addition to the samples sequenced for this study, reads used for assemblies included all *Artocarpus* samples sequenced in Johnson et al. (2016) (available under the same BioProject number). Common methods for target capture assembly include mapping reads to a reference (Weitemier et al. 2014; Hart et al. 2016) and *de novo* assemblies (Mandel et al. 2014; Faircloth 2015), but both have drawbacks. Read mapping can result in lost data, particularly indels and non-coding regions, unless a close reference is available. On the other hand, *de novo* assemblies can also result in lost data if loci cannot be assembled into single scaffolds. A compromise approach, implemented in HybPiper, is to combine local *de novo* assemblies—which may result in many small contigs per locus—with scaffolding based on a reference coding sequence, which need not be closely related; a reference with less than 30% sequence, typically within the same family or order, will usually suffice (Johnson et al. 2016, 2019). The resulting assemblies thus cover the maximum available portion of each locus, notwithstanding the existence of long gaps, and also make use of all available on-target reads, including introns, not simply those that can be aligned to a reference.

We assembled sequences using HybPiper 1.2, which represented an update of the original pipeline optimized for short reads from highly-fragmented DNA from museum specimens. HybPiper's guided assembly method uses the reference to scaffold localized de novo assemblies. This is particularly advantageous when dealing with very short reads from degraded DNA, because for those samples, reads covering a single exon may assemble into more than one contig. In those cases, HybPiper uses the reference to scaffold and concatenate multiple contigs into a "supercontig" containing the gene of interest as well as any flanking noncoding sequences (Johnson et al. 2016). The new version of HybPiper is optimized to accurately handle many small contigs covering a single gene, deduplicating overlaps and outputting high-confidence predicted coding sequences even in the presence of many gaps caused by fragmentary local assemblies. HybPiper as well as all related scripts used in this study are available at https://github.com/mossmatters/HybPiper and https://github.com/mossmatters/phyloscripts. We generated a new HybPiper reference for this study, using reads from all four subgenera of

*Artocarpus*. Target-enriched reads from *A. camansi* Blanco (the same individual used for whole-genome sequencing in the original marker development (Gardner et al. 2016), *A. limpato* Miq., *A. heterophyllus,* and *A. lacucha* (the latter three from reads sequenced in Johnson et al. (2016)) were assembled de novo using SPAdes (Bankevich et al. 2012), and genes were predicted using Augustus (Keller et al. 2011), with *Arabidopsis* Hehyn. as the reference. Predicted genes were annotated using a BLASTn search seeded with the HybPiper target file of 333 phylogenetic marker genes from Johnson et al. (2016). Paralogs were annotated as follows: genes covering at least 75% of the primary ortholog (labeled “p0”) were labeled as “paralogs” (“p1”, “p2”, etc.). Genes covering less than 75% of the primary ortholog (labeled “e0”) were labeled as “extras” (“e1”, “e2”, etc.), denoting uncertainty as to whether they are paralogs or merely genes with a shared domain. Single copy genes were labeled as “single” in the new reference. We used this new 4-taxon reference to guide all ingroup assemblies, and we used the original set of *Morus notabilis* targets (Johnson et al., 2016) to guide all outgroup assemblies.

We set the per-gene coverage cutoff to 8x, except for certain low-read samples where gene recovery was improved by lowering the coverage cutoff to 4x (10 samples) or 2x (18 samples). HybPiper relies on SPAdes for local *de novo* assemblies. SPAdes creates several assemblies with different k-mer values, with the maximum estimated from the reads (up to 127bp), and then merges them into a final assembly. For herbarium samples that initially recovered fewer than 400 genes, we reran HybPiper, manually setting the maximum k-mer values for assembly to 55 instead of allowing SPAdes to automatically set it. To extract non-coding sequences and annotate gene features along assembled contigs, we used the HybPiper script “intronerate.py”. We assessed target recovery success using the get_seq_lengths.py and gene_recovery_heatmap.r scripts from HybPiper.

To mask low-coverage regions likely to contain sequencing errors, we mapped each sample’s reads to its HybPiper supercontigs using BWA (Li and Durbin 2009), removed PCR duplicates using Picard (Broad Institute 2016), and calculated the depth at each position with Samtools (Li et al. 2009). Using BedTools (Quinlan and Hall 2010), we then hard-masked all positions covered by less than two unique reads. We then used the masked supercontigs and the HybPiper gene annotation files to generate masked versions of the standard HybPiper outputs (using intron_exon_extractor.py): (1) the predicted coding sequence for each target gene (“exon”); (2) the entire contig assembled for each gene (“supercontig”); and (3) the predicted non-coding sequences for each gene (“non-coding”, including introns, UTRs, and intergenic sequences).

To the HybPiper output, we added the original orthologs (CDS only) identified in *Morus notabilis* (Gardner et al., 2016). Because paralogs were only assembled for ingroup samples (due to an *Artocarpus*-specific whole-genome duplication (Gardner et al. 2016)), we added the corresponding “p0” or “e0” from *Morus* to each paralog alignment to serve as an outgroup.

We filtered each set of sequences as follows. For “exon” sequences, we subtracted masked bases (Ns) and removed sequences less than 150 bp and sequences covering less than 20% of the average sequence length for that gene. For “supercontig” sequences, we removed sequences whose corresponding “exon” sequences had been removed. Samples with less than 100 genes remaining after filtering were excluded from the main analyses.

Alignment and trimming then proceeded as follows. For “exon” output, after removing the genes and sequences identified during the filtering stage, we created in-frame alignments using MACSE (Ranwez 2011). For “supercontig” output, we used MAFFT for alignment (--

maxiter 1000) (Katoh and Standley 2013). We trimmed all alignments to remove all columns with >75% gaps using Trimal (Capella-Gutiérrez et al. 2009).

To quickly inspect gene trees for artifacts, we built gene trees from the trimmed "exon" alignments using FastTree (Price et al. 2009) and visually inspected the gene trees for outlier long branches within the ingroup to identify alignments containing improperly sorted paralogous sequences. In some cases, we visually inspected alignments using AliView (Larsson 2014). We discarded a small number of genes whose alignments contained paralogous sequences, for a final set of 517 genes, including all of the original 333 genes.

We used the trimmed alignments to create three sets of gene alignment datasets:

1. *CDS:* "exon" alignments, not partitioned by codon position;
2. *Partitioned CDS*: 333 "exon" alignments, partitioned by codon position; and
3. *Supercontig:* "supercontig" alignments, not partitioned within genes

We also attempted to create a codon-partitioned supercontig alignment by separately aligning "exon" and "intron" sequences and then concatenating them, resulting in three partitions per gene. However, this dataset differed substantially from the *supercontig* dataset, resulting in substantially differing (and nonsensical) topologies even when the partitions were removed; samples with a high proportion of very short or missing non-coding sequences clustered together, perhaps because aligning very short non-coding sequences without longer coding sequences to anchor them produced unreliable alignments. We therefore did not include the *partitioned exon+intron* dataset in the main analyses (discussed further in Appendix 2).

To investigate whether including both copies of a paralogous locus impacted phylogenetic reconstruction, we created versions of each dataset with and without paralogs. We analyzed each of these six datasets using the following two methods, for a total of 12 analyses: (A) *Concatenated supermatrix:* all genes were concatenated into a supermatrix, with each gene partitioned separately (i.e. 1 or 3 partitions per gene, depending on the dataset) and analyzed using RAxML 10 (Stamatakis, 2006) under GTR+CAT model with 200 rapid bootstrap replicates, rooted with the Moreae outgroups; (B) *Species tree:* each gene alignment was analyzed using RAxML 10 under the GTR+CAT model with 200 rapid bootstrap replicates, rooted with the Moreae outgroups. Nodes with <33% support were collapsed into polytomies using SumTrees (Sukumaran and Holder 2010), and the resulting trees were used to estimate a species tree with ASTRAL-III (Mirarab and Warnow, 2015). We estimated node support with multilocus bootstrapping (-r, 160 bootstrap replicates) and by calculating the proportion of quartet trees that support each node (-t 1) (Mirarab and Warnow 2015; Zhang et al. 2017). For the final trees, we also used SumTrees to calculate the proportion of gene trees supporting each split. Quartet support is directly related to the method ASTRAL uses for estimating species trees—decomposing gene trees into quartets (Mirarab and Warnow 2015); it is also less sensitive to occasional out-of-place taxa than raw gene-tree support.

Because all RAxML analyses were conducted using the GTRCAT model, we also repeated the analyses of the CDS datasets using the GTRGAMMA model to investigate the robustness of the recovered topologies to slight model differences.

To summarize the overall bootstrap support of each tree with a single statistic, we calculated "percent resolution" as the number of bipartitions with >50% bootstrap support divided by the total number of bipartitions and represents the proportion of nodes that one might consider resolved (Kates et al. 2018). We visualized trees using FigTree (Rambaut 2016) and the APE package in R (Paradis et al. 2004). To compare trees, we used the phytools package in R (Revell 2012) to plot a consensus tree and to calculate a Robison-Foulds (RF) distance matrix for all trees. The RF distance between tree *A* and tree *B* equals the number of bipartitions unique to *A* plus the number of bipartitions unique to *B*. We visualized the first two principal components of the matrix using the Lattice package in R (Sarkar 2008). In addition, we conducted pairwise topology comparisons using the "phylo.diff" function from the Phangorn package in R (Schliep 2011) and an updated version of "cophylo" from phytools (github.com/liamrevell/phytools/). All statistical analyses took place in R (R Core Development Team, 2008).

Supermatrix analyses took place on the CIPRES Science Gateway (Miller et al. 2010). All other analyses took place on a computing cluster at the Chicago Botanic Garden, and almost all processes were run in parallel using GNU Parallel (Tange 2018). Alignments and trees have been deposited in the Dryad Data Repository (accession no. TBA).

### APPENDIX 2 – PARTITIONED EXON+NON-CODING DATASET

#### *Results*

The *partitioned supermatrix* analyses in which exons and introns were aligned separately were extremely divergent, particularly in the supermatrix analyses. The mean RF distance to any other tree for the *partitioned supercontig* trees was 138 (180 for the supermatrix trees and 96 for the ASTRAL trees). Likewise, the strict consensus of all 16 trees had only 48/159 nodes resolved. Re-running the supermatrix analysis with partitions by gene only did not improve concordance (mean RF 176) (Figure S8). The supermatrix trees for which introns and exons were aligned separately all contained a unique, taxonomically nonsensical clade of 38 samples, nested either within subgenus *Pseudojaca* or subgenus *Artocarpus*, characterized by increased missing data. A high proportion of missing "intron" sequences (>50%) appead to be the best predictor for membership in the nonsense clade; all members also had below-average "intron" sequence lengths, although sequence length seemed somewhat less correlated membership in the clade. Pruning tips with missing "intron" sequences dramatically reduced the divergence of the trees in question from other trees (Figure S9). The ASTRAL trees inferred from exons and introns aligned separately were not as severely divergent as the supematrix trees and did not contain the same nonsense clade (Figure S8).

#### *Discussion*

The divergent and questionable topologies of the supermatrix analyses for which introns and exons were aligned separately seems to relate at least in part to the alignment method, as removing the codon partitions did not improve the concordance of the supermatrix analysis. It is likely that intronic sequences, especially incomplete ones from samples with fewer or shorter reads, do not align well absent exons to anchor the alignments. Thus, even if missing data do not bias analyses per se (de la Torre-Bárcena et al., 2009), it may in effect create phylogenetically-misleading artifacts due to improper alignment or lack of sufficient characters (Rubin et al., 2012). However, partitioning the introns separately seems to have amplified this problem. Because RAxML does not provide for sub-partitions, each gene's first two codons, third codon, and introns were treated as independent, unlinked partitions in the supermatrix analyses. By contrast, the ASTRAL analyses of the same data set, which effectively had sub-partitions because each gene tree containing three partitions was estimated separately, were much less divergent, suggesting that overpartitioning can bias phylogenetic analyses.

Rubin, B.E.R., Ree, R.H., Moreau, C.S., 2012. Inferring phylogenies from RAD sequence data. PLoS One 7, e33394. doi:10.1371/journal.pone.0033394

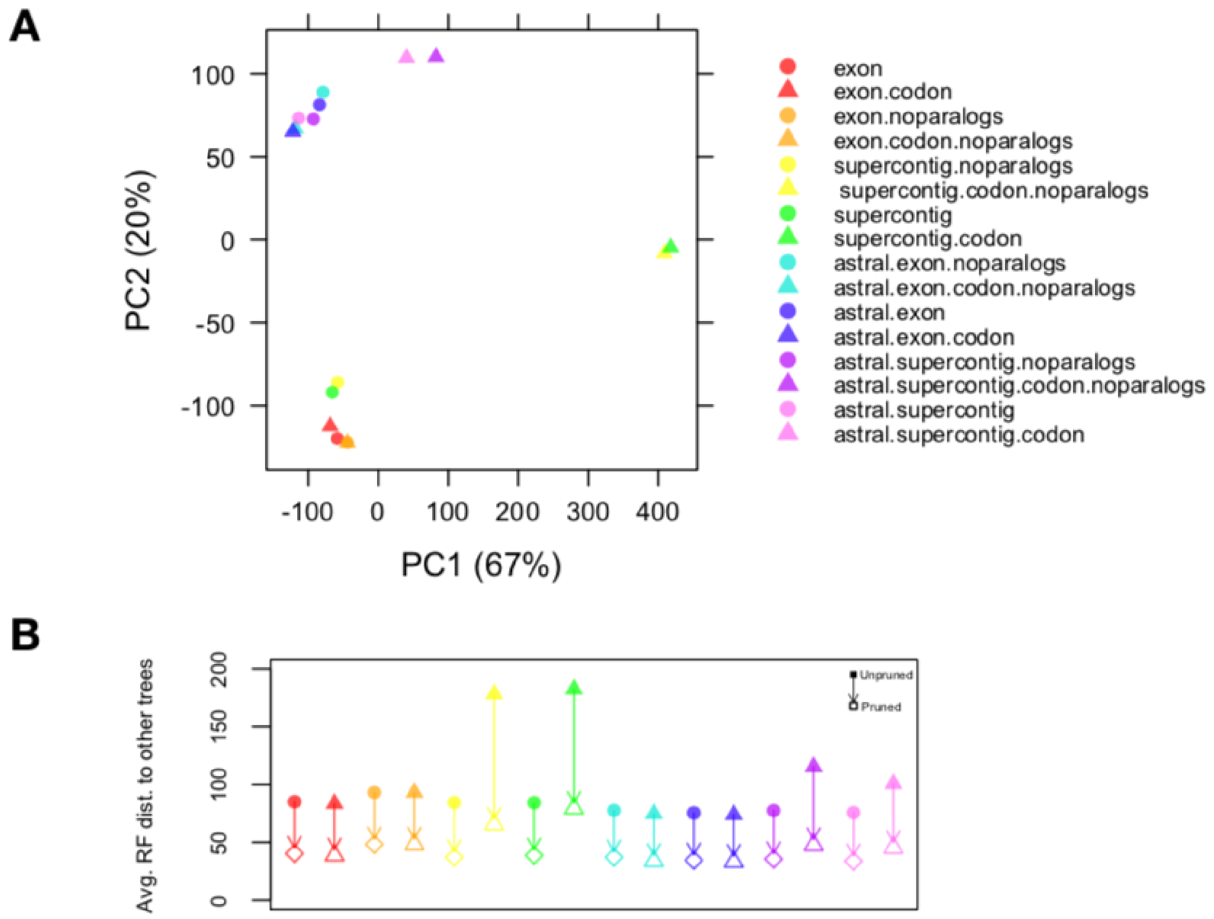


Figure S8. A) PCA of Robinson-Foulds (RF) distances between all 16 analyses showing extreme divergence in the supermatrix analysis of trees for which introns and exons were aligned separately. (B) Reductions in RF distances when 38 taxa with high (>50%) amounts of missing intron sequences were pruned from all trees.

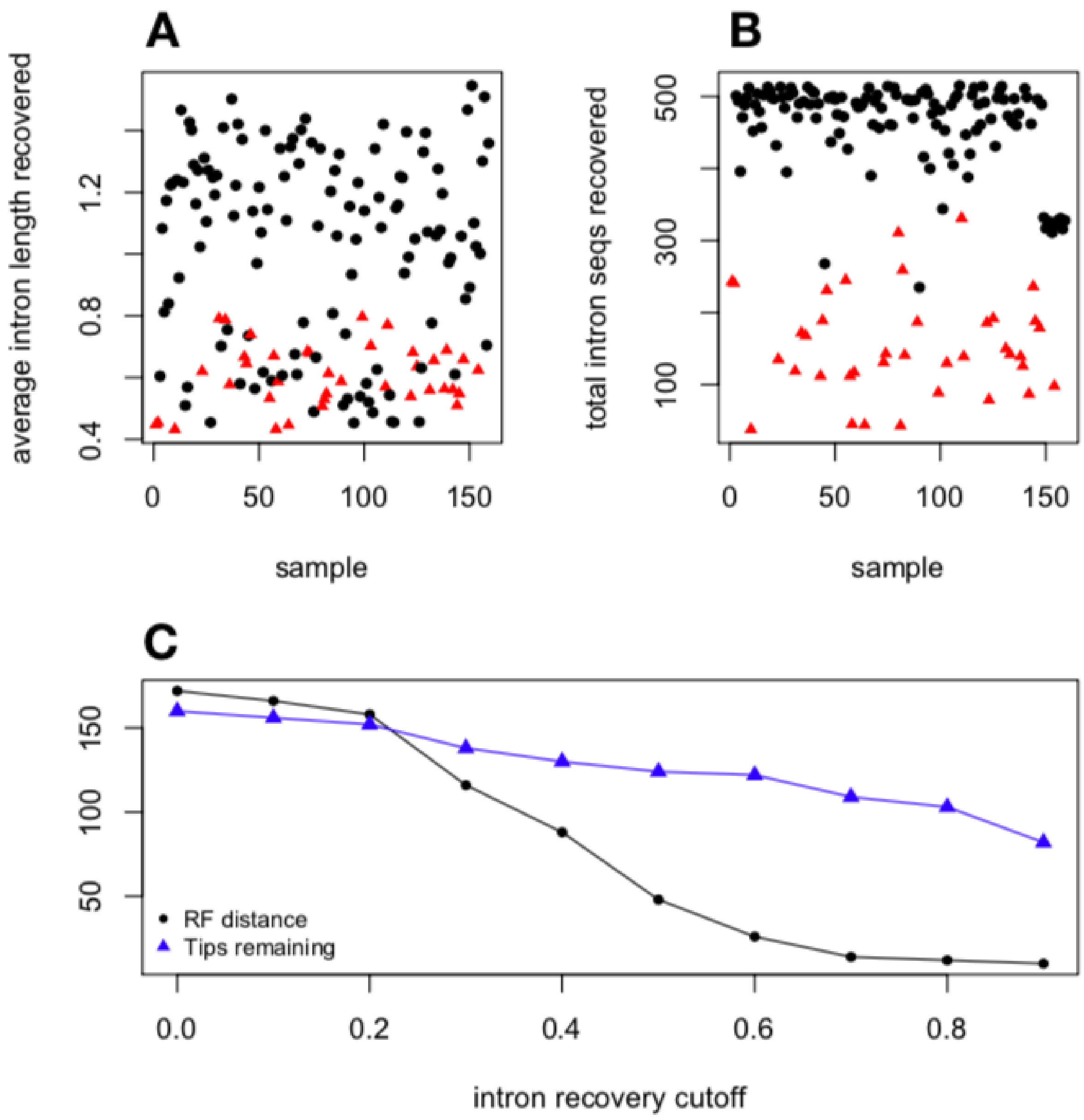


Figure S9. Investigation into the nonsense clade appearing in supermatrix analyses for which introns and exons were aligned separately. (A) Average intron length per sample after filtering (normalized within each gene as a percentage of mean length) and (B) total number of intron sequences present per sample after filtering, with samples in the nonsense clade as red triangles. (C) Comparison of two supermatrix trees, one based on supercontig alignments and one based on separate intron and exon alignments. Tips were pruned based on the proportion of genes with successfully-recovered non-coding sequences (x- axis). The y-axis displays the number of remaining tips in each tree (blue triangles) and the RF distance between the two trees (black circles).
